# Taxonomic classification of strain PO100/5 shows a broader geographic distribution and genetic markers of the recently described *Corynebacterium silvaticum*

**DOI:** 10.1101/2020.07.17.206599

**Authors:** Marcus Vinicius Canário Viana, Rodrigo Profeta, Alessandra Lima da Silva, Raquel Hurtado, Janaína Canário Cerqueira, Bruna Ferreira Sampaio Ribeiro, Marcelle Oliveira Almeida, Francielly Morais-Rodrigues, Manuela Oliveira, Luís Tavares, Henrique Figueiredo, Alice Rebecca Wattam, Debmalya Barh, Preetam Ghosh, Artur Silva, Vasco Azevedo

**Affiliations:** Department of Genetics, Ecology and Evolution, Institute of Biological Sciences, Federal University of Minas Gerais, Belo Horizonte, Minas Gerais, Brazil; Department of Genetics, Institute of Biological Sciences, Federal University of Pará, Belém, Pará, Brazil; Centre for Interdisciplinary Research in Animal Health, Faculty of Veterinary Medicine, University of Lisbon, Lisboa, Portugal; National Reference Laboratory of Aquatic Animal Disease, Federal University of Minas Gerais, Belo Horizonte, Minas Gerais, Brazil; Biocomplexity Institute, University of Virginia, Charlottesville, Virginia, USA; Institute of Integrative Omics and Applied Biotechnology, Purba Medinipur, West Bengal, India; Department of Computer Science, Virginia Commonwealth University, Richmond, Virginia, USA

**Author notes:** Corresponding author: Dr. Vasco Azevedo, (VA).

## Abstract

The bacterial strain PO100/5 was isolated from a skin abscess of a pig (*Sus scrofa domesticus*) in the Alentejo region of southern Portugal. It was identified as *Corynebacterium pseudotuberculosis* using biochemical tests, multiplex PCR and Pulsed Field Gel Electrophoresis. After genome sequencing and *rpoB* phylogeny, the strain was classified as *C. ulcerans*. To better understand the taxonomy of this strain and improve identification methods, we compared strain PO100/5 to other publicly available genomes from the *C. diphtheriae* group. Taxonomic analysis reclassified it and three others strains as belonging to the recently described *C. silvaticum*, which have been isolated from wild boar and roe deer in Germany and Austria. The results showed that PO100/5 is the first sequenced genome of a *C. silvaticum* strain from a domestic animal and a different geographical region, is a putative producer of the *diphtheriae* toxin, and has a unique sequence type. Genomic analysis of PO100/5 showed four prophages and eight conserved genomic islands when compared to *C. ulcerans*. Pangenome analysis of 38 *C. silvaticum* and 76 *C. ulcerans* samples suggest that *C. silvaticum* is a clonal species, with 73.6% of conserved genes and a pangenome near to being closed (α > 0.952). 172 conserved genes are unique to *C. silvaticum* when compared to *C. ulcerans*, with most related to nutrient uptake and metabolism, prophages or immune evasion. These unique genes could be used as genetic markers for species identification. This information can be useful for identification and surveillance of this pathogen, especially in regard to the possibility of zoonotic transmission.

## Introduction

The genus *Corynebacterium* from phylum Actinobacteria has Gram-positive bacteria of biotechnological, veterinary and medical relevance with free, commensal and pathogenic lifestyles. Within its pathogenic members, the most prominent species are the nearly exclusively human pathogen *C. diphtheriae* and the zoonotic species, *C. pseudotuberculosis* and *C. ulcerans*. These species compose the *C. diphtheriae* group, a clade of species that can be lysogenized by phages harboring the diphtheria toxin (DT) gene (*tox*) [1]. In this group, three new species were recently described. *C. belfantii* is a reclassification of *C. diphtheriae* biovar Belfanti [2] and a synonym of *C. diphtheriae* subspecies *lausannense* [3], *C. rouxii* [3] is reclassifications of *C. diphtheriae* biovar Belfanti, and *C. silvaticum* [4] is a reclassification of atypical *C. ulcerans* strains. Strains of *C. silvaticum* were previously described as atypical non-toxigenic but *tox*-gene-bearing (NTTB) strains of *C. ulcerans*, isolated from wild boars and roe deer in Germany, which caused caseous lymphadenitis similar to *C. pseudotuberculosis* infections [5–7]. This variant, examined using genomics and proteomics, was initially named as a “wild boar cluster” (WBC) of *C. ulcerans* [5–7] and later reclassified as *C. silvaticum* [4].

The strain PO100/5 was isolated from caseous lymphadenitis lesions in a Black Alentejano pig (*Sus scrofa domesticus*) from a swine farm in the Alentejo region of Portugal. It was identified as *Corynebacterium pseudotuberculosis* by both biochemical tests (Api Coryne® kit) and by multiplex PCR and Pulsed Field Gel Electrophoresis [8]. Genome sequencing and *rpoB* phylogeny showed that this strain was closer to *C. ulcerans* and the genome was deposited in GenBank as a strain from this species (accession number CP021417.1). During the submission and review of this manuscript, the description of *C. silvaticum* was published and suggested PO100/5 as a strain of this species by *rpoB* phylogeny [4], while a genomic analysis of 28 *C. ulcerans* strains suggested that PO100/5, W25 and KL1196 could represent a new species [9], although KL1196 had already been classified as *C. silvaticum* [4].

Pigs and boars are reservoirs of *C. silvaticum* [4–7] and *C. ulcerans* [10–12] and are known to transmit pathogens to humans and domestic animals [10,11,13]. Rapid, simple and reliable identification of species is essential for diagnosis, treatment and surveillance [14,15]. To better understand the taxonomy of PO100/5, the genome diversity of *C. silvaticum* and to identify molecular markers of this species, we performed a comparative analysis of 34 *C. silvaticum* and 80 *C. ulcerans* genomes, as well as other publicly available genomes from the *C. diphtheriae* group. We reclassified PO100/5 and other three strains recently deposited as *C. silvaticum* and found a unique sequence type and genes that can be useful for species classification.

## Materials and Methods

### Genomes, assembly and annotation

For the taxonomic, phylogenetic and genome plasticity analyses, a total of 120 genomes (S1 File) were selected, including 80 *C. ulcerans* and 34 *C. silvaticum* strains and six type strains of *C. diphtheriae* group. Assembled genomes were retrieved from Pathosystems Resource Integration Center (PATRIC) [16], while genomes available as sequencing reads were assembled in PATRIC using SPAdes [17] strategy. All genomes were annotated using the Rapid Annotation using Subsystems Technology (RASTtk) pipeline [18] that is available in PATRIC.

### Taxonomic analysis

Average Nucleotide Identity (ANI) was estimated using FastANI v1.3 [19]. An automatic genome-based taxonomic analysis was performed using Type (Strain) Genome Server (TYGS) (https://tygs.dsmz.de) [20]. First, TYGS pipeline identifies the closest type strains using MASH [21] for genomic sequences and BLAST [22] for 16S rRNA sequences. Second, it identifies the 10 closest type strains using Genome Blast Distance Phylogeny (GBDP) [23]. Third, it uses digital DNA:DNA hybridization (dDDH) [23] to cluster species and subspecies using a threshold of 70 and 79%, respectively [24].

Phylogenetic trees of *rpoB* and *tox* were built using the Maximum Likelihood method [25] implemented in MEGA v10.1.6 [26]. The *tox* tree included all sequences in genomes of *C. ulcerans* and outgroups from *C. silvaticum, C. pseudotuberculosis* and *C. belfantii. C. rouxii* was not included due to all sequenced genomes being *tox-* [3]. All trees were visualized using iTOL [27].

Multi Locus Sequence Typing (MLST) was performed using MLSTcheck [28], using the scheme for *C. diphtheriae* and *C. ulcerans* (genes *atpA, dnaE, dnaK, fusA, leuA, odhA* and *rpoB*) [12]. The Minimmum Spanning Tree (MST) generated using goeBURST algorithm was built using PHYLOViZ [29].

### Genome plasticity analysis

Prophages of PO100/5 were predicted using PHASTER [30]. Genomic islands were predicted using GIPSy v1.1.2 [31], using *C. ulcerans* NCTC7910^T^ and *C. pseudotuberculosis* ATCC19410^T^ as references. A circular map was generated using BRIG v0.95 [32]. The presence of true or candidate virulence factors of *Corynebacterium* [33,34] were verified using PATRIC’s Protein Family Sorter tool, Proteome Comparison tool, and by comparing the gene neighborhood with other strains using Artemis Comparison Tool 17.0.1 [35]. Signal peptide and conserved protein domains were verified using InterProScan [36], while cell wall sorting signal (CWSS) was verified using CW-PRED [37]. The identification of homologous genes (orthogroups) was performed using OrthoFinder v2.3.12 [38]. In house scripts were used to extract all orthogroups (pangenome) and subsets representing orthogroups conserved across all genomes (core genome), orthogroups shared by more than one but not all genomes (shared genome), orthogroups exclusive from a genome (singletons), orthogroups shared by more than one but not all genomes or exclusive of a genome (accessory genome) [39] and orthogroups conserved and exclusive to a group of genomes (exclusive core); and to calculate the development of pangenome, core genome and singletons [39]. The development of the pangenome was calculated according to Heaps’ law fit formula *n* = κ\**N*^γ^, in which *n* is the number of genes, *N* is the number of genomes, and *κ* and γ (α = γ -1) are free parameters determined empirically. The pangenome is estimated as closed when α > 1 (γ < 0), which means no significant increase with the addition of new genomes. The pangenome is estimated to be open when α ≤ 1 (0 < γ < 1). The development of core genome and singletons were calculated using least-squares fit of the exponential regression decay n = κ*exp[-N/τ] + tg(θ), in which *n* is the number of genes, *N* is the number of genomes, and κ, τ, and tg(θ) are free parameters determined empirically [39]. A functional annotation of genes was performed using eggNOG-mapper v2 [40].

## Results

### Taxonomic analysis

ANI results showed that *C. ulcerans* strains PO100/5, 04-13, 05-13 and W25 had identity values ≥ 99.74% with *C. silvaticum* KL0182^T^ and ≤ 91.03% with *C. ulcerans* NCTC7910^T^ (S2 File). The taxonomic classification using TYGS classified those strains as *C. silvaticum*, due to genome and 16S rRNA GBDP trees, dDDh > 70% in the formula *d4* and GC content difference > 1% with *C. ulcerans* genomes (S3 File). In the *rpoB* phylogeny, the *C. ulcerans* strains PO100/5, 04-13, 05-13 and W25 clustered with *C. silvaticum* KL0182^T^, while other *C. ulcerans* strains formed two clades (Fig 1). In the phylogenetic tree of *tox* gene, the same four strains (PO100/5, 04-13, 05-13 and W25) also cluster with *C. silvaticum*, separated from *C. ulcerans, C. diphtheriae* and *C. pseudotuberculosis* clusters (Fig 2). MLST analysis classified strains 04-13, 05-13 and W25 as ST578 (53-60-121-70-76-66-57) and PO100/5 as a new ST, differing from ST578 by a new allele in locus *odhA* (66∼) (S4 File, Fig 3). Due to those results those four strains were reclassified for the next analyses, changing the number of *C. silvaticum* genomes from 34 to 38 and *C. ulcerans* genomes from 80 to 76.

**Fig 1.**
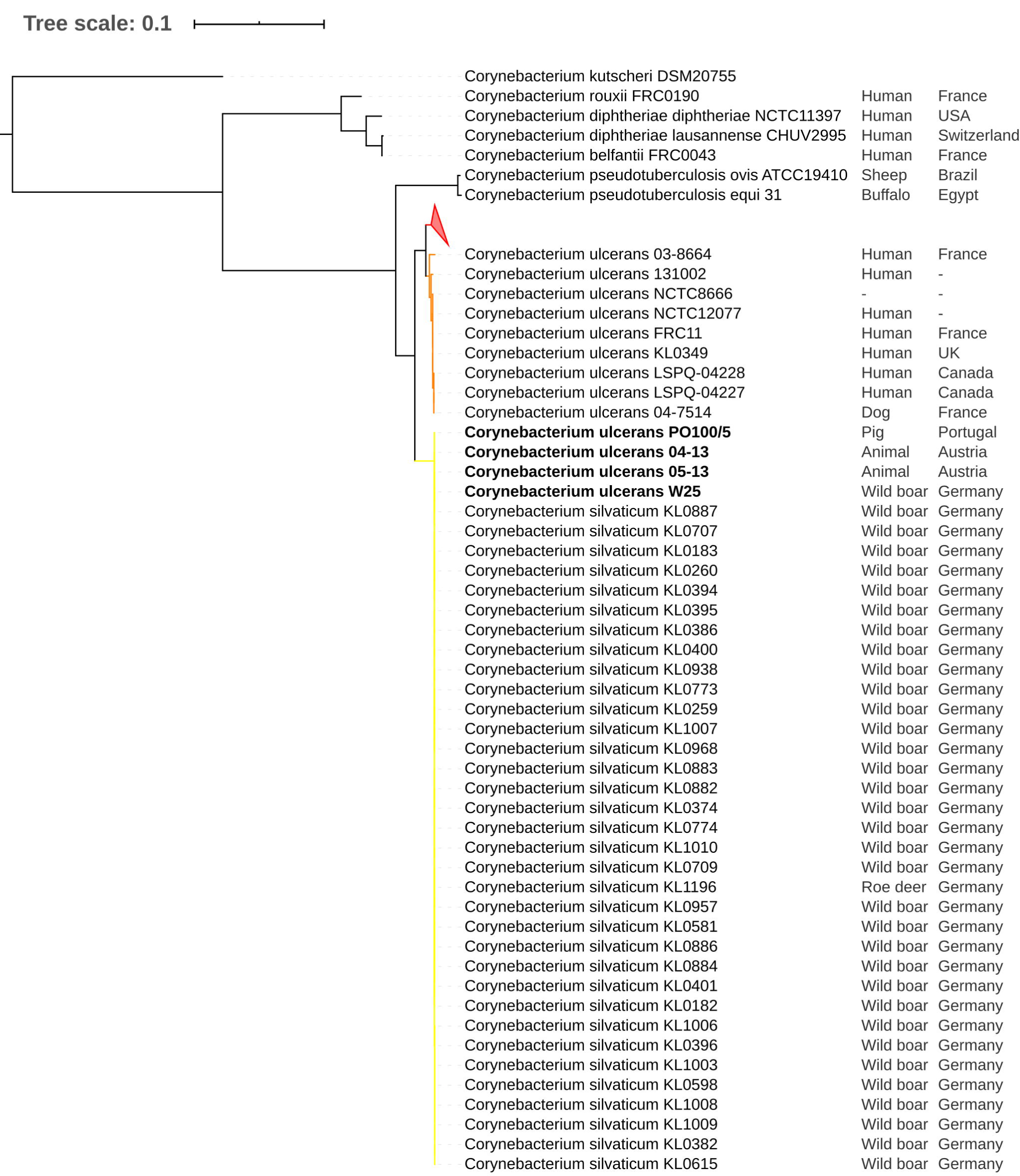
Phylogenetic tree of *rpoB* gene from *Corynebacterium* species. The phylogeny was inferred using the Maximum Likelihood method and the Tamura-Nei (TN93 + G) model implemented in the Mega v10.1.6. The *Corynebacterium ulcerans* strains PO100/5, 04-13, W25 and 05-13 cluster with the *Corynebacterium silvaticum* KL0182^T^ (yellow). *C. ulcerans* strains form lineage 1 (red, colapsed) and lineage 2 (orange).

**Fig 2.**
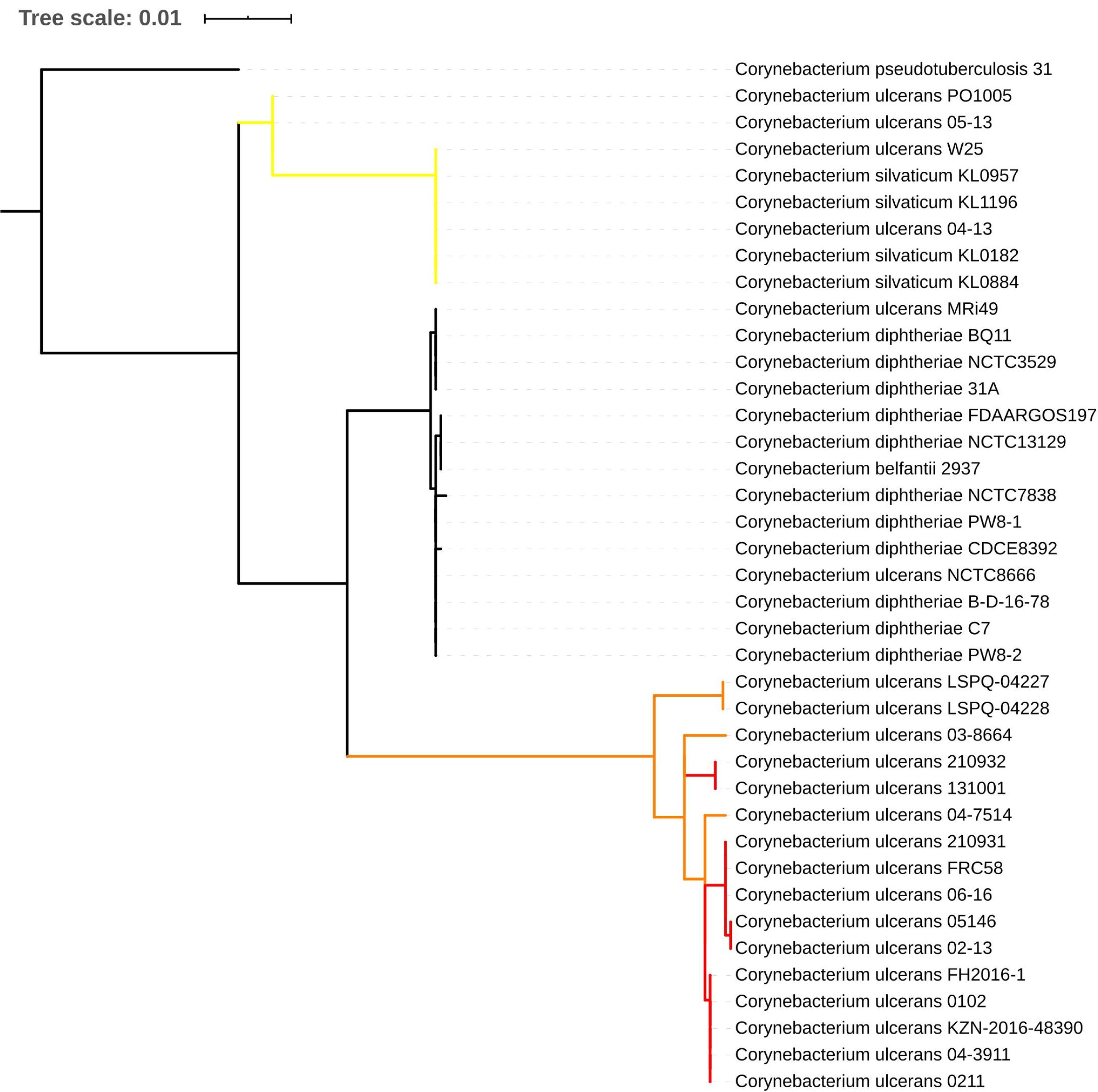
Phylogenetic tree of the *tox* gene from *Corynebacterium* species. The phylogeny was inferred using the Maximum Likelihood method and the Tamura-Nei (T92 + G) model implemented in Mega v10.1.6. The strains PO100/5, 04-13, W25 and 05-13 cluster with *Corynebacterium silvaticum*. Clade colors represent *rpoB* clades of *C. silvaticum* (yellow) and *C. ulcerans* (red and orange).

**Fig 3.**
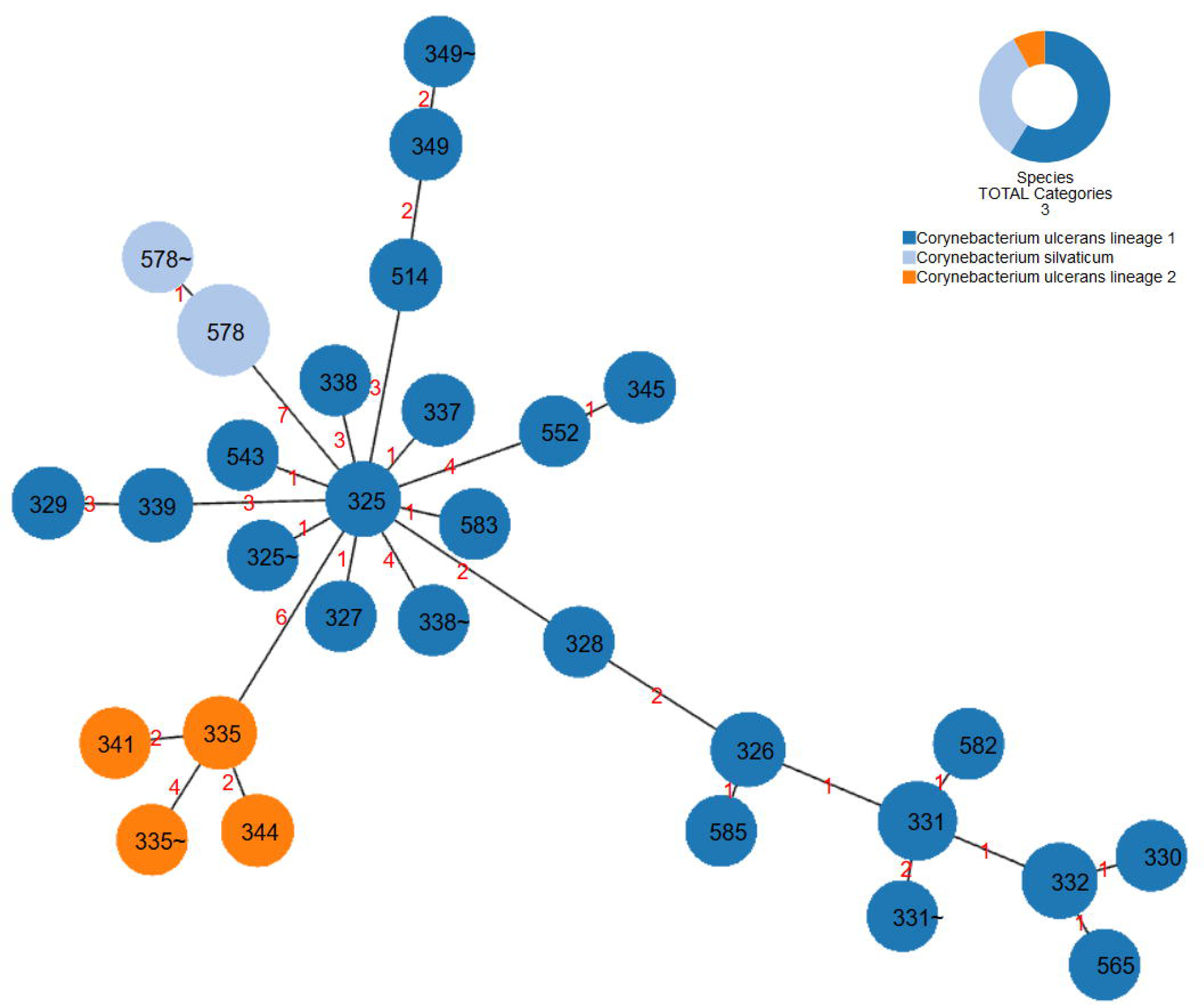
goeBURST diagram for the MLST data set of 38 *Corynebacterium silvaticum* and 76 *C. ulcerans* strains generated using PHILOViZ. The red numbers on the links indicate the number of divergent alleles between STs.

The taxonomic analysis led to additional insights. TYGS classified nine *C. ulcerans* strains as a potential new species: 03-8664, 04-7514, 131002, FRC11, KL0349, LSPQ-04227, LSPQ-04228, NCTC8666 and NCTC12077. These genomes had dDDH values greater than 70% (99.8 – 75.9%) within them and less than 70% (63 to 67.2%) with other *C. ulcerans* genomes, although the GC content difference was less than 1% (S3 File). In ANI analysis, those nine genomes were more similar to each other than to the other genomes. They had values between 95.52 and 96.57% with *C. ulcerans* NCTC7910^T^, and ≥ 97.82% when one of them (NCTC12077) was used as reference for the other eight (S2 File). MLST analysis classified them as having the unique sequence types ST345, ST341, ST344 and a new ST derived from ST345 (S4 File). The ANI analysis showed 99.3% identity between *C. diphtheriae lausannense* strains CHUV2995 and *C. belfantii* FRC0043^T^ (S2 File). A further analysis using TYGS classified *C. diphtheriae lausannense* strains CHUV2995^T^ and CMCNS703 as *C. belfantii*, and the nine *C. belfantii* genomes, besides the type strain FRC0043^T^, as *C. diphtheriae* (S3 File).

### Genome plasticity analysis

The presence of genes encoding 16 known true or candidate virulence factors of *Corynebacterium* [33,34] in *C. silvaticum* is shown in Table 1. The genes *rhuM, rpb* and *tspA* are absent, while all the pilus genes except *spaB* are pseudogenized, lacking the signal peptide or CWSS. *C. silvaticum* has the two pilus gene clusters structured as *srtA, spaB*C, and *srtB, spaD, srtC* and *spaEF*, despite fragmentation of pilin genes. Only eight genomes had the *tox* gene (04-13, 05-13, KL0182, KL0884, KL0957, KL1196, PO100/5 and W25), and only PO100/5 does not have a two bases insertion (GG) in position 48 that introduces a frameshift. In *C. ulcerans, tspA* is present in all strains, *rpb* is present only in strain 809, and *rhuM* is present in 16 strains from Austria, France and Germany isolated from humans, cats and dogs (02-13, FRC58, KL0195, KL0246-cb3, KL0251-cb4, KL0252-cb5, KL0349, KL0387-cb8, KL0475, KL0497, KL0541, KL0547, KL0796, KL0867, KL0880, NCTC12077).

**Table 1.**
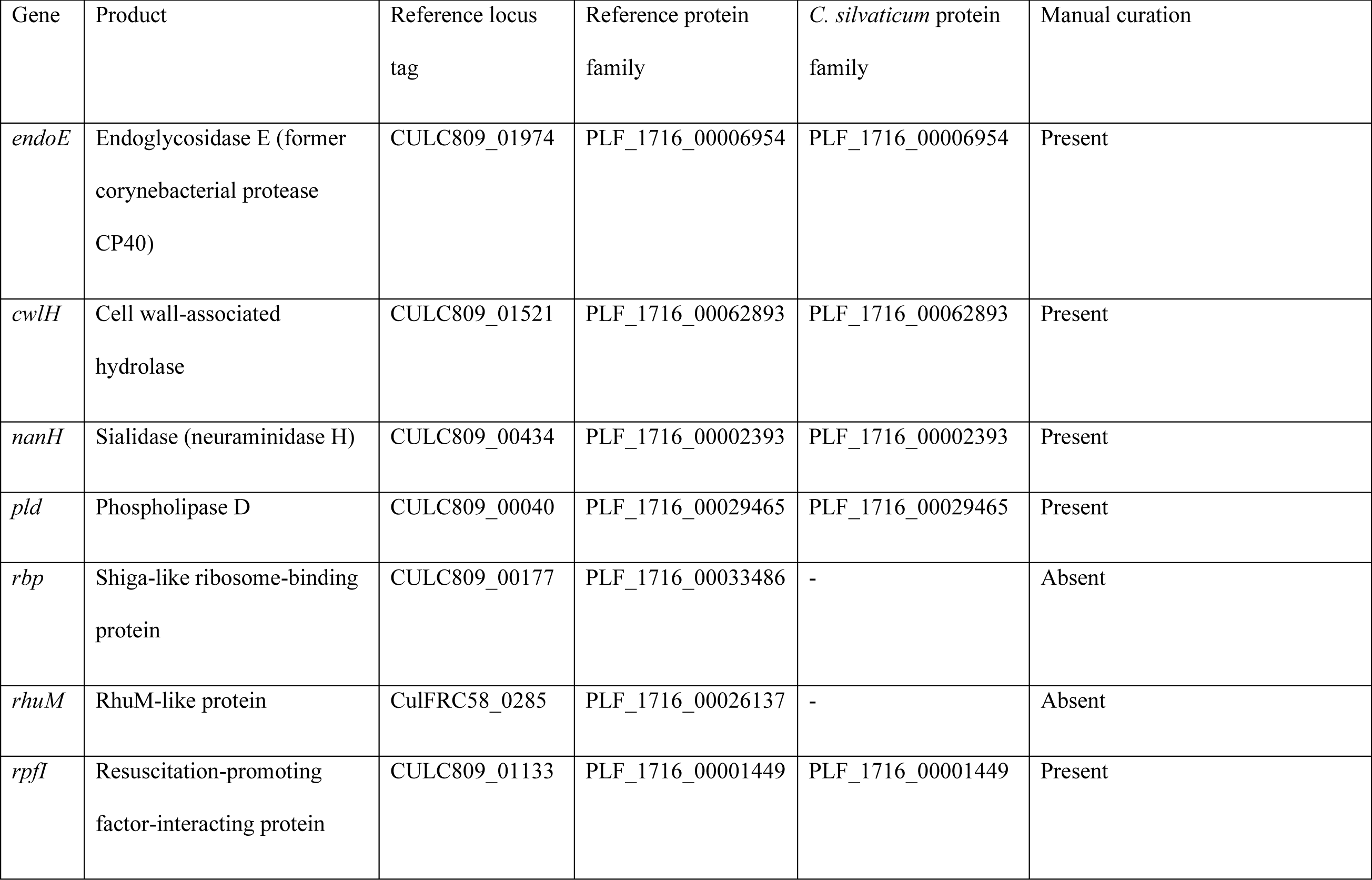
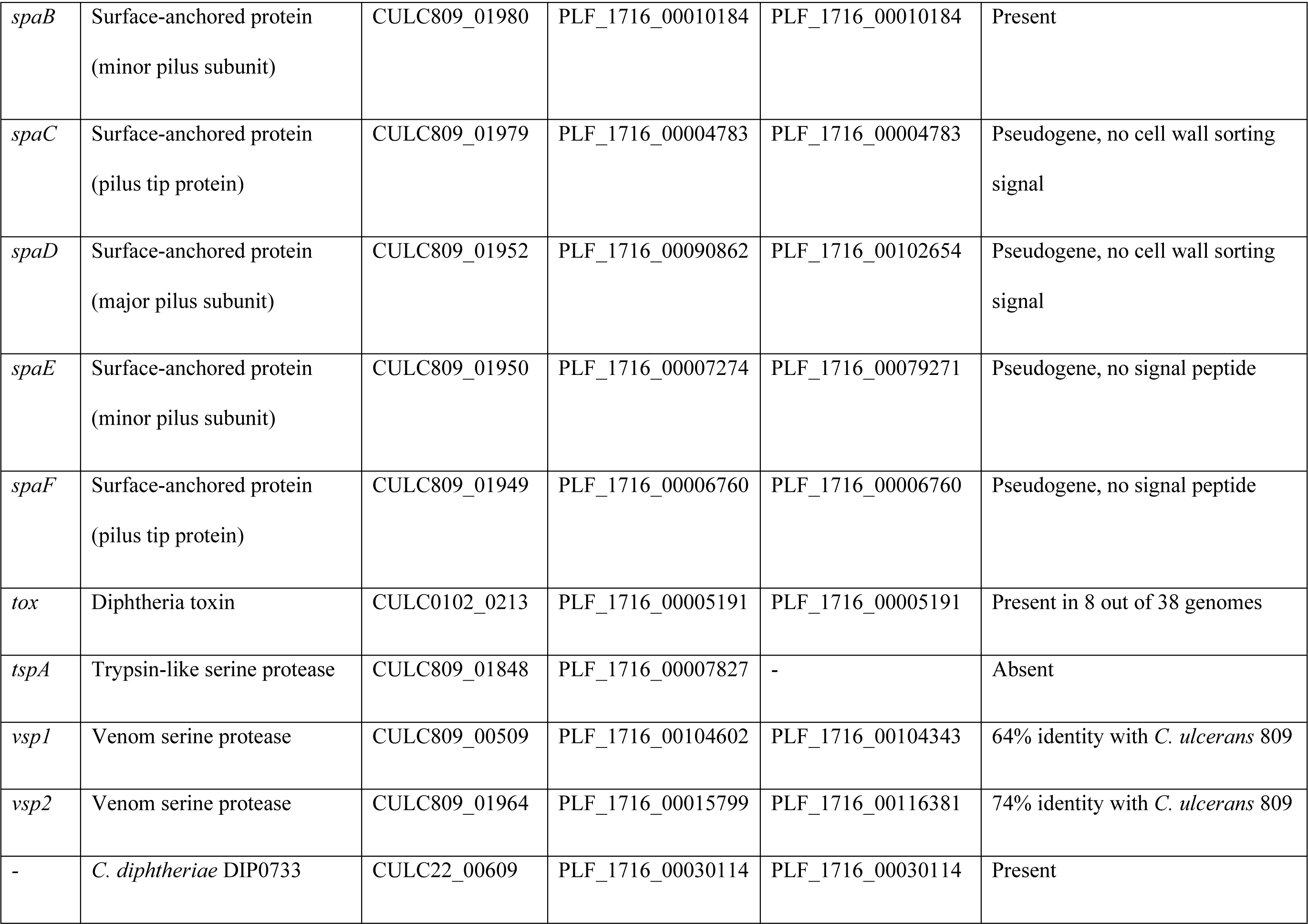
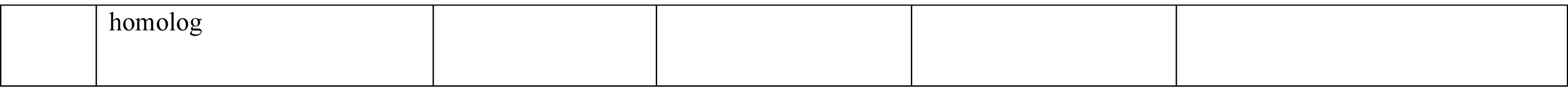
Presence of 16 known true or candidate virulence factors of *Corynebacterium* in *C. silvaticum*.

Sixteen and eight islands were predicted by comparing PO100/5 with the reference strains *C. pseudotuberculosis* ATCC19410^T^ and *C. ulcerans* NCTC7910^T^, respectively, and no island was detected in comparison to *C. silvaticum* KL0182^T^ (Table 2, Fig 4). The content of the islands is described in S5 File. Within them, four incomplete prophages were predicted, one of them harboring the *tox* gene (Table 3, Fig 4). The number of orthogroups in pangenome, core genome, accessory genome and singletons of *C. silvaticum* and *C. ulcerans* is shown in Table 4 and S6 File. The core genome represented 73.6 and 40% for *C. silvaticum* and *C. ulcerans*, respectively. The pangenome, core genome and singletons development graphs and formulas are shown in Fig 5.

**Table 2.** Genomic islands in strain PO100/5 in comparison to *Corynebacterium pseudotuberculosis* ATCC19410^T^ and *C. ulcerans* NCTC7910^T^.

| n | Position compared to Cp | Size | Position compared to Cul | Size | Type | Prophage content |
| --- | --- | --- | --- | --- | --- | --- |
| 1 | 29362-38109 | 8.75 kb | - | - | - | - |
| 2 | 55224-60448 | 5.22 kb | 55224-60448 | 5.22 kb | PA | - |
| 3 | 69256-74863 | 5.6 kb | - | - | PA, RE, SY | - |
| 4 | 97842-103378 | 5.53 kb | - | - | PA, RE | - |
| 5 | 167859-206438 | 38.58 kb | 167859-206209 | 38.35 kb | - | Prophage I |
| 6 | 311533-318567 | 7.03 kb | 311533-318567 | 70.34 kb | - | - |
| 7 | 422293-430839 | 85.46 kb | - | - | RE, SY | - |
| 8 | 694806-746966 | 52.16 kb | 694806-746966 | 52.16 kb | RE | Prophage II |
| 9 | 925047-938225 | 13.18 kb | 925047-938225 | 13.18 kb | - | - |
| 10 | 1235769-1244625 | 8.86 kb | 1235769-1244625 | 8.86 kb | - | - |
| 11 | 1613625-1644953 | 31.33 kb | 1614013-1639302 | 25.29 kb | - | Prophage III |
| 12 | 1793740-1802246 | 8.5 kb | - | - | PA, ME | - |
| 13 | 2029016-2035120 | 6.1 kb | - | - | - | - |
| 14 | 2109299-2139505 | 30.2 kb | 2110317-2139505 | 29.19 kb | - | Prophage IV |
| 15 | 2255307-2261370 | 6.06 kb | - | - | - | - |
| 16 | 2517632-2529863 | 12.23 kb | - | - | ME | - |

Cp – *C. pseudotuberculosis*, Cul – *C. ulcerans*, PA – pathogenicity island, RE – resistance island, ME – metabolic island, SY – symbiotic island.

**Table 3.** Incomplete prophages in strain PO100/5.

| Prophage | Length | Position |
| --- | --- | --- |
| Prophage I (*tox*<sup>+</sup>) | 44.5 kb | 161483-206048 |
| Prophage II | 23.5 kb | 719365-742927 |
| Prophage III | 44.3 kb | 1594987-1639315 |
| Prophage IV | 9.9 kb | 2129289-2139193 |

**Table 4.** Pangenomics of *Corynebacterium silvaticum* and *C. ulcerans*.

| Species | Genomes | Pangenome | Core genome | Shared genome | Singletons | $\alpha$ |
| --- | --- | --- | --- | --- | --- | --- |
| *C. silvaticum* | 38 | 3,002 | 2,209 | 603 | 190 | 0.9520 |
| *C. ulcerans* | 76 | 4,351 | 1,747 | 1,706 | 898 | 0.8142 |
| *C. silvaticum*<br>and *C. ulcerans* | 114 | 4,916 | 1,618 | 2,349 | 949 | — |

**Fig 4.**
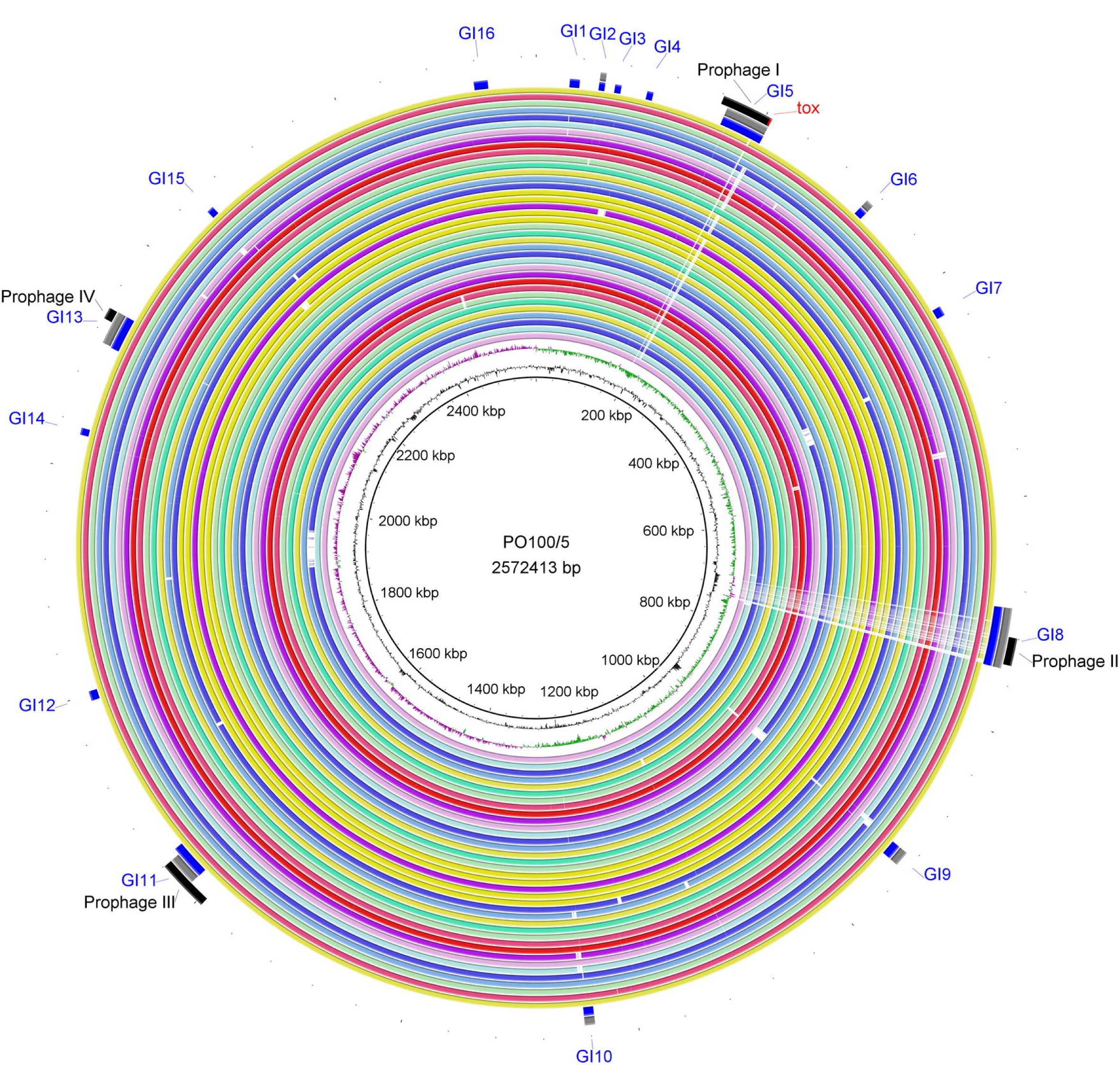
Circular map of *Corynebacterium silvaticum* genomes generated using BRIG v0.95. From inner to outer circle: strain PO100/5 (reference); CG content; CG Skew; strains KL0182^T^, KL0183, KL0259, KL0260, KL0374, KL0382, KL0386, KL0394, KL0395, KL0396, KL400, KL0401, KL0581, KL0598, KL0615, KL0707, KL0709, KL0773, KL0774, KL0882, KL0883, KL0884, KL0886, KL0887, KL0938, KL0957, KL0968, KL1003, KL1006, KL1007, KL1008, KL1009, KL1010, KL1196, 05-13, 04-13, W25; genomic islands compared to *C. pseudotuberculosis* ATCC19410^T^ (blue) and *C. ulcerans* strain NCTC7910^T^ (grey); and prophages (black). Genomic islands (GI) and prophage detection were performed using GIPSy and PHASTER, respectively.

**Fig 5.**
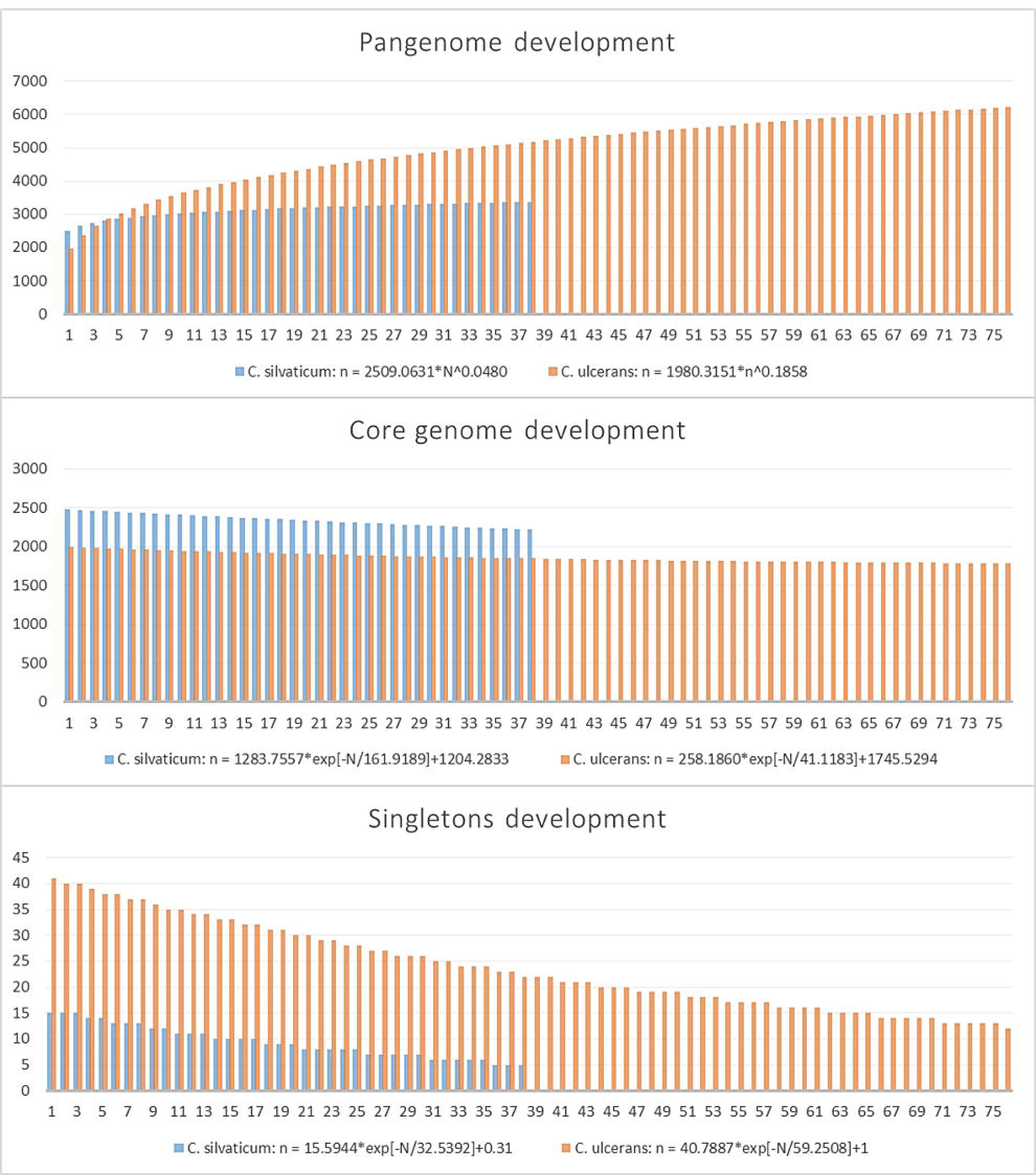
Pangenome, core genome and singletons development graphs and formulas for 38 genomes of *Corynebacterium silvaticum* and 76 *C. ulcerans* genomes.

Both species had genes conserved in all strains that were absent in the other species, or the exclusive core. In *C. silvaticum*, 172 orthogroups were detected in this subset. They are represented in strain PO100/5 by 174 proteins, 81 of them located across genomic islands 1, 2, 5, 6, 8, 9, 10, 11, 12 and 14. *C. ulcerans* lineage 2 had a hypothetical protein with 37 amino acids (S6 File). A graph comparing the distribution of Cluster of Homologous Groups (COG) categories of the exclusive core genome of both species is shown in Fig 6.

**Fig 6.**
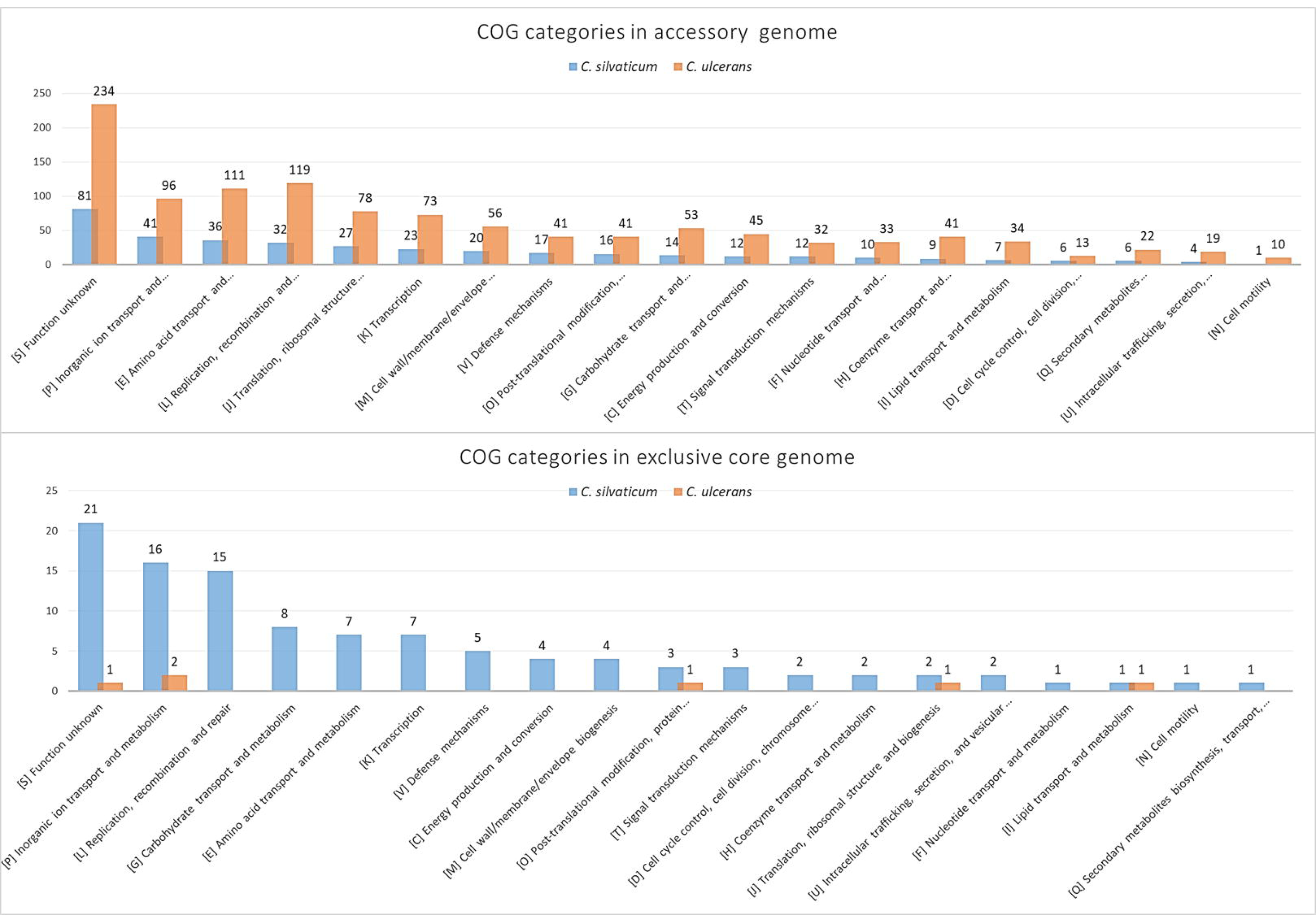
Clusters of Orthologous Groups (COGs) in the accessory and exclusive core genomes of *Corynebacteirum silvaticum* and *C. ulcerans* annotated using eggNOG-mapper v2. COG categories are sorted from most abundant to less abundant in *C. silvaticum*. In *C. silvaticum*, 356 out of 793 proteins from the accessory genome and 93 out of 174 proteins of the exclusive core genome had a COG category. In *C. ulcerans*, 1,065 out of 2,064 proteins from the accessory genome and six out of nine proteins of the exclusive core genome had a COG category

## Discussion

Strain PO100/5 was isolated from a caseous lymphadenitis lesion of a pig from a swine farm in the Alentejo region of Portugal. It was classified as *C. pseudotuberculosis* by biochemical tests (Api Coryne® kit), multiplex PCR and Pulsed Field Gel Electrophoresis [8]. After genome sequencing, a *rpoB* phylogeny clustered it close to *C. ulcerans* and the genome was deposited in GenBank as this species in 2017 (accession number CP021417.1). During the submission and review of this manuscript, PO100/5 was suggested as *C. silvaticum* by *rpoB* phylogeny, while another study of 28 *C. ulcerans* genomes suggested strains PO100/5, W25 and KL1196 [4] as a new species [9]. Here, we confirmed the taxonomy of PO100/5 as *C. silvaticum* and analyzed the genome diversity of this species, using 34 *C. silvaticum* and 80 *C. ulcerans* genomes, four of the *C. ulcerans* reclassified, as well as other publicly available genomes from the *C. diphtheriae* group (S1 File).

Taxonomic analysis showed that PO100/5, W25, 04-13 and 05-13 are strains that belong to the recently described *C. silvaticum* [4]. This is supported by ANI values above 95% [19] (S2 File), genome and 16S rRNA GBDP clustering [23], dDDH > 70% in the formula *d4*, GC content difference > 1% with *C. ulcerans* genomes [20,23,41] (S3 File), *rpoB* phylogenetic clustering (Fig 1) and the unique sequence type ST578 from *C. silvaticum* [4] (Fig 3). Strain PO100/5 has a new allele in locus *odhA* (S4 File, Fig 4), which results in a new ST derived from ST578. The original misclassification of those strains was understandable, as prior to the development of methods to identify *C. silvaticum* [4,5], the use of biochemistry tests (API Coryne and VITEK2-compact) and the clinical picture would classify this strains as *C. pseudotuberculosis* [4,42], while DNA sequence analysis and fourier-transformed infrared spectroscopy would classify it as *C. ulcerans* [6,7,42].

Analysis of genome plasticity identified unique characteristics of *C. silvaticum*. The analysis of 16 known true or candidate virulence factors showed the absence of *rpb, rhuM, and tsA*, and *spaB* as the only non-fragmented pili gene in *C. silvaticum* (Table 1). The Shiga-like ribosome-binding protein (*rpb*) has a ribosome inactivating protein domain and was reported only in *C. ulcerans* 809 [33,43]. The RhuM-like protein (*rhuM*) was reported only in *C. ulcerans* strain KL0387 from a human, probably transmitted by a cat [44]. A RhuM mutant of *Salmonella enterica* had a significant decrease in epithelial cell invasion [45]. Here, we identified this protein in 15 other strains from humans, dogs and cats form Austria, France and Germany. Serine proteases can promote the survival and dissemination of pathogens in the host [46]. Venom serine proteases (*vsp1* and *vsp2*) and Trypsin-like serine protease (*tspA*) are secreted proteases that could have multiple potential pathogenic functions [47]. The effect of the absence of *tspA* in *C. silvaticum* is unknown, but it can be used as a marker to differentiate it from *C. ulcerans*. Bacterial pili are adhesion structures required for colonization of host tissues. The *Corynebacterium* pili are SpaA-type, a heterotrimeric structure composed by major (pilus shaft), minor and tip pilins, the last two required for adhesion. The pilus is assembled and anchored to the cell wall by the housekeeping sortase SrtA and pili sortases SrtB and SrtC [48]. As with *C. ulcerans* [33], *C. silvaticum* has the two pili gene clusters *spaB*C and *spaDEF*, although only *spaB* appears to be functional due to the presence of a signal peptide and a CWSS. The SpaB is a minor pilin that in *C. diphtheriae* has a role in adhesion on pharyngeal epithelial cells and could be functional when linked to the cell wall [49] as shown for the heterodimeric structure SpaB-SpaC in *C. diphtheriae* [50] and suggested for *C. ulcerans* [33]. Intriguingly, only eight of the 38 genomes had the *tox* gene (04-13, 05-13, KL0182, KL0884, KL0957, KL1196, PO100/5 and W25), although the strains lacking it was reported to be *tox*^*+*^ [6]. The absence of *tox* could be the result of an assembly artifact, due to a repetitive region prior to this gene. This can be seen in the circular map as a blank space in the *tox* gene region of the other strains (Fig 3). In a previous study, the *tox* gene from *C. silvaticum* strains W25 and KL1196 was shown to have an insertion of two bases (GG) in position 48, turning it into a pseudogene, while PO100/5 did not has this insertion [9]. This suggest PO100/5 could be a toxigenic *C. silvaticum*.

In PO100/5, four incomplete prophages (Table 2, Fig 3) were found, one harboring the *tox* gene. Sixteen and eight genomic islands were found when compared to *C. pseudotuberculosis* ATCC 19410^T^ and *C. ulcerans* NCTC7910^T^ (Table 3, S5 File, Fig 3), with four of them containing the incomplete prophages. No island was found when compared to *C. silvaticum* KL0182^T^. Genomic islands are regions acquired by horizontal gene transfer that can provide adaptive traits [31]. In previous studies, the phylogenetic analysis of prophages and other mobile elements, *tox* and DtxR gene sequences showed species-specific clades, including the atypical *C. ulcerans* clade that now represents *C. silvaticum* [6]. This implies different or independent events of acquisition of virulence factors that could be related to different disease manifestations, progression, as well as an expanded host reservoir that can result in zoonotic transmission [6,43].

*C. silvaticum* was estimated to be a more clonal species than *C. ulcerans* and to have a pangenome near to being closed, and bigger core genome, with higher values of core genome development (Fig 5) and α closer to 1 (Table 4). This result could be influenced by the samples of *C. silvaticum* being from two countries, Germany (n = 37) and Portugal (n = 1), and two species of host (*Sus scrofa* and *Capreolus capreolus*). A total of 172 and 8 orthogroups were uniquely shared by all *C. silvaticum* and *C. ulcerans*, respectively, some in genomic islands (S6 File, Fig 6). For *C. silvaticum*, the most abundant functions are involved in nutrient acquisition such as transport and metabolism of inorganic ions, carbohydrates and amino acids (COG categories E, G and P), or are related to phages or immunity against them (COG category L). For example, two of them are a Type I restriction-modification system [51] in genomic island 11 and an “ABC-type dipeptide oligopeptide nickel transport system”. The function of those genes in the phenotype and infection must be investigated, but they are candidates for genetic markers for a rapid and cost-effective diagnostic using multiplex polymerase chain reaction (PCR) [52–54], along with the MLST analysis, including the new ST of PO100/5, and other established methods [4,5].

Additionally, the TYGS results suggest that the nine *C. ulcerans* corresponding to lineage 2 [43] represent a potential new species, with dDDH of less than 70% with lineage 1 genomes. MLST classified these genomes with the unique sequence types ST345, ST341, ST344 and a new one derived from ST345 (S3 and S4 Files). In a previous analysis, ST335 and ST344 were the only known STs found in lineage 2 [43]. Further investigation is required to verify whether this lineage could be classified as a new species. Recently, *C. belfantii* and *C. diphtheriae lausannense* were suggested as synonyms [3]. Our analysis using TYGS corroborated the suggestion and showed that other genomes of *C. belfantii* besides the type strain (FRC0043^T^) are *C. diphtheriae* (S3 File), showing the difficulty in classifying this recently described species [2].

We showed that strains PO100/5 (domestic pig), W25 (wild boar), 04-13 and 05-13 are *C. silvaticum*. This species was previously described as atypical NTTB *C. ulcerans* or “wild boar cluster” (WBC) that infects wild boars and roe deer in Germany and Austria and causes lymphadenomegaly similar to *C. pseudotuberculosis* infections [5–7]. As other strains of this group are found in Germany and Austria [5,7], PO100/5 is the first genome of this species isolated from a different geographical area (Portugal) and from a domesticated animal (farm pig) [8] and to have a unique ST. The production on DT was not tested in strain PO100/5 [8], but the absence of a two bases insertion in the *tox* gene, that occurs in the other strains, suggest that it may be the first toxigenic strain described for this species [6,7].

In addition to the veterinary concerns posed by this pathogen, *C. silvaticum* could also have a medical relevance. Its known host range is limited to wild boars, domestic pig and roe deer [4,7,8]. Wild boars are reservoirs for viruses, bacteria and other parasites that can be transmitted to domestic animals and humans, during opportunities provided by deforestation and use of lands for agricultural purposes, hunting activities and consumption of wild boar meat [13]. Besides other hosts, pigs and boars are a reservoir of *C. ulcerans* that can cause zoonotic transmission to humans [10–12]. By the same route, *C. silvaticum* could be transmitted to humans and cause infection. In addition, it could be misidentified as *C. ulcerans* or *C. pseudotuberculosis* due to limitations of the standard methods [4,5]. The data generated in this study along with the available literature can be used in the identification, treatment and surveillance of this pathogen.

## Conclusions

The taxonomic analysis shows PO100/5 and four other genomes deposited as *C. ulcerans* should be included in the recently described species *C. silvaticum*. The comparative genomic analysis showed this species is more clonal than *C. ulcerans*, has SpaB as the only probably functional pilin subunit, and has conserved genomic islands and 172 genes that could be used as molecular makers for PCR identification. In contrast to the other strains from the same species, PO100/5 is the first one to be isolated from a domestic animal, the first isolation from Germany, and also has a unique ST and a non-pseudogenized *tox* gene. This data can be used in the identification, treatment and surveillance of this pathogen.

## Supporting information

**S1 File. Genomes of *Corynebacterium* species used for taxonomic analysis of the strain PO100/5.**

**S2 File. Average Nucleotide Identity and digital DNA-DNA hybridization among strains of *Corynebacterium*.**

**S3 File. Taxonomic classification of *Corynebacterium ulcerans* and *C. silvaticum*.** The analysis was performed using Type Strain Genome Server.

**S4 File. Multilocus Sequence Typing data of *Corynebacterium silvaticum* and *C. ulcerans* genomes.** The analysis was performed using MLSTchecker.

**S5 File. Genomic islands content in strain PO100/5.**

**S6 File. Pangenome analysis of 38 *Corynebacterium silvaticum* and 76 *C. ulcerans* samples.** Gene homology groups were predicted using OrthoFinder v2.12.2 and functional annotation was performed using eggNOG-mapper v2.

