## Supplementary File 3 for "Taxonomic classification of strain PO100/5 shows a broader geographic distribution and genetic markers of the recently described *Corynebacterium silvaticum*": Cb_Cd_lausannense.pdf

### TYPE (STRAIN) GENOME SERVER

### JOB SUMMARY

| | |
|---|---|
| PRINT DATE: | 2020-06-19 03:57:04 +0200 |
| JOB ID: | c75ec994-28cf-45e2-94e6-5a4f6afce39a |
| RESULT PAGE: | https://tygs.dsmz.de/user_results/show?guid=c75ec994-28cf-45e2-94e6-5a4f6afce39a |

#### Table 1: Phylogenies

**Publication-ready versions** of both the genome-scale GBDP tree and the 16S rRNA gene sequence tree can be customized and exported either in SVG (vector graphic) or PNG format from within the phylogeny viewers in your TYGS result page. For publications the **SVG format is recommended** because it is lossless, always keeps its high resolution and can also be easily converted to other popular formats such as PDF or EPS. Please follow the link provided above!

#### Table 2: Identification

The below list contains the result of the TYGS species identification routine.

Explanation of remarks that might occur in the below table:

**remark [R1]:** The TYGS type strain database is automatically updated on an almost daily basis. However, if a particular type strain genome is not available in the TYGS database, this can have several reasons which are detailed in the FAQ. You can request an extended 16S rRNA gene analysis via the 16S tree viewer found in your result page to detect **not yet genome-sequenced** type strains relevant for your study.

**remark [R2]:** > 70% dDDH value (formula $d_4$) and (almost) minimal dDDH values for gene-content formulae $d_0$ and $d_6$ indicate a potentially unreliable identification result and should thus be checked via the 16S rRNA gene sequence similarity. Such strong deviations can, in principle, be caused by sequence contamination.

**remark [R3]:** G+C content difference of > 1 % indicates a potentially unreliable identification result because within species G+C content varies no more than 1 %, if computed from genome sequences (PMID: 24505073).

| Strain | Conclusion | Identification result | Remark |
|---|---|---|---|
| **'CHUV2995'** | belongs to known species | *Corynebacterium belfantii* | |
| **'CMCNS703'** | belongs to known species | *Corynebacterium belfantii* | |
| **'FRC0043'** | belongs to known species | *Corynebacterium belfantii* | |
| **'631'** | belongs to known species | *Corynebacterium diphtheriae* | |
| **'1137'** | belongs to known species | *Corynebacterium diphtheriae* | |
| **'1556'** | belongs to known species | *Corynebacterium diphtheriae* | |
| **'1734'** | belongs to known species | *Corynebacterium diphtheriae* | |
| **'2937'** | belongs to known species | *Corynebacterium diphtheriae* | |
| **'4499'** | belongs to known species | *Corynebacterium diphtheriae* | |
| **'4663'** | belongs to known species | *Corynebacterium diphtheriae* | |
| **'4766'** | belongs to known species | *Corynebacterium diphtheriae* | |
| **'5130'** | belongs to known species | *Corynebacterium diphtheriae* | |

#### Table 3: Pairwise comparisons of user genomes vs. type-strain genomes

The following table contains the pairwise dDDH values between your user genomes and the selected type-strain genomes. The dDDH values are provided along with their confidence intervals (C.I.) for the three different GBDP formulas:

- formula $d_0$ (a.k.a. GGDC formula 1): length of all HSPs divided by total genome length
- formula $d_4$ (a.k.a. GGDC formula 2): sum of all identities found in HSPs divided by overall HSP length
- formula $d_6$ (a.k.a. GGDC formula 3): sum of all identities found in HSPs divided by total genome length

**Note:** Formula $d_4$ is independent of genome length and is thus robust against the use of incomplete draft genomes. For other reasons for preferring formula $d_4$, see the FAQ.

| Query | Subject | $d_0$ | C.I. $d_0$ | $d_4$ | C.I. $d_4$ | $d_6$ | C.I. $d_6$ | Diff. G+C Percent |
|---|---|---|---|---|---|---|---|---|
| **'1137'** | **'1556'** | 99.6 | [99.3 - 99.8] | 100.0 | [99.9 - 100.0] | 99.9 | [99.7 - 99.9] | 0.01 |
| **'1556'** | **'4499'** | 99.6 | [99.3 - 99.8] | 100.0 | [99.9 - 100.0] | 99.9 | [99.7 - 99.9] | 0.01 |
| **'4499'** | **'4766'** | 99.7 | [99.3 - 99.8] | 100.0 | [100.0 - 100.0] | 99.9 | [99.8 - 99.9] | 0.09 |
| **'FRC0043'** | *Corynebacterium belfantii* FRC0043 | 100.0 | [100.0 - 100.0] | 100.0 | [100.0 - 100.0] | 100.0 | [100.0 - 100.0] | 0.01 |
| **'4663'** | **'5130'** | 100.0 | [100.0 - 100.0] | 100.0 | [99.9 - 100.0] | 100.0 | [100.0 - 100.0] | 0.01 |
| **'1556'** | **'4766'** | 98.9 | [98.1 - 99.4] | 100.0 | [99.9 - 100.0] | 99.5 | [99.2 - 99.7] | 0.11 |
| **'631'** | **'1137'** | 100.0 | [100.0 - 100.0] | 100.0 | [100.0 - 100.0] | 100.0 | [100.0 - 100.0] | 0.0 |
| **'631'** | **'4499'** | 100.0 | [100.0 - 100.0] | 100.0 | [100.0 - 100.0] | 100.0 | [100.0 - 100.0] | 0.0 |
| **'4766'** | **'5130'** | 99.7 | [99.3 - 99.9] | 100.0 | [99.9 - 100.0] | 99.9 | [99.8 - 99.9] | 0.1 |
| **'1734'** | **'5130'** | 100.0 | [100.0 - 100.0] | 100.0 | [99.9 - 100.0] | 100.0 | [100.0 - 100.0] | 0.01 |
| **'1734'** | **'4663'** | 100.0 | [100.0 - 100.0] | 100.0 | [99.9 - 100.0] | 100.0 | [100.0 - 100.0] | 0.0 |
| **'CHUV2995'** | *Corynebacterium diphtheriae* subsp. *lausannense* CHUV2995 | 100.0 | [100.0 - 100.0] | 100.0 | [100.0 - 100.0] | 100.0 | [100.0 - 100.0] | 0.0 |
| **'631'** | **'1556'** | 99.6 | [99.3 - 99.8] | 100.0 | [100.0 - 100.0] | 99.9 | [99.7 - 99.9] | 0.01 |
| **'1137'** | **'4499'** | 100.0 | [100.0 - 100.0] | 100.0 | [99.9 - 100.0] | 100.0 | [100.0 - 100.0] | 0.0 |
| **'631'** | **'4766'** | 99.7 | [99.3 - 99.8] | 100.0 | [99.9 - 100.0] | 99.9 | [99.7 - 99.9] | 0.09 |
| **'4663'** | **'4766'** | 99.7 | [99.3 - 99.8] | 100.0 | [100.0 - 100.0] | 99.9 | [99.7 - 99.9] | 0.09 |
| **'1556'** | **'5130'** | 99.6 | [99.2 - 99.8] | 99.9 | [99.9 - 100.0] | 99.8 | [99.7 - 99.9] | 0.01 |
| **'1734'** | **'4766'** | 99.7 | [99.3 - 99.8] | 99.9 | [99.9 - 100.0] | 99.9 | [99.7 - 99.9] | 0.09 |
| **'1556'** | **'4663'** | 99.6 | [99.2 - 99.8] | 99.9 | [99.8 - 99.9] | 99.9 | [99.7 - 99.9] | 0.02 |
| **'1137'** | **'4766'** | 99.7 | [99.3 - 99.8] | 99.9 | [99.9 - 100.0] | 99.9 | [99.7 - 99.9] | 0.09 |
| **'1556'** | **'1734'** | 99.6 | [99.2 - 99.8] | 99.9 | [99.9 - 100.0] | 99.9 | [99.7 - 99.9] | 0.01 |

| Query | Subject | $d_0$ | C.I. $d_0$ | $d_4$ | C.I. $d_4$ | $d_6$ | C.I. $d_6$ | Diff. G+C Percent |
|---|---|---|---|---|---|---|---|---|
| **'631'** | **'4663'** | 100.0 | [100.0 - 100.0] | 99.8 | [99.6 - 99.9] | 100.0 | [100.0 - 100.0] | 0.0 |
| **'4499'** | **'4663'** | 100.0 | [100.0 - 100.0] | 99.8 | [99.6 - 99.9] | 100.0 | [100.0 - 100.0] | 0.0 |
| **'4499'** | **'5130'** | 100.0 | [100.0 - 100.0] | 99.8 | [99.7 - 99.9] | 100.0 | [100.0 - 100.0] | 0.01 |
| **'1734'** | **'4499'** | 100.0 | [100.0 - 100.0] | 99.8 | [99.6 - 99.9] | 100.0 | [100.0 - 100.0] | 0.0 |
| **'1137'** | **'5130'** | 100.0 | [100.0 - 100.0] | 99.8 | [99.6 - 99.9] | 100.0 | [100.0 - 100.0] | 0.01 |
| **'631'** | **'5130'** | 100.0 | [100.0 - 100.0] | 99.8 | [99.7 - 99.9] | 100.0 | [100.0 - 100.0] | 0.01 |
| **'631'** | **'1734'** | 100.0 | [100.0 - 100.0] | 99.8 | [99.7 - 99.9] | 100.0 | [100.0 - 100.0] | 0.0 |
| **'1137'** | **'1734'** | 100.0 | [100.0 - 100.0] | 99.7 | [99.6 - 99.8] | 100.0 | [100.0 - 100.0] | 0.0 |
| **'1137'** | **'4663'** | 100.0 | [100.0 - 100.0] | 99.6 | [99.3 - 99.7] | 100.0 | [100.0 - 100.0] | 0.0 |
| **'CHUV2995'** | **'CMCNS703'** | 95.6 | [93.5 - 97.1] | 94.1 | [92.3 - 95.5] | 97.1 | [95.7 - 98.0] | 0.21 |
| **'CMCNS703'** | *Corynebacterium belfantii* FRC0043 | 94.4 | [92.0 - 96.2] | 94.1 | [92.3 - 95.5] | 96.2 | [94.6 - 97.4] | 0.11 |
| **'CMCNS703'** | *Corynebacterium diphtheriae* subsp. *lausannense* CHUV2995 | 95.6 | [93.5 - 97.1] | 94.1 | [92.3 - 95.5] | 97.1 | [95.7 - 98.0] | 0.21 |
| **'CMCNS703'** | **'FRC0043'** | 94.4 | [92.0 - 96.2] | 94.1 | [92.3 - 95.5] | 96.2 | [94.6 - 97.4] | 0.11 |
| **'CHUV2995'** | *Corynebacterium belfantii* FRC0043 | 92.5 | [89.7 - 94.6] | 88.5 | [86.1 - 90.6] | 94.2 | [92.1 - 95.8] | 0.31 |
| **'FRC0043'** | *Corynebacterium diphtheriae* subsp. *lausannense* CHUV2995 | 92.5 | [89.7 - 94.6] | 88.5 | [86.1 - 90.6] | 94.2 | [92.1 - 95.8] | 0.32 |
| **'CHUV2995'** | **'FRC0043'** | 92.5 | [89.7 - 94.6] | 88.5 | [86.1 - 90.6] | 94.2 | [92.1 - 95.8] | 0.32 |
| **'4663'** | *Corynebacterium diphtheriae* NCTC 11397 | 94.7 | [92.3 - 96.4] | 86.9 | [84.4 - 89.2] | 95.6 | [93.8 - 96.9] | 0.08 |
| **'5130'** | *Corynebacterium diphtheriae* NCTC 11397 | 94.7 | [92.3 - 96.4] | 86.9 | [84.3 - 89.1] | 95.6 | [93.8 - 96.9] | 0.09 |
| **'4766'** | *Corynebacterium diphtheriae* NCTC 11397 | 94.3 | [91.8 - 96.0] | 86.9 | [84.3 - 89.1] | 95.3 | [93.4 - 96.6] | 0.01 |
| **'1734'** | *Corynebacterium diphtheriae* NCTC 11397 | 94.7 | [92.4 - 96.4] | 86.8 | [84.2 - 89.0] | 95.6 | [93.8 - 96.9] | 0.08 |
| **'631'** | *Corynebacterium diphtheriae* NCTC 11397 | 94.8 | [92.4 - 96.5] | 86.8 | [84.2 - 89.0] | 95.7 | [93.9 - 96.9] | 0.08 |
| **'1137'** | *Corynebacterium diphtheriae* NCTC 11397 | 94.8 | [92.4 - 96.5] | 86.8 | [84.2 - 89.0] | 95.7 | [93.9 - 96.9] | 0.08 |
| **'4499'** | *Corynebacterium diphtheriae* NCTC 11397 | 94.8 | [92.4 - 96.5] | 86.7 | [84.1 - 89.0] | 95.7 | [93.9 - 96.9] | 0.08 |
| **'1556'** | *Corynebacterium diphtheriae* NCTC 11397 | 93.8 | [91.1 - 95.6] | 86.6 | [84.0 - 88.8] | 94.9 | [92.9 - 96.3] | 0.1 |
| **'2937'** | *Corynebacterium diphtheriae* NCTC 11397 | 92.1 | [89.1 - 94.3] | 85.3 | [82.6 - 87.6] | 93.5 | [91.2 - 95.2] | 0.04 |
| **'2937'** | **'5130'** | 94.8 | [92.4 - 96.4] | 85.1 | [82.4 - 87.4] | 95.4 | [93.6 - 96.8] | 0.05 |
| **'631'** | **'2937'** | 94.8 | [92.5 - 96.5] | 85.1 | [82.3 - 87.4] | 95.5 | [93.7 - 96.8] | 0.05 |

| Query | Subject | $d_0$ | C.I. $d_0$ | $d_4$ | C.I. $d_4$ | $d_6$ | C.I. $d_6$ | Diff. G+C Percent |
|---|---|---|---|---|---|---|---|---|
| **'2937'** | **'4499'** | 94.8 | [92.5 - 96.5] | 85.1 | [82.4 - 87.4] | 95.5 | [93.7 - 96.8] | 0.05 |
| **'2937'** | **'4766'** | 94.1 | [91.6 - 95.9] | 85.1 | [82.4 - 87.5] | 95.0 | [93.0 - 96.4] | 0.05 |
| **'2937'** | **'4663'** | 94.8 | [92.5 - 96.5] | 85.1 | [82.4 - 87.5] | 95.5 | [93.7 - 96.8] | 0.04 |
| **'1137'** | **'2937'** | 94.8 | [92.5 - 96.5] | 85.1 | [82.3 - 87.4] | 95.5 | [93.7 - 96.8] | 0.05 |
| **'1734'** | **'2937'** | 94.8 | [92.5 - 96.5] | 85.0 | [82.3 - 87.4] | 95.5 | [93.6 - 96.8] | 0.05 |
| **'1556'** | **'2937'** | 93.4 | [90.8 - 95.4] | 84.9 | [82.1 - 87.2] | 94.4 | [92.4 - 95.9] | 0.06 |
| **'2937'** | **'CMCNS703'** | 74.5 | [70.6 - 78.2] | 63.4 | [60.5 - 66.2] | 74.9 | [71.4 - 78.1] | 0.17 |
| **'CMCNS703'** | *Corynebacterium diphtheriae* NCTC 11397 | 76.1 | [72.1 - 79.7] | 63.1 | [60.2 - 65.9] | 76.1 | [72.7 - 79.3] | 0.21 |
| **'5130'** | **'CMCNS703'** | 75.6 | [71.6 - 79.2] | 62.9 | [60.0 - 65.7] | 75.6 | [72.2 - 78.8] | 0.12 |
| **'CHUV2995'** | *Corynebacterium diphtheriae* NCTC 11397 | 71.1 | [67.2 - 74.8] | 62.9 | [60.0 - 65.7] | 71.8 | [68.3 - 75.0] | 0.42 |
| **'4663'** | **'CMCNS703'** | 75.5 | [71.6 - 79.1] | 62.9 | [60.0 - 65.7] | 75.6 | [72.1 - 78.8] | 0.13 |
| **'4499'** | **'CMCNS703'** | 75.6 | [71.6 - 79.2] | 62.8 | [59.9 - 65.6] | 75.7 | [72.2 - 78.8] | 0.12 |
| **'2937'** | **'CHUV2995'** | 69.0 | [65.1 - 72.6] | 62.8 | [59.9 - 65.6] | 69.9 | [66.5 - 73.1] | 0.38 |
| **'1734'** | **'CMCNS703'** | 75.5 | [71.5 - 79.1] | 62.8 | [59.9 - 65.6] | 75.6 | [72.1 - 78.8] | 0.12 |
| **'2937'** | *Corynebacterium diphtheriae* subsp. *lausannense* CHUV2995 | 69.0 | [65.1 - 72.6] | 62.8 | [59.9 - 65.6] | 69.9 | [66.5 - 73.1] | 0.38 |
| **'1137'** | **'CMCNS703'** | 75.6 | [71.6 - 79.2] | 62.8 | [59.9 - 65.6] | 75.7 | [72.2 - 78.8] | 0.12 |
| **'1556'** | **'CMCNS703'** | 75.4 | [71.4 - 79.0] | 62.8 | [59.9 - 65.7] | 75.5 | [72.0 - 78.7] | 0.11 |
| **'631'** | **'CMCNS703'** | 75.6 | [71.6 - 79.2] | 62.8 | [59.9 - 65.7] | 75.6 | [72.2 - 78.8] | 0.12 |
| **'4766'** | **'CMCNS703'** | 74.9 | [70.9 - 78.5] | 62.8 | [59.9 - 65.6] | 75.0 | [71.6 - 78.2] | 0.22 |
| **'1137'** | *Corynebacterium diphtheriae* subsp. *lausannense* CHUV2995 | 70.7 | [66.8 - 74.3] | 62.3 | [59.4 - 65.1] | 71.3 | [67.8 - 74.5] | 0.33 |
| **'4499'** | *Corynebacterium diphtheriae* subsp. *lausannense* CHUV2995 | 70.7 | [66.7 - 74.3] | 62.3 | [59.4 - 65.1] | 71.3 | [67.8 - 74.5] | 0.33 |
| **'631'** | **'CHUV2995'** | 70.7 | [66.7 - 74.3] | 62.3 | [59.4 - 65.1] | 71.3 | [67.8 - 74.5] | 0.33 |
| **'1734'** | *Corynebacterium diphtheriae* subsp. *lausannense* CHUV2995 | 70.6 | [66.7 - 74.3] | 62.3 | [59.4 - 65.1] | 71.2 | [67.8 - 74.5] | 0.33 |
| **'1137'** | **'CHUV2995'** | 70.7 | [66.8 - 74.3] | 62.3 | [59.4 - 65.1] | 71.3 | [67.8 - 74.5] | 0.33 |
| **'5130'** | **'CHUV2995'** | 70.6 | [66.7 - 74.3] | 62.3 | [59.4 - 65.1] | 71.3 | [67.8 - 74.5] | 0.33 |
| **'4663'** | *Corynebacterium diphtheriae* subsp. *lausannense* CHUV2995 | 70.6 | [66.7 - 74.3] | 62.3 | [59.4 - 65.1] | 71.2 | [67.8 - 74.5] | 0.34 |

| Query | Subject | $d_0$ | C.I. $d_0$ | $d_4$ | C.I. $d_4$ | $d_6$ | C.I. $d_6$ | Diff. G+C Percent |
|---|---|---|---|---|---|---|---|---|
| **'5130'** | *Corynebacterium diphtheriae* subsp. *lausannense* CHUV2995 | 70.6 | [66.7 - 74.3] | 62.3 | [59.4 - 65.1] | 71.3 | [67.8 - 74.5] | 0.33 |
| **'4766'** | **'CHUV2995'** | 70.0 | [66.0 - 73.6] | 62.3 | [59.4 - 65.1] | 70.7 | [67.2 - 73.9] | 0.43 |
| **'4663'** | **'CHUV2995'** | 70.6 | [66.7 - 74.3] | 62.3 | [59.4 - 65.1] | 71.2 | [67.8 - 74.5] | 0.34 |
| **'4766'** | *Corynebacterium diphtheriae* subsp. *lausannense* CHUV2995 | 70.0 | [66.0 - 73.6] | 62.3 | [59.4 - 65.1] | 70.7 | [67.2 - 73.9] | 0.43 |
| **'631'** | *Corynebacterium diphtheriae* subsp. *lausannense* CHUV2995 | 70.7 | [66.7 - 74.3] | 62.3 | [59.4 - 65.1] | 71.3 | [67.8 - 74.5] | 0.33 |
| **'4499'** | **'CHUV2995'** | 70.7 | [66.7 - 74.3] | 62.3 | [59.4 - 65.1] | 71.3 | [67.8 - 74.5] | 0.33 |
| **'1734'** | **'CHUV2995'** | 70.6 | [66.7 - 74.3] | 62.3 | [59.4 - 65.1] | 71.2 | [67.8 - 74.5] | 0.33 |
| **'1556'** | *Corynebacterium diphtheriae* subsp. *lausannense* CHUV2995 | 70.3 | [66.3 - 73.9] | 62.2 | [59.4 - 65.0] | 70.9 | [67.5 - 74.2] | 0.32 |
| **'1556'** | **'CHUV2995'** | 70.3 | [66.3 - 73.9] | 62.2 | [59.4 - 65.0] | 70.9 | [67.5 - 74.2] | 0.32 |
| **'2937'** | *Corynebacterium belfantii* FRC0043 | 76.7 | [72.7 - 80.3] | 62.0 | [59.1 - 64.8] | 76.4 | [72.9 - 79.6] | 0.07 |
| **'2937'** | **'FRC0043'** | 76.7 | [72.7 - 80.3] | 62.0 | [59.1 - 64.8] | 76.4 | [72.9 - 79.6] | 0.06 |
| **'FRC0043'** | *Corynebacterium diphtheriae* NCTC 11397 | 77.0 | [73.1 - 80.6] | 61.9 | [59.0 - 64.7] | 76.6 | [73.2 - 79.8] | 0.1 |
| **'5130'** | *Corynebacterium belfantii* FRC0043 | 77.3 | [73.3 - 80.8] | 61.6 | [58.7 - 64.4] | 76.8 | [73.3 - 79.9] | 0.01 |
| **'5130'** | **'FRC0043'** | 77.3 | [73.3 - 80.8] | 61.6 | [58.7 - 64.4] | 76.8 | [73.3 - 79.9] | 0.01 |
| **'4663'** | **'FRC0043'** | 77.3 | [73.3 - 80.8] | 61.6 | [58.7 - 64.3] | 76.7 | [73.3 - 79.9] | 0.02 |
| **'4663'** | *Corynebacterium belfantii* FRC0043 | 77.3 | [73.3 - 80.8] | 61.6 | [58.7 - 64.3] | 76.7 | [73.3 - 79.9] | 0.02 |
| **'1556'** | *Corynebacterium belfantii* FRC0043 | 76.0 | [72.1 - 79.6] | 61.5 | [58.7 - 64.3] | 75.7 | [72.2 - 78.9] | 0.0 |
| **'1137'** | *Corynebacterium belfantii* FRC0043 | 77.3 | [73.3 - 80.9] | 61.5 | [58.6 - 64.3] | 76.8 | [73.3 - 79.9] | 0.02 |
| **'4499'** | **'FRC0043'** | 77.3 | [73.3 - 80.8] | 61.5 | [58.7 - 64.3] | 76.8 | [73.3 - 79.9] | 0.01 |
| **'631'** | **'FRC0043'** | 77.3 | [73.3 - 80.8] | 61.5 | [58.7 - 64.3] | 76.8 | [73.3 - 79.9] | 0.01 |
| **'4499'** | *Corynebacterium belfantii* FRC0043 | 77.3 | [73.3 - 80.8] | 61.5 | [58.7 - 64.3] | 76.8 | [73.3 - 79.9] | 0.02 |
| **'1137'** | **'FRC0043'** | 77.3 | [73.3 - 80.9] | 61.5 | [58.6 - 64.3] | 76.8 | [73.3 - 79.9] | 0.01 |
| **'4766'** | **'FRC0043'** | 76.6 | [72.6 - 80.1] | 61.5 | [58.6 - 64.3] | 76.1 | [72.7 - 79.3] | 0.11 |
| **'631'** | *Corynebacterium belfantii* FRC0043 | 77.3 | [73.3 - 80.8] | 61.5 | [58.7 - 64.3] | 76.8 | [73.3 - 79.9] | 0.02 |
| **'1734'** | *Corynebacterium belfantii* FRC0043 | 77.3 | [73.3 - 80.8] | 61.5 | [58.6 - 64.3] | 76.7 | [73.3 - 79.9] | 0.02 |
| **'1556'** | **'FRC0043'** | 76.0 | [72.1 - 79.6] | 61.5 | [58.7 - 64.3] | 75.7 | [72.2 - 78.9] | 0.0 |

| Query | Subject | $d_0$ | C.I. $d_0$ | $d_4$ | C.I. $d_4$ | $d_6$ | C.I. $d_6$ | Diff. G+C Percent |
|---|---|---|---|---|---|---|---|---|
| **'1734'** | **'FRC0043'** | 77.3 | [73.3 - 80.8] | 61.5 | [58.6 - 64.3] | 76.7 | [73.3 - 79.9] | 0.01 |
| **'4766'** | *Corynebacterium belfantii* FRC0043 | 76.6 | [72.6 - 80.1] | 61.5 | [58.6 - 64.3] | 76.1 | [72.7 - 79.3] | 0.11 |
| **'4663'** | *Corynebacterium rouxii* FRC0190 T | 78.0 | [74.0 - 81.5] | 49.1 | [46.5 - 51.7] | 73.6 | [70.1 - 76.8] | 0.38 |
| **'5130'** | *Corynebacterium rouxii* FRC0190 T | 77.9 | [73.9 - 81.4] | 49.1 | [46.4 - 51.7] | 73.5 | [70.0 - 76.7] | 0.38 |
| **'1734'** | *Corynebacterium rouxii* FRC0190 T | 78.0 | [74.1 - 81.5] | 49.0 | [46.4 - 51.7] | 73.6 | [70.2 - 76.8] | 0.38 |
| **'4766'** | *Corynebacterium rouxii* FRC0190 T | 77.6 | [73.6 - 81.1] | 49.0 | [46.4 - 51.6] | 73.2 | [69.8 - 76.4] | 0.29 |
| **'4499'** | *Corynebacterium rouxii* FRC0190 T | 78.1 | [74.1 - 81.6] | 49.0 | [46.4 - 51.6] | 73.7 | [70.2 - 76.9] | 0.38 |
| **'1137'** | *Corynebacterium rouxii* FRC0190 T | 78.1 | [74.1 - 81.6] | 49.0 | [46.4 - 51.6] | 73.6 | [70.2 - 76.8] | 0.38 |
| **'1556'** | *Corynebacterium rouxii* FRC0190 T | 76.5 | [72.5 - 80.1] | 49.0 | [46.4 - 51.6] | 72.4 | [68.9 - 75.6] | 0.39 |
| **'631'** | *Corynebacterium rouxii* FRC0190 T | 78.1 | [74.1 - 81.6] | 49.0 | [46.4 - 51.6] | 73.6 | [70.2 - 76.9] | 0.38 |
| **'2937'** | *Corynebacterium rouxii* FRC0190 T | 77.7 | [73.7 - 81.2] | 48.7 | [46.1 - 51.3] | 73.2 | [69.8 - 76.4] | 0.33 |
| **'CHUV2995'** | *Corynebacterium rouxii* FRC0190 T | 61.5 | [57.8 - 65.1] | 45.7 | [43.1 - 48.2] | 59.0 | [55.7 - 62.1] | 0.71 |
| **'CMCNS703'** | *Corynebacterium rouxii* FRC0190 T | 65.7 | [61.9 - 69.3] | 45.5 | [43.0 - 48.1] | 62.3 | [59.0 - 65.5] | 0.5 |
| **'FRC0043'** | *Corynebacterium rouxii* FRC0190 T | 73.0 | [69.0 - 76.6] | 45.4 | [42.8 - 47.9] | 68.2 | [64.8 - 71.4] | 0.39 |
| **'1734'** | *Corynebacterium singulare* DSM 44357 | 13.3 | [10.5 - 16.6] | 34.3 | [31.9 - 36.8] | 13.7 | [11.3 - 16.5] | 6.51 |
| **'5130'** | *Corynebacterium singulare* DSM 44357 | 13.3 | [10.5 - 16.6] | 34.3 | [31.9 - 36.9] | 13.7 | [11.3 - 16.5] | 6.51 |
| **'1556'** | *Corynebacterium singulare* DSM 44357 | 13.3 | [10.5 - 16.6] | 34.3 | [31.9 - 36.8] | 13.7 | [11.3 - 16.5] | 6.5 |
| **'4499'** | *Corynebacterium singulare* DSM 44357 | 13.3 | [10.5 - 16.6] | 34.3 | [31.9 - 36.8] | 13.7 | [11.3 - 16.5] | 6.51 |
| **'1137'** | *Corynebacterium singulare* DSM 44357 | 13.3 | [10.5 - 16.6] | 34.3 | [31.9 - 36.8] | 13.7 | [11.3 - 16.5] | 6.51 |
| **'631'** | *Corynebacterium singulare* DSM 44357 | 13.3 | [10.5 - 16.6] | 34.3 | [31.9 - 36.8] | 13.7 | [11.3 - 16.5] | 6.51 |
| **'4766'** | *Corynebacterium singulare* DSM 44357 | 13.3 | [10.5 - 16.6] | 34.3 | [31.9 - 36.8] | 13.7 | [11.3 - 16.5] | 6.6 |
| **'4663'** | *Corynebacterium singulare* DSM 44357 | 13.3 | [10.5 - 16.6] | 34.3 | [31.9 - 36.9] | 13.7 | [11.3 - 16.5] | 6.51 |
| **'2937'** | *Corynebacterium singulare* DSM 44357 | 13.3 | [10.6 - 16.6] | 33.4 | [31.0 - 35.9] | 13.7 | [11.3 - 16.5] | 6.56 |
| **'CHUV2995'** | *Corynebacterium striatum* ATCC 6940 | 13.4 | [10.6 - 16.7] | 31.8 | [29.4 - 34.3] | 13.8 | [11.4 - 16.6] | 5.48 |
| **'CMCNS703'** | *Corynebacterium striatum* ATCC 6940 | 13.3 | [10.6 - 16.7] | 31.4 | [29.0 - 33.9] | 13.7 | [11.4 - 16.5] | 5.69 |
| **'4766'** | *Corynebacterium hadale* NBT06-6 | 13.0 | [10.3 - 16.3] | 30.2 | [27.8 - 32.7] | 13.4 | [11.0 - 16.1] | 11.66 |
| **'1137'** | *Corynebacterium hadale* NBT06-6 | 13.0 | [10.3 - 16.2] | 30.1 | [27.7 - 32.6] | 13.4 | [11.0 - 16.1] | 11.57 |

| Query | Subject | $d_0$ | C.I. $d_0$ | $d_4$ | C.I. $d_4$ | $d_6$ | C.I. $d_6$ | Diff. G+C Percent |
|---|---|---|---|---|---|---|---|---|
| **'631'** | *Corynebacterium hadale* NBT06-6 | 13.0 | [10.3 - 16.2] | 30.1 | [27.7 - 32.6] | 13.4 | [11.0 - 16.1] | 11.57 |
| **'4663'** | *Corynebacterium hadale* NBT06-6 | 13.0 | [10.3 - 16.2] | 30.1 | [27.7 - 32.6] | 13.4 | [11.0 - 16.1] | 11.57 |
| **'1556'** | *Corynebacterium hadale* NBT06-6 | 13.0 | [10.2 - 16.2] | 30.1 | [27.7 - 32.6] | 13.4 | [11.0 - 16.1] | 11.55 |
| **'5130'** | *Corynebacterium hadale* NBT06-6 | 13.0 | [10.3 - 16.2] | 30.1 | [27.7 - 32.6] | 13.4 | [11.0 - 16.1] | 11.56 |
| **'1734'** | *Corynebacterium hadale* NBT06-6 | 13.0 | [10.3 - 16.3] | 30.1 | [27.7 - 32.6] | 13.4 | [11.0 - 16.1] | 11.57 |
| **'4499'** | *Corynebacterium hadale* NBT06-6 | 13.0 | [10.3 - 16.2] | 30.1 | [27.7 - 32.6] | 13.4 | [11.0 - 16.1] | 11.57 |
| **'CMCNS703'** | *Corynebacterium singulare* DSM 44357 | 13.4 | [10.6 - 16.7] | 30.0 | [27.6 - 32.5] | 13.8 | [11.4 - 16.6] | 6.39 |
| **'CHUV2995'** | *Corynebacterium singulare* DSM 44357 | 13.4 | [10.6 - 16.7] | 29.7 | [27.4 - 32.2] | 13.7 | [11.4 - 16.5] | 6.18 |
| **'CMCNS703'** | *Corynebacterium phoceense* MC1 | 13.3 | [10.6 - 16.6] | 28.9 | [26.5 - 31.4] | 13.7 | [11.3 - 16.5] | 9.49 |
| **'5130'** | *Corynebacterium phoceense* MC1 | 13.2 | [10.5 - 16.6] | 28.9 | [26.5 - 31.4] | 13.6 | [11.3 - 16.4] | 9.61 |
| **'2937'** | *Corynebacterium hadale* NBT06-6 | 13.0 | [10.3 - 16.3] | 28.9 | [26.6 - 31.4] | 13.4 | [11.0 - 16.2] | 11.61 |
| **'4766'** | *Corynebacterium phoceense* MC1 | 13.3 | [10.5 - 16.6] | 28.8 | [26.5 - 31.3] | 13.6 | [11.3 - 16.4] | 9.71 |
| **'4499'** | *Corynebacterium phoceense* MC1 | 13.2 | [10.5 - 16.6] | 28.8 | [26.4 - 31.3] | 13.6 | [11.3 - 16.4] | 9.62 |
| **'631'** | *Corynebacterium phoceense* MC1 | 13.2 | [10.5 - 16.6] | 28.8 | [26.5 - 31.3] | 13.6 | [11.3 - 16.4] | 9.62 |
| **'4663'** | *Corynebacterium phoceense* MC1 | 13.2 | [10.5 - 16.6] | 28.8 | [26.5 - 31.3] | 13.6 | [11.3 - 16.4] | 9.62 |
| **'1556'** | *Corynebacterium phoceense* MC1 | 13.2 | [10.5 - 16.6] | 28.8 | [26.4 - 31.3] | 13.6 | [11.3 - 16.4] | 9.6 |
| **'1137'** | *Corynebacterium phoceense* MC1 | 13.2 | [10.5 - 16.6] | 28.8 | [26.5 - 31.3] | 13.6 | [11.3 - 16.4] | 9.62 |
| **'CHUV2995'** | *Corynebacterium phoceense* MC1 | 13.3 | [10.5 - 16.6] | 28.8 | [26.5 - 31.3] | 13.7 | [11.3 - 16.5] | 9.28 |
| **'2937'** | *Corynebacterium phoceense* MC1 | 13.3 | [10.5 - 16.6] | 28.7 | [26.3 - 31.2] | 13.7 | [11.3 - 16.4] | 9.66 |
| **'CMCNS703'** | *Corynebacterium hadale* NBT06-6 | 13.1 | [10.4 - 16.4] | 28.6 | [26.2 - 31.1] | 13.5 | [11.1 - 16.3] | 11.44 |
| **'1734'** | *Corynebacterium phoceense* MC1 | 13.2 | [10.5 - 16.6] | 28.6 | [26.3 - 31.1] | 13.6 | [11.3 - 16.4] | 9.62 |
| **'CHUV2995'** | *Corynebacterium hadale* NBT06-6 | 13.0 | [10.3 - 16.3] | 28.2 | [25.9 - 30.7] | 13.4 | [11.1 - 16.2] | 11.23 |
| **'5130'** | *Corynebacterium striatum* ATCC 6940 | 13.2 | [10.5 - 16.5] | 27.6 | [25.3 - 30.1] | 13.6 | [11.2 - 16.4] | 5.81 |
| **'1734'** | *Corynebacterium striatum* ATCC 6940 | 13.2 | [10.5 - 16.5] | 27.6 | [25.3 - 30.1] | 13.6 | [11.2 - 16.4] | 5.81 |
| **'4499'** | *Corynebacterium striatum* ATCC 6940 | 13.2 | [10.5 - 16.5] | 27.6 | [25.3 - 30.1] | 13.6 | [11.2 - 16.4] | 5.81 |
| **'1556'** | *Corynebacterium striatum* ATCC 6940 | 13.2 | [10.5 - 16.5] | 27.6 | [25.2 - 30.1] | 13.6 | [11.2 - 16.4] | 5.8 |
| **'631'** | *Corynebacterium striatum* ATCC 6940 | 13.2 | [10.5 - 16.5] | 27.6 | [25.3 - 30.1] | 13.6 | [11.2 - 16.4] | 5.81 |

| Query | Subject | $d_0$ | C.I. $d_0$ | $d_4$ | C.I. $d_4$ | $d_6$ | C.I. $d_6$ | Diff. G+C Percent |
|---|---|---|---|---|---|---|---|---|
| **'1137'** | *Corynebacterium striatum* ATCC 6940 | 13.2 | [10.5 - 16.5] | 27.6 | [25.3 - 30.1] | 13.6 | [11.2 - 16.4] | 5.81 |
| **'4663'** | *Corynebacterium striatum* ATCC 6940 | 13.2 | [10.5 - 16.5] | 27.6 | [25.3 - 30.1] | 13.6 | [11.2 - 16.4] | 5.81 |
| **'4766'** | *Corynebacterium striatum* ATCC 6940 | 13.2 | [10.5 - 16.5] | 26.6 | [24.2 - 29.1] | 13.6 | [11.2 - 16.4] | 5.9 |
| **'CHUV2995'** | *Corynebacterium kutscheri* DSM 20755 | 13.1 | [10.4 - 16.4] | 26.6 | [24.3 - 29.1] | 13.5 | [11.1 - 16.2] | 7.48 |
| **'CMCNS703'** | *Corynebacterium kutscheri* DSM 20755 | 13.1 | [10.4 - 16.4] | 26.2 | [23.9 - 28.7] | 13.5 | [11.1 - 16.3] | 7.27 |
| **'2937'** | *Corynebacterium striatum* ATCC 6940 | 13.1 | [10.4 - 16.4] | 26.0 | [23.7 - 28.5] | 13.5 | [11.2 - 16.3] | 5.86 |
| **'FRC0043'** | *Corynebacterium striatum* ATCC 6940 | 13.1 | [10.4 - 16.4] | 25.8 | [23.5 - 28.3] | 13.5 | [11.2 - 16.3] | 5.8 |
| **'CHUV2995'** | *Corynebacterium vitaeruminis* DSM 20294 | 13.4 | [10.7 - 16.8] | 25.2 | [22.9 - 27.7] | 13.8 | [11.4 - 16.6] | 11.58 |
| **'2937'** | *Corynebacterium argentoratense* DSM 44202 | 13.2 | [10.5 - 16.5] | 25.2 | [22.8 - 27.7] | 13.6 | [11.2 - 16.4] | 5.34 |
| **'4766'** | *Corynebacterium argentoratense* DSM 44202 | 13.2 | [10.5 - 16.6] | 25.1 | [22.8 - 27.6] | 13.6 | [11.2 - 16.4] | 5.39 |
| **'CMCNS703'** | *Corynebacterium vitaeruminis* DSM 20294 | 13.4 | [10.7 - 16.7] | 25.1 | [22.8 - 27.6] | 13.8 | [11.4 - 16.6] | 11.79 |
| **'FRC0043'** | *Corynebacterium phoceense* MC1 | 13.2 | [10.5 - 16.5] | 25.1 | [22.8 - 27.6] | 13.6 | [11.2 - 16.4] | 9.6 |
| **'1556'** | *Corynebacterium argentoratense* DSM 44202 | 13.2 | [10.5 - 16.5] | 25.0 | [22.7 - 27.5] | 13.6 | [11.2 - 16.4] | 5.28 |
| **'5130'** | *Corynebacterium argentoratense* DSM 44202 | 13.2 | [10.5 - 16.5] | 25.0 | [22.7 - 27.5] | 13.6 | [11.2 - 16.4] | 5.29 |
| **'4499'** | *Corynebacterium argentoratense* DSM 44202 | 13.2 | [10.5 - 16.5] | 25.0 | [22.7 - 27.5] | 13.6 | [11.2 - 16.4] | 5.29 |
| **'2937'** | *Corynebacterium kutscheri* DSM 20755 | 13.1 | [10.4 - 16.4] | 25.0 | [22.7 - 27.5] | 13.5 | [11.1 - 16.3] | 7.1 |
| **'1137'** | *Corynebacterium argentoratense* DSM 44202 | 13.2 | [10.5 - 16.5] | 25.0 | [22.7 - 27.5] | 13.6 | [11.2 - 16.4] | 5.29 |
| **'4663'** | *Corynebacterium argentoratense* DSM 44202 | 13.2 | [10.5 - 16.5] | 25.0 | [22.7 - 27.5] | 13.6 | [11.2 - 16.4] | 5.3 |
| **'631'** | *Corynebacterium argentoratense* DSM 44202 | 13.2 | [10.5 - 16.5] | 25.0 | [22.7 - 27.5] | 13.6 | [11.2 - 16.4] | 5.29 |
| **'1734'** | *Corynebacterium argentoratense* DSM 44202 | 13.2 | [10.5 - 16.5] | 25.0 | [22.7 - 27.5] | 13.6 | [11.2 - 16.4] | 5.29 |
| **'CMCNS703'** | *Corynebacterium tuscaniense* CCUG 51321 | 13.3 | [10.5 - 16.6] | 24.9 | [22.6 - 27.4] | 13.6 | [11.2 - 16.4] | 5.69 |
| **'FRC0043'** | *Corynebacterium kutscheri* DSM 20755 | 13.1 | [10.4 - 16.4] | 24.9 | [22.6 - 27.4] | 13.4 | [11.1 - 16.2] | 7.16 |
| **'FRC0043'** | *Corynebacterium vitaeruminis* DSM 20294 | 13.4 | [10.6 - 16.7] | 24.9 | [22.6 - 27.4] | 13.8 | [11.4 - 16.5] | 11.9 |
| **'4663'** | *Corynebacterium kutscheri* DSM 20755 | 13.1 | [10.4 - 16.4] | 24.8 | [22.5 - 27.3] | 13.5 | [11.1 - 16.3] | 7.14 |

| Query | Subject | $d_0$ | C.I. $d_0$ | $d_4$ | C.I. $d_4$ | $d_6$ | C.I. $d_6$ | Diff. G+C Percent |
|---|---|---|---|---|---|---|---|---|
| **'631'** | *Corynebacterium vitaeruminis* DSM 20294 | 13.3 | [10.6 - 16.7] | 24.8 | [22.5 - 27.3] | 13.7 | [11.3 - 16.5] | 11.92 |
| **'1556'** | *Corynebacterium kutscheri* DSM 20755 | 13.1 | [10.4 - 16.4] | 24.8 | [22.5 - 27.3] | 13.5 | [11.1 - 16.3] | 7.16 |
| **'4499'** | *Corynebacterium vitaeruminis* DSM 20294 | 13.3 | [10.6 - 16.7] | 24.8 | [22.5 - 27.3] | 13.7 | [11.3 - 16.5] | 11.92 |
| **'1137'** | *Corynebacterium kutscheri* DSM 20755 | 13.1 | [10.4 - 16.4] | 24.8 | [22.5 - 27.3] | 13.5 | [11.1 - 16.3] | 7.15 |
| **'4766'** | *Corynebacterium kutscheri* DSM 20755 | 13.1 | [10.4 - 16.4] | 24.8 | [22.5 - 27.3] | 13.5 | [11.1 - 16.3] | 7.05 |
| **'5130'** | *Corynebacterium kutscheri* DSM 20755 | 13.1 | [10.4 - 16.4] | 24.8 | [22.5 - 27.3] | 13.5 | [11.1 - 16.3] | 7.15 |
| **'4663'** | *Corynebacterium vitaeruminis* DSM 20294 | 13.3 | [10.6 - 16.7] | 24.8 | [22.5 - 27.3] | 13.7 | [11.3 - 16.5] | 11.92 |
| **'1734'** | *Corynebacterium vitaeruminis* DSM 20294 | 13.3 | [10.6 - 16.7] | 24.8 | [22.5 - 27.3] | 13.7 | [11.3 - 16.5] | 11.92 |
| **'5130'** | *Corynebacterium vitaeruminis* DSM 20294 | 13.3 | [10.6 - 16.7] | 24.8 | [22.5 - 27.3] | 13.7 | [11.3 - 16.5] | 11.91 |
| **'4499'** | *Corynebacterium kutscheri* DSM 20755 | 13.1 | [10.4 - 16.4] | 24.8 | [22.5 - 27.3] | 13.5 | [11.1 - 16.3] | 7.14 |
| **'1556'** | *Corynebacterium vitaeruminis* DSM 20294 | 13.3 | [10.6 - 16.7] | 24.8 | [22.5 - 27.3] | 13.7 | [11.3 - 16.5] | 11.9 |
| **'631'** | *Corynebacterium kutscheri* DSM 20755 | 13.1 | [10.4 - 16.4] | 24.8 | [22.5 - 27.3] | 13.5 | [11.1 - 16.3] | 7.15 |
| **'1137'** | *Corynebacterium vitaeruminis* DSM 20294 | 13.3 | [10.6 - 16.7] | 24.8 | [22.5 - 27.3] | 13.7 | [11.3 - 16.5] | 11.92 |
| **'4766'** | *Corynebacterium vitaeruminis* DSM 20294 | 13.4 | [10.6 - 16.7] | 24.8 | [22.5 - 27.3] | 13.7 | [11.3 - 16.5] | 12.01 |
| **'1734'** | *Corynebacterium kutscheri* DSM 20755 | 13.1 | [10.4 - 16.4] | 24.7 | [22.4 - 27.2] | 13.5 | [11.1 - 16.3] | 7.14 |
| **'2937'** | *Corynebacterium vitaeruminis* DSM 20294 | 13.3 | [10.6 - 16.6] | 24.7 | [22.4 - 27.2] | 13.7 | [11.3 - 16.5] | 11.97 |
| **'CMCNS703'** | *Corynebacterium argentoratense* DSM 44202 | 13.2 | [10.5 - 16.5] | 24.7 | [22.4 - 27.2] | 13.6 | [11.2 - 16.4] | 5.17 |
| **'CHUV2995'** | *Corynebacterium argentoratense* DSM 44202 | 13.2 | [10.5 - 16.5] | 24.7 | [22.4 - 27.2] | 13.6 | [11.2 - 16.4] | 4.96 |
| **'FRC0043'** | *Corynebacterium singulare* DSM 44357 | 13.0 | [10.3 - 16.3] | 24.7 | [22.4 - 27.2] | 13.4 | [11.0 - 16.2] | 6.5 |
| **'CHUV2995'** | *Corynebacterium tuscaniense* CCUG 51321 | 13.3 | [10.6 - 16.6] | 24.6 | [22.3 - 27.1] | 13.7 | [11.3 - 16.5] | 5.48 |
| **'2937'** | *Corynebacterium tuscaniense* CCUG 51321 | 13.0 | [10.3 - 16.3] | 24.2 | [21.9 - 26.6] | 13.4 | [11.0 - 16.1] | 5.86 |
| **'CHUV2995'** | *Corynebacterium silvaticum* KL0182 | 14.4 | [11.5 - 17.7] | 23.8 | [21.5 - 26.3] | 14.6 | [12.2 - 17.5] | 0.51 |
| **'FRC0043'** | *Corynebacterium argentoratense* DSM 44202 | 13.2 | [10.5 - 16.5] | 23.7 | [21.4 - 26.1] | 13.5 | [11.2 - 16.3] | 5.28 |
| **'1137'** | *Corynebacterium ulcerans* NCTC 7910 | 14.2 | [11.4 - 17.5] | 23.4 | [21.1 - 25.9] | 14.5 | [12.0 - 17.3] | 0.3 |
| **'631'** | *Corynebacterium ulcerans* NCTC 7910 | 14.2 | [11.4 - 17.5] | 23.4 | [21.1 - 25.9] | 14.5 | [12.0 - 17.3] | 0.3 |
| **'4499'** | *Corynebacterium ulcerans* NCTC 7910 | 14.2 | [11.4 - 17.5] | 23.4 | [21.1 - 25.9] | 14.5 | [12.0 - 17.3] | 0.3 |

| Query | Subject | $d_0$ | C.I. $d_0$ | $d_4$ | C.I. $d_4$ | $d_6$ | C.I. $d_6$ | Diff. G+C Percent |
|---|---|---|---|---|---|---|---|---|
| **'1556'** | *Corynebacterium ulcerans* NCTC 7910 | 14.2 | [11.3 - 17.5] | 23.4 | [21.1 - 25.8] | 14.4 | [12.0 - 17.3] | 0.31 |
| **'5130'** | *Corynebacterium ulcerans* NCTC 7910 | 14.2 | [11.3 - 17.5] | 23.3 | [21.0 - 25.8] | 14.4 | [12.0 - 17.3] | 0.3 |
| **'4663'** | *Corynebacterium ulcerans* NCTC 7910 | 14.2 | [11.3 - 17.5] | 23.3 | [21.0 - 25.8] | 14.4 | [12.0 - 17.3] | 0.29 |
| **'5130'** | *Corynebacterium tuscaniense* CCUG 51321 | 13.0 | [10.3 - 16.3] | 23.3 | [21.0 - 25.7] | 13.4 | [11.0 - 16.1] | 5.81 |
| **'4766'** | *Corynebacterium ulcerans* NCTC 7910 | 14.2 | [11.4 - 17.6] | 23.3 | [21.0 - 25.8] | 14.5 | [12.0 - 17.3] | 0.2 |
| **'1734'** | *Corynebacterium ulcerans* NCTC 7910 | 14.2 | [11.3 - 17.5] | 23.3 | [21.1 - 25.8] | 14.5 | [12.0 - 17.3] | 0.29 |
| **'1556'** | *Corynebacterium tuscaniense* CCUG 51321 | 13.0 | [10.3 - 16.3] | 23.3 | [21.0 - 25.7] | 13.4 | [11.0 - 16.1] | 5.8 |
| **'631'** | *Corynebacterium tuscaniense* CCUG 51321 | 13.0 | [10.3 - 16.3] | 23.3 | [21.0 - 25.7] | 13.4 | [11.0 - 16.1] | 5.81 |
| **'1137'** | *Corynebacterium tuscaniense* CCUG 51321 | 13.0 | [10.3 - 16.3] | 23.3 | [21.0 - 25.7] | 13.4 | [11.0 - 16.1] | 5.81 |
| **'4663'** | *Corynebacterium tuscaniense* CCUG 51321 | 13.0 | [10.3 - 16.3] | 23.3 | [21.0 - 25.7] | 13.4 | [11.0 - 16.1] | 5.82 |
| **'1734'** | *Corynebacterium tuscaniense* CCUG 51321 | 13.0 | [10.3 - 16.3] | 23.3 | [21.0 - 25.7] | 13.4 | [11.0 - 16.1] | 5.82 |
| **'4499'** | *Corynebacterium tuscaniense* CCUG 51321 | 13.0 | [10.3 - 16.3] | 23.3 | [21.0 - 25.7] | 13.4 | [11.0 - 16.1] | 5.81 |
| **'4766'** | *Corynebacterium tuscaniense* CCUG 51321 | 13.0 | [10.3 - 16.3] | 23.2 | [20.9 - 25.7] | 13.4 | [11.0 - 16.1] | 5.91 |
| **'CHUV2995'** | *Corynebacterium ulcerans* NCTC 7910 | 14.2 | [11.3 - 17.5] | 23.1 | [20.8 - 25.6] | 14.4 | [12.0 - 17.3] | 0.63 |
| **'CMCNS703'** | *Corynebacterium silvaticum* KL0182 | 14.2 | [11.4 - 17.6] | 23.1 | [20.8 - 25.6] | 14.5 | [12.0 - 17.3] | 0.72 |
| **'CMCNS703'** | *Corynebacterium ulcerans* NCTC 7910 | 14.2 | [11.4 - 17.6] | 23.0 | [20.7 - 25.5] | 14.5 | [12.1 - 17.4] | 0.42 |
| **'2937'** | *Corynebacterium ulcerans* NCTC 7910 | 14.2 | [11.4 - 17.6] | 23.0 | [20.7 - 25.4] | 14.5 | [12.1 - 17.3] | 0.25 |
| **'FRC0043'** | *Corynebacterium silvaticum* KL0182 | 14.2 | [11.4 - 17.6] | 22.7 | [20.4 - 25.2] | 14.5 | [12.0 - 17.3] | 0.83 |
| **'FRC0043'** | *Corynebacterium ulcerans* NCTC 7910 | 14.2 | [11.4 - 17.6] | 22.6 | [20.3 - 25.0] | 14.5 | [12.0 - 17.3] | 0.31 |
| **'2937'** | *Corynebacterium silvaticum* KL0182 | 14.3 | [11.4 - 17.6] | 22.5 | [20.2 - 24.9] | 14.5 | [12.1 - 17.4] | 0.89 |
| **'FRC0043'** | *Corynebacterium hadale* NBT06-6 | 12.9 | [10.2 - 16.2] | 22.4 | [20.2 - 24.9] | 13.3 | [10.9 - 16.1] | 11.55 |
| **'631'** | *Corynebacterium silvaticum* KL0182 | 14.2 | [11.4 - 17.6] | 22.2 | [19.9 - 24.6] | 14.5 | [12.0 - 17.3] | 0.84 |
| **'1556'** | *Corynebacterium silvaticum* KL0182 | 14.2 | [11.4 - 17.6] | 22.2 | [19.9 - 24.6] | 14.5 | [12.0 - 17.3] | 0.83 |
| **'4766'** | *Corynebacterium silvaticum* KL0182 | 14.3 | [11.4 - 17.6] | 22.2 | [19.9 - 24.6] | 14.5 | [12.1 - 17.4] | 0.93 |
| **'4663'** | *Corynebacterium silvaticum* KL0182 | 14.2 | [11.4 - 17.6] | 22.1 | [19.8 - 24.6] | 14.5 | [12.0 - 17.3] | 0.84 |
| **'4499'** | *Corynebacterium silvaticum* KL0182 | 14.2 | [11.4 - 17.6] | 22.1 | [19.9 - 24.6] | 14.5 | [12.1 - 17.3] | 0.84 |
| **'1734'** | *Corynebacterium silvaticum* KL0182 | 14.2 | [11.4 - 17.6] | 22.1 | [19.8 - 24.6] | 14.5 | [12.0 - 17.3] | 0.84 |

| Query | Subject | $d_0$ | C.I. $d_0$ | $d_4$ | C.I. $d_4$ | $d_6$ | C.I. $d_6$ | Diff. G+C Percent |
|---|---|---|---|---|---|---|---|---|
| **'1137'** | *Corynebacterium silvaticum* KL0182 | 14.2 | [11.4 - 17.6] | 22.1 | [19.9 - 24.6] | 14.5 | [12.1 - 17.3] | 0.84 |
| **'5130'** | *Corynebacterium silvaticum* KL0182 | 14.2 | [11.4 - 17.6] | 22.1 | [19.9 - 24.6] | 14.5 | [12.0 - 17.3] | 0.83 |
| **'FRC0043'** | *Corynebacterium tuscaniense* CCUG 51321 | 12.9 | [10.2 - 16.2] | 21.9 | [19.6 - 24.3] | 13.3 | [11.0 - 16.1] | 5.8 |
| **'CMCNS703'** | *Corynebacterium pseudotuberculosis* DSM 20689 | 14.2 | [11.3 - 17.5] | 21.6 | [19.4 - 24.1] | 14.4 | [12.0 - 17.2] | 1.55 |
| **'CMCNS703'** | *Corynebacterium pseudotuberculosis* ATCC 19410 | 14.2 | [11.3 - 17.5] | 21.6 | [19.4 - 24.1] | 14.4 | [12.0 - 17.2] | 1.55 |
| **'CHUV2995'** | *Corynebacterium pseudotuberculosis* DSM 20689 | 14.1 | [11.3 - 17.5] | 21.5 | [19.3 - 23.9] | 14.4 | [11.9 - 17.2] | 1.76 |
| **'CHUV2995'** | *Corynebacterium pseudotuberculosis* ATCC 19410 | 14.1 | [11.3 - 17.5] | 21.5 | [19.3 - 23.9] | 14.4 | [11.9 - 17.2] | 1.76 |
| **'2937'** | *Corynebacterium pseudotuberculosis* ATCC 19410 | 14.2 | [11.4 - 17.5] | 21.3 | [19.0 - 23.7] | 14.4 | [12.0 - 17.3] | 1.38 |
| **'2937'** | *Corynebacterium pseudotuberculosis* DSM 20689 | 14.2 | [11.4 - 17.5] | 21.3 | [19.0 - 23.7] | 14.4 | [12.0 - 17.3] | 1.38 |
| **'FRC0043'** | *Corynebacterium pseudotuberculosis* ATCC 19410 | 14.2 | [11.4 - 17.5] | 21.1 | [18.8 - 23.5] | 14.4 | [12.0 - 17.2] | 1.44 |
| **'FRC0043'** | *Corynebacterium pseudotuberculosis* DSM 20689 | 14.2 | [11.4 - 17.5] | 21.1 | [18.8 - 23.5] | 14.4 | [12.0 - 17.2] | 1.44 |
| **'631'** | *Corynebacterium pseudotuberculosis* DSM 20689 | 14.3 | [11.5 - 17.7] | 20.9 | [18.6 - 23.3] | 14.6 | [12.1 - 17.4] | 1.42 |
| **'1556'** | *Corynebacterium pseudotuberculosis* DSM 20689 | 14.3 | [11.5 - 17.7] | 20.9 | [18.6 - 23.3] | 14.5 | [12.1 - 17.4] | 1.44 |
| **'4766'** | *Corynebacterium pseudotuberculosis* ATCC 19410 | 14.4 | [11.5 - 17.7] | 20.9 | [18.6 - 23.3] | 14.6 | [12.1 - 17.4] | 1.33 |
| **'4766'** | *Corynebacterium pseudotuberculosis* DSM 20689 | 14.4 | [11.5 - 17.7] | 20.9 | [18.6 - 23.3] | 14.6 | [12.1 - 17.4] | 1.33 |
| **'631'** | *Corynebacterium pseudotuberculosis* ATCC 19410 | 14.3 | [11.5 - 17.7] | 20.9 | [18.6 - 23.3] | 14.6 | [12.1 - 17.4] | 1.43 |
| **'4499'** | *Corynebacterium pseudotuberculosis* ATCC 19410 | 14.3 | [11.5 - 17.7] | 20.9 | [18.6 - 23.3] | 14.6 | [12.1 - 17.4] | 1.43 |
| **'1137'** | *Corynebacterium pseudotuberculosis* DSM 20689 | 14.3 | [11.5 - 17.7] | 20.9 | [18.6 - 23.3] | 14.6 | [12.1 - 17.4] | 1.42 |
| **'4499'** | *Corynebacterium pseudotuberculosis* DSM 20689 | 14.3 | [11.5 - 17.7] | 20.9 | [18.6 - 23.3] | 14.6 | [12.1 - 17.4] | 1.42 |
| **'1556'** | *Corynebacterium pseudotuberculosis* ATCC 19410 | 14.3 | [11.5 - 17.7] | 20.9 | [18.6 - 23.3] | 14.5 | [12.1 - 17.4] | 1.44 |
| **'1137'** | *Corynebacterium pseudotuberculosis* ATCC 19410 | 14.3 | [11.5 - 17.7] | 20.9 | [18.6 - 23.3] | 14.6 | [12.1 - 17.4] | 1.43 |

| Query | Subject | $d_0$ | C.I. $d_0$ | $d_4$ | C.I. $d_4$ | $d_6$ | C.I. $d_6$ | Diff. G+C Percent |
|---|---|---|---|---|---|---|---|---|
| **'4663'** | *Corynebacterium pseudotuberculosis* ATCC 19410 | 14.3 | [11.5 - 17.7] | 20.8 | [18.6 - 23.3] | 14.6 | [12.1 - 17.4] | 1.42 |
| **'4663'** | *Corynebacterium pseudotuberculosis* DSM 20689 | 14.3 | [11.5 - 17.7] | 20.8 | [18.6 - 23.3] | 14.6 | [12.1 - 17.4] | 1.42 |
| **'1734'** | *Corynebacterium pseudotuberculosis* DSM 20689 | 14.3 | [11.5 - 17.7] | 20.8 | [18.6 - 23.3] | 14.6 | [12.1 - 17.4] | 1.42 |
| **'1734'** | *Corynebacterium pseudotuberculosis* ATCC 19410 | 14.3 | [11.5 - 17.7] | 20.8 | [18.6 - 23.3] | 14.6 | [12.1 - 17.4] | 1.42 |
| **'5130'** | *Corynebacterium pseudotuberculosis* ATCC 19410 | 14.3 | [11.5 - 17.7] | 20.8 | [18.6 - 23.3] | 14.6 | [12.1 - 17.4] | 1.43 |
| **'5130'** | *Corynebacterium pseudotuberculosis* DSM 20689 | 14.3 | [11.5 - 17.7] | 20.8 | [18.6 - 23.3] | 14.6 | [12.1 - 17.4] | 1.43 |

#### Table 4: Strains in your dataset

Joint dataset of automatically determined closest type strains (if this mode was chosen), manually selected type strains (if selected accordingly) and the provided user strains, if provided (marked in yellow).

| Strain | Authority | Other deposits | Synonyms | Base pairs | Percent G+C | No. proteins | Goldstamp | Bioproject accession | Biosample accession | Assembly accession | IMG OID |
|---|---|---|---|---|---|---|---|---|---|---|---|
| *Corynebacterium vitaeruminis* DSM 20294 | (Bechdel et al. 1928) Lanéelle et al. 1980 | CCUG 28792; ATCC 10234; JCM 1323; IFO 12143; NBRC 12143; VKM B-1211; CIP 82.07; NCIB 9291; NCIMB 9291 | *Brevibacterium vitaeruminis*; *Corynebacterium vitaeruminis*; *Flavobacterium vitarumen* | 2931 780 | 65.5 | 2577 | Gp0023683 | PRJNA172966 | SAMN03081455 | GCA_000550805 | 2558860221 |
| *Corynebacterium phoceense* MC1 | Cresci et al. 2016 | DSM 100570; CSUR P1905 | *Corynebacterium phoceense* | 2772 735 | 63.2 | 2701 | | PRJNA224116 | SAMEA4059842 | GCF_900092335 | |
| *Corynebacterium diphtheriae* subsp. *lausannense* CHUV2995 | Tagini et al. 2019 | CCUG 72509; DSM 107520 | *Corynebacterium diphtheriae* subsp. *lausannense* | 3060 363 | 53.9 | 3145 | Gp0442955 | PRJEB24256 | SAMEA104679569 | GCA_900312965 | |
| *Corynebacterium belfantii* FRC0043 | Dazas et al. 2018 | DSM 105776; CIP 111412 | *Corynebacterium belfantii* | 2598 827 | 53.6 | 2557 | Gp0364753 | PRJEB22103 | SAMEA104208677 | GCA_900205605 | |
| *Corynebacterium silvaticum* KL0182 | Dangel et al. 2020 | LMG 31313; DSM 109166; CIP 111672 | *Corynebacterium silvaticum* | 2548 487 | 54.4 | 2017 | | PRJNA517029 | SAMN10039578 | GCA_004382825 | |
| *Corynebacterium tuscaniense* CCUG 51321 | Riegel et al. 2006 | DSM 45101; ATCC BAA-1141; JCM 15294; ISS-5309 | *Corynebacterium tuscaniense* | 2232 117 | 59.4 | 2073 | | PRJNA224116 | SAMN12771122 | GCF_008693065 | |

| Strain | Authority | Other deposits | Synonyms | Base pairs | Percent G+C | No. proteins | Goldstamp | Bioproject accession | Biosample accession | Assembly accession | IMG OID |
|---|---|---|---|---|---|---|---|---|---|---|---|
| *Corynebacterium rouxii* FRC0190 T | Badell et al. 2020 | DSM 110354; CIP 111752 | *Corynebacterium rouxii* | 2451 019 | 53.2 | 2365 | | PRJNA224116 | SAMEA5992727 | GCF_902702935 | |
| *Corynebacterium hadale* NBT06-6 | Wei et al. 2018 | MCCC 1K03347; DSM 105365 | *Corynebacterium hadale* | 2679 199 | 65.2 | 2362 | Gp0374689 | PRJNA396693 | SAMN07460149 | GCA_002273005 | |
| *Corynebacterium pseudotuberculosis* ATCC 19410 | (Buchanan 1911) Eberson 1918 emend. Nouioui et al. 2018 | CCUG 2806; DSM 20689; ATCC 19410; NCTC 3450; JCM 9389; IFO 15363; NBRC 15363; CIP 102968 | *Bacillus pseudotuberculosis*; *Corynebacterium pseudotuberculosis* | 2337 763 | 52.2 | 2146 | Gp0223239 | PRJNA382169 | SAMN06701041 | GCA_002155265 | |
| *Corynebacterium pseudotuberculosis* DSM 20689 | (Buchanan 1911) Eberson 1918 emend. Nouioui et al. 2018 | CCUG 2806; DSM 20689; ATCC 19410; NCTC 3450; JCM 9389; IFO 15363; NBRC 15363; CIP 102968 | *Bacillus pseudotuberculosis*; *Corynebacterium pseudotuberculosis* | 2338 546 | 52.2 | 2084 | Gp0220522 | PRJNA442833 | SAMN08778220 | GCA_003634885 | 2756170169 |
| *Corynebacterium ulcerans* NCTC 7910 | (ex Gilbert and Stewart 1927) Riegel et al. 1995 | CCUG 2708; DSM 46325; ATCC 51799; JCM 10387; CIP 106504 | *Corynebacterium ulcerans* | 2453 761 | 53.3 | 2178 | Gp0262745 | PRJEB6403 | SAMEA4504038 | GCA_900187135 | |
| *Corynebacterium striatum* ATCC 6940 | (Chester 1901) Eberson 1918 emend. Nouioui et al. 2018 | CCUG 27949; DSM 20668; NCTC 764; JCM 9390; IFO 15291; NBRC 15291; CIP 81.15 | *Bacterium striatum*; *Corynebacterium striatum* | 2717 381 | 59.4 | 2677 | Gp0003467 | PRJNA31449 | SAMN00001507 | GCA_000159135 | 643886057 |

| Strain | Authority | Other deposits | Synonyms | Base pairs | Percent G+C | No. proteins | Goldstamp | Bioproject accession | Biosample accession | Assembly accession | IMG OID |
|---|---|---|---|---|---|---|---|---|---|---|---|
| *Corynebacterium kutscheri* DSM 20755 | (Migula 1900) Bergey et al. 1925 emend. Nouioui et al. 2018 | CCUG 27535; ATCC 15677; NCTC 11138; JCM 9385; IFO 15288; NBRC 15288; CIP 103423 | *Bacterium kutscheri*; *Corynebacterium kutscheri* | 2354 065 | 46.5 | 2047 | Gp0110293 | PRJNA276037 | SAMN03365283 | GCA_000980835 | |
| *Corynebacterium singulare* DSM 44357 | Riegel et al. 1997 emend. Nouioui et al. 2018 | CCUG 37330; JCM 10385; IFO 16162; NBRC 16162; CIP 105491; IBS B52218 | *Corynebacterium singulare* | 2830 499 | 60.1 | 2561 | Gp0109683 | PRJNA246651 | SAMN03177398 | GCA_000833575 | |
| *Corynebacterium diphtheriae* NCTC 11397 | (Kruse 1886) Lehmann and Neumann 1896 emend. Nouioui et al. 2018 | DSM 44123; ATCC 27010; CIP 100721 | *Bacillus diphtheriae*; *Corynebacterium diphtheriae*; *Corynebacterium diphtheriae* subsp. *diphtheriae* | 2463 666 | 53.5 | 2337 | Gp0132011 | PRJEB6403 | SAMEA2517360 | GCA_001457455 | |
| *Corynebacterium argentoratense* DSM 44202 | Riegel et al. 1995 emend. Nouioui et al. 2018 | CCUG 34893; ATCC 51927; JCM 10392; CIP 104296; IBS B10697 | *Corynebacterium argentoratense* | 2031 862 | 58.9 | 1875 | Gp0044215 | PRJNA209048 | SAMN02603032 | GCA_000590555 | 2554235426 |
| 631 | | | | 2397 667 | 53.6 | 2275 | | | | | |
| 1137 | | | | 2398 650 | 53.6 | 2276 | | | | | |
| 1556 | | | | 2445 187 | 53.6 | 2351 | | | | | |
| 1734 | | | | 2394 567 | 53.6 | 2263 | | | | | |
| 2937 | | | | 2466 271 | 53.6 | 2377 | | | | | |

| Strain | Authority | Other deposits | Synonyms | Base pairs | Percent G+C | No. proteins | Goldstamp | Bioproject accession | Biosample accession | Assembly accession | IMG OID |
|---|---|---|---|---|---|---|---|---|---|---|---|
| 4499 | | | | 2397 617 | 53.6 | 2273 | | | | | |
| 4663 | | | | 2393 381 | 53.6 | 2268 | | | | | |
| 4766 | | | | 2352 296 | 53.5 | 2221 | | | | | |
| 5130 | | | | 2392 741 | 53.6 | 2267 | | | | | |
| CHUV2995 | | | | 3060 363 | 53.9 | 3176 | | | | | |
| CMCNS703 | | | | 2892 379 | 53.7 | 3007 | | | | | |
| FRC0043 | | | | 2609 417 | 53.6 | 2663 | | | | | |

The genome sequence data were uploaded to the Type (Strain) Genome Server (TYGS), a free bioinformatics platform available under https://tygs.dsmz.de, for a whole genome-based taxonomic analysis [1]. The results were provided by the TYGS on 2020-06-19. In brief, the TYGS analysis was subdivided into the following steps:

#### Determination of closely related type strains

Determination of closest type strain genomes was done in two complementary ways: First, all user genomes were compared against all type strain genomes available in the TYGS database via the MASH algorithm, a fast approximation of intergenomic relatedness [2], and, the ten type strains with the smallest MASH distances chosen per user genome. Second, an additional set of ten closely related type strains was determined via the 16S rDNA gene sequences. These were extracted from the user genomes using RNAmmer [3] and each sequence was subsequently BLASTed [4] against the 16S rDNA gene sequence of each of the currently 11861 type strains available in the TYGS database. This was used as a proxy to find the best 50 matching type strains (according to the bitscore) for each user genome and to subsequently calculate precise distances using the Genome BLAST Distance Phylogeny approach (GBDP) under the algorithm 'coverage' and distance formula $d_5$ [5]. These distances were finally used to determine the 10 closest type strain genomes for each of the user genomes.

#### Pairwise comparison of genome sequences

All pairwise comparisons among the set of genomes were conducted using GBDP and accurate intergenomic distances inferred under the algorithm 'trimming' and distance formula $d_5$ [5]. 100 distance replicates were calculated each. Digital DDH values and confidence intervals were calculated using the recommended settings of the GGDC 2.1 [5].

#### Phylogenetic inference

The resulting intergenomic distances were used to infer a balanced minimum evolution tree with branch support via FASTME 2.1.4 including SPR postprocessing [6]. Branch support was inferred from 100 pseudo-bootstrap replicates each. The trees were rooted at the midpoint [7] and visualized with PhyD3 [8].

#### Type-based species and subspecies clustering

The type-based species clustering using a 70% dDDH radius around each of the 16 type strains was done as previously described [1]. The resulting groups are shown in Table 1 and 4. Subspecies clustering was done using a 79% dDDH threshold as previously introduced [9].

### Results

#### Type-based species and subspecies clustering

The resulting species and subspecies clusters are listed in Table 4, whereas the taxonomic identification of the query strains is found in Table 1. Briefly, the clustering yielded 14 species clusters and the provided query strains were assigned to 2 of these. Moreover, user strains were located in 2 of 14 subspecies clusters.

#### Figure caption genome tree

**Figure 1.** Tree inferred with FastME 2.1.6.1 [6] from GBDP distances calculated from genome sequences. The branch lengths are scaled in terms of GBDP distance formula $d_5$. The numbers above branches are GBDP pseudo-bootstrap support values > 60 % from 100 replications, with an average branch support of 37.1 %. The tree was rooted at the midpoint [7].

#### Figure caption SSU tree

**Figure 2.** Tree inferred with FastME 2.1.6.1 [6] from GBDP distances calculated from 16S rDNA gene sequences. The branch lengths are scaled in terms of GBDP distance formula $d_5$. The numbers above branches are GBDP pseudo-bootstrap support values > 60 % from 100 replications, with an average branch support of 51.6 %. The tree was rooted at the midpoint [7].
