## Supplementary File 3 for "Taxonomic classification of strain PO100/5 shows a broader geographic distribution and genetic markers of the recently described *Corynebacterium silvaticum*": Cul_assembled_p2_report.pdf

TYPE (STRAIN) GENOME SERVER

### JOB SUMMARY

| | |
|---|---|
| PRINT DATE: | 2020-06-11 22:42:34 +0200 |
| JOB ID: | 3f1a8641-39bd-4f4c-b3ac-0c6455711329 |
| RESULT PAGE: | https://tygs.dsmz.de/user_results/show?guid=3f1a8641-39bd-4f4c-b3ac-0c6455711329 |

#### Table 1: Phylogenies

**Publication-ready versions** of both the genome-scale GBDP tree and the 16S rRNA gene sequence tree can be customized and exported either in SVG (vector graphic) or PNG format from within the phylogeny viewers in your TYGS result page. For publications the **SVG format is recommended** because it is lossless, always keeps its high resolution and can also be easily converted to other popular formats such as PDF or EPS. Please follow the link provided above!

#### Table 2: Identification

The below list contains the result of the TYGS species identification routine.

Explanation of remarks that might occur in the below table:

**remark [R1]:** The TYGS type strain database is automatically updated on an almost daily basis. However, if a particular type strain genome is not available in the TYGS database, this can have several reasons which are detailed in the FAQ. You can request an extended 16S rRNA gene analysis via the 16S tree viewer found in your result page to detect **not yet genome-sequenced** type strains relevant for your study.

**remark [R2]:** > 70% dDDH value (formula $d_4$) and (almost) minimal dDDH values for gene-content formulae $d_0$ and $d_6$ indicate a potentially unreliable identification result and should thus be checked via the 16S rRNA gene sequence similarity. Such strong deviations can, in principle, be caused by sequence contamination.

**remark [R3]:** G+C content difference of > 1 % indicates a potentially unreliable identification result because within species G+C content varies no more than 1 %, if computed from genome sequences (PMID: 24505073).

| Strain | Conclusion | Identification result | Remark |
|---|---|---|---|
| **'KL0475'** | belongs to known species | *Corynebacterium ulcerans* | |
| **'KL0483'** | belongs to known species | *Corynebacterium ulcerans* | |
| **'KL0497'** | belongs to known species | *Corynebacterium ulcerans* | |
| **'KL0501'** | belongs to known species | *Corynebacterium ulcerans* | |
| **'KL0515'** | belongs to known species | *Corynebacterium ulcerans* | |
| **'KL0540'** | belongs to known species | *Corynebacterium ulcerans* | |
| **'KL0541'** | belongs to known species | *Corynebacterium ulcerans* | |
| **'KL0547'** | belongs to known species | *Corynebacterium ulcerans* | |
| **'KL0556'** | belongs to known species | *Corynebacterium ulcerans* | |
| **'KL0785'** | belongs to known species | *Corynebacterium ulcerans* | |
| **'KL0796'** | belongs to known species | *Corynebacterium ulcerans* | |
| **'KL0818'** | belongs to known species | *Corynebacterium ulcerans* | |
| **'KL0825'** | belongs to known species | *Corynebacterium ulcerans* | |
| **'KL0832'** | belongs to known species | *Corynebacterium ulcerans* | |
| **'KL0840'** | belongs to known species | *Corynebacterium ulcerans* | |

| Strain | Conclusion | Identification result | Remark |
|---|---|---|---|
| **'KL0846'** | belongs to known species | *Corynebacterium ulcerans* | |
| **'KL0853'** | belongs to known species | *Corynebacterium ulcerans* | |
| **'KL0867'** | belongs to known species | *Corynebacterium ulcerans* | |
| **'KL0870'** | belongs to known species | *Corynebacterium ulcerans* | |
| **'KL0876'** | belongs to known species | *Corynebacterium ulcerans* | |

#### Table 3: Pairwise comparisons of user genomes vs. type-strain genomes

The following table contains the pairwise dDDH values between your user genomes and the selected type-strain genomes. The dDDH values are provided along with their confidence intervals (C.I.) for the three different GBDP formulas:

- formula $d_0$ (a.k.a. GGDC formula 1): length of all HSPs divided by total genome length
- formula $d_4$ (a.k.a. GGDC formula 2): sum of all identities found in HSPs divided by overall HSP length
- formula $d_6$ (a.k.a. GGDC formula 3): sum of all identities found in HSPs divided by total genome length

**Note:** Formula $d_4$ is independent of genome length and is thus robust against the use of incomplete draft genomes. For other reasons for preferring formula $d_4$, see the FAQ.

| Query | Subject | $d_0$ | C.I. $d_0$ | $d_4$ | C.I. $d_4$ | $d_6$ | C.I. $d_6$ | Diff. G+C Percent |
|---|---|---|---|---|---|---|---|---|
| **'KL0785'** | **'KL0818'** | 100.0 | [100.0 - 100.0] | 100.0 | [100.0 - 100.0] | 100.0 | [100.0 - 100.0] | 0.01 |
| **'KL0515'** | **'KL0785'** | 99.4 | [98.8 - 99.7] | 99.9 | [99.8 - 100.0] | 99.7 | [99.5 - 99.9] | 0.04 |
| **'KL0515'** | **'KL0818'** | 99.3 | [98.7 - 99.6] | 99.9 | [99.8 - 100.0] | 99.7 | [99.4 - 99.8] | 0.03 |
| **'KL0483'** | **'KL0846'** | 98.2 | [97.0 - 98.9] | 99.6 | [99.4 - 99.8] | 99.1 | [98.6 - 99.5] | 0.08 |
| **'KL0825'** | **'KL0846'** | 99.6 | [99.1 - 99.8] | 99.6 | [99.4 - 99.8] | 99.8 | [99.6 - 99.9] | 0.0 |
| **'KL0475'** | **'KL0846'** | 95.2 | [93.0 - 96.8] | 99.6 | [99.3 - 99.8] | 97.4 | [96.1 - 98.3] | 0.01 |
| **'KL0483'** | **'KL0832'** | 98.4 | [97.3 - 99.1] | 99.6 | [99.3 - 99.7] | 99.2 | [98.7 - 99.6] | 0.09 |
| **'KL0825'** | **'KL0840'** | 99.6 | [99.3 - 99.8] | 99.6 | [99.3 - 99.8] | 99.8 | [99.7 - 99.9] | 0.0 |
| **'KL0840'** | **'KL0846'** | 100.0 | [99.9 - 100.0] | 99.6 | [99.3 - 99.8] | 100.0 | [99.9 - 100.0] | 0.0 |
| **'KL0540'** | **'KL0547'** | 97.9 | [96.6 - 98.8] | 99.6 | [99.3 - 99.7] | 99.0 | [98.3 - 99.4] | 0.02 |
| **'KL0475'** | **'KL0832'** | 95.5 | [93.4 - 97.0] | 99.5 | [99.2 - 99.7] | 97.6 | [96.4 - 98.4] | 0.01 |
| **'KL0475'** | **'KL0840'** | 95.6 | [93.5 - 97.1] | 99.5 | [99.2 - 99.7] | 97.7 | [96.4 - 98.5] | 0.01 |
| **'KL0547'** | **'KL0867'** | 99.7 | [99.4 - 99.9] | 99.5 | [99.3 - 99.7] | 99.9 | [99.7 - 99.9] | 0.0 |
| **'KL0796'** | **'KL0846'** | 97.7 | [96.2 - 98.6] | 99.5 | [99.2 - 99.7] | 98.8 | [98.1 - 99.3] | 0.04 |
| **'KL0483'** | **'KL0840'** | 98.4 | [97.2 - 99.0] | 99.5 | [99.2 - 99.7] | 99.2 | [98.6 - 99.5] | 0.08 |
| **'KL0796'** | **'KL0832'** | 97.9 | [96.5 - 98.7] | 99.5 | [99.2 - 99.7] | 98.9 | [98.3 - 99.4] | 0.04 |
| **'KL0540'** | **'KL0867'** | 98.7 | [97.7 - 99.2] | 99.5 | [99.2 - 99.7] | 99.4 | [98.9 - 99.6] | 0.02 |
| **'KL0832'** | **'KL0846'** | 100.0 | [99.9 - 100.0] | 99.5 | [99.2 - 99.7] | 100.0 | [100.0 - 100.0] | 0.01 |
| **'KL0825'** | **'KL0832'** | 99.7 | [99.3 - 99.8] | 99.5 | [99.2 - 99.7] | 99.9 | [99.7 - 99.9] | 0.0 |
| **'KL0796'** | **'KL0840'** | 97.8 | [96.4 - 98.7] | 99.5 | [99.3 - 99.7] | 98.9 | [98.2 - 99.3] | 0.04 |
| **'KL0832'** | **'KL0840'** | 100.0 | [100.0 - 100.0] | 99.5 | [99.2 - 99.7] | 100.0 | [100.0 - 100.0] | 0.0 |
| **'KL0483'** | **'KL0825'** | 98.2 | [96.9 - 98.9] | 99.4 | [99.0 - 99.6] | 99.1 | [98.5 - 99.5] | 0.08 |

| Query | Subject | $d_0$ | C.I. $d_0$ | $d_4$ | C.I. $d_4$ | $d_6$ | C.I. $d_6$ | Diff. G+C Percent |
|---|---|---|---|---|---|---|---|---|
| **'KL0796'** | **'KL0825'** | 97.9 | [96.5 - 98.7] | 99.4 | [99.0 - 99.6] | 98.9 | [98.2 - 99.4] | 0.04 |
| **'KL0475'** | **'KL0796'** | 97.0 | [95.3 - 98.1] | 99.3 | [99.0 - 99.6] | 98.5 | [97.6 - 99.0] | 0.03 |
| **'KL0475'** | **'KL0825'** | 96.4 | [94.5 - 97.6] | 99.2 | [98.7 - 99.5] | 98.0 | [97.0 - 98.7] | 0.01 |
| **'KL0483'** | **'KL0796'** | 98.9 | [98.0 - 99.4] | 98.9 | [98.3 - 99.3] | 99.4 | [99.0 - 99.7] | 0.05 |
| **'KL0475'** | **'KL0483'** | 96.3 | [94.4 - 97.6] | 98.8 | [98.1 - 99.2] | 98.0 | [96.9 - 98.7] | 0.08 |
| **'KL0541'** | **'KL0867'** | 97.3 | [95.7 - 98.3] | 98.5 | [97.8 - 99.0] | 98.5 | [97.7 - 99.1] | 0.03 |
| **'KL0541'** | **'KL0547'** | 97.8 | [96.5 - 98.7] | 98.5 | [97.7 - 99.0] | 98.8 | [98.1 - 99.3] | 0.02 |
| **'KL0540'** | **'KL0541'** | 97.0 | [95.3 - 98.1] | 98.4 | [97.6 - 98.9] | 98.3 | [97.4 - 98.9] | 0.01 |
| **'KL0541'** | **'KL0840'** | 95.4 | [93.2 - 96.9] | 97.0 | [95.9 - 97.9] | 97.2 | [95.9 - 98.1] | 0.05 |
| **'KL0541'** | **'KL0846'** | 95.2 | [93.0 - 96.8] | 97.0 | [95.9 - 97.9] | 97.1 | [95.7 - 98.0] | 0.05 |
| **'KL0846'** | **'KL0867'** | 98.4 | [97.3 - 99.1] | 97.0 | [95.8 - 97.8] | 99.0 | [98.4 - 99.4] | 0.02 |
| **'KL0541'** | **'KL0832'** | 95.5 | [93.3 - 97.0] | 97.0 | [95.9 - 97.9] | 97.3 | [96.0 - 98.2] | 0.06 |
| **'KL0840'** | **'KL0867'** | 98.5 | [97.5 - 99.2] | 96.9 | [95.7 - 97.8] | 99.1 | [98.5 - 99.5] | 0.03 |
| **'KL0832'** | **'KL0867'** | 98.6 | [97.6 - 99.2] | 96.9 | [95.7 - 97.8] | 99.2 | [98.6 - 99.5] | 0.03 |
| **'KL0547'** | **'KL0846'** | 97.7 | [96.2 - 98.6] | 96.9 | [95.7 - 97.7] | 98.6 | [97.8 - 99.1] | 0.03 |
| **'KL0547'** | **'KL0832'** | 97.9 | [96.6 - 98.7] | 96.8 | [95.6 - 97.7] | 98.7 | [98.0 - 99.2] | 0.03 |
| **'KL0547'** | **'KL0840'** | 97.8 | [96.5 - 98.7] | 96.8 | [95.6 - 97.7] | 98.7 | [97.9 - 99.2] | 0.03 |
| **'KL0540'** | **'KL0846'** | 99.3 | [98.7 - 99.6] | 96.8 | [95.6 - 97.7] | 99.6 | [99.2 - 99.8] | 0.04 |
| **'KL0483'** | **'KL0867'** | 98.2 | [96.9 - 98.9] | 96.7 | [95.5 - 97.6] | 98.9 | [98.2 - 99.3] | 0.06 |
| **'KL0825'** | **'KL0867'** | 98.3 | [97.1 - 99.0] | 96.7 | [95.4 - 97.6] | 98.9 | [98.3 - 99.4] | 0.03 |
| **'KL0541'** | **'KL0825'** | 95.5 | [93.3 - 97.0] | 96.7 | [95.4 - 97.6] | 97.2 | [95.9 - 98.2] | 0.05 |
| **'KL0483'** | **'KL0547'** | 97.4 | [95.9 - 98.4] | 96.7 | [95.5 - 97.6] | 98.4 | [97.5 - 99.0] | 0.05 |
| **'KL0497'** | **'KL0547'** | 98.4 | [97.3 - 99.1] | 96.6 | [95.3 - 97.5] | 99.0 | [98.4 - 99.4] | 0.04 |
| **'KL0547'** | **'KL0825'** | 97.5 | [96.0 - 98.5] | 96.6 | [95.3 - 97.5] | 98.5 | [97.6 - 99.1] | 0.03 |
| **'KL0540'** | **'KL0840'** | 99.4 | [98.8 - 99.7] | 96.6 | [95.3 - 97.5] | 99.6 | [99.3 - 99.8] | 0.05 |
| **'KL0547'** | **'KL0796'** | 97.1 | [95.5 - 98.2] | 96.6 | [95.4 - 97.5] | 98.3 | [97.3 - 98.9] | 0.01 |
| **'KL0483'** | **'KL0540'** | 98.6 | [97.6 - 99.2] | 96.6 | [95.4 - 97.5] | 99.1 | [98.5 - 99.5] | 0.04 |

| Query | Subject | $d_0$ | C.I. $d_0$ | $d_4$ | C.I. $d_4$ | $d_6$ | C.I. $d_6$ | Diff. G+C Percent |
|---|---|---|---|---|---|---|---|---|
| **'KL0497'** | **'KL0867'** | 98.9 | [98.0 - 99.4] | 96.5 | [95.3 - 97.5] | 99.3 | [98.8 - 99.6] | 0.03 |
| **'KL0796'** | **'KL0867'** | 97.7 | [96.3 - 98.6] | 96.5 | [95.3 - 97.5] | 98.6 | [97.7 - 99.1] | 0.01 |
| **'KL0540'** | **'KL0832'** | 99.5 | [98.9 - 99.7] | 96.5 | [95.2 - 97.5] | 99.6 | [99.3 - 99.8] | 0.05 |
| **'KL0540'** | **'KL0825'** | 98.8 | [97.9 - 99.3] | 96.4 | [95.1 - 97.4] | 99.2 | [98.7 - 99.6] | 0.05 |
| **'KL0497'** | **'KL0540'** | 98.8 | [97.9 - 99.3] | 96.4 | [95.1 - 97.4] | 99.2 | [98.7 - 99.6] | 0.06 |
| **'KL0475'** | **'KL0867'** | 95.6 | [93.5 - 97.1] | 96.3 | [95.0 - 97.3] | 97.3 | [95.9 - 98.2] | 0.02 |
| **'KL0501'** | **'KL0540'** | 99.1 | [98.4 - 99.5] | 96.3 | [95.0 - 97.3] | 99.4 | [99.0 - 99.7] | 0.05 |
| **'KL0540'** | **'KL0796'** | 97.4 | [95.9 - 98.4] | 96.3 | [94.9 - 97.3] | 98.4 | [97.5 - 99.0] | 0.01 |
| **'KL0475'** | **'KL0547'** | 94.9 | [92.5 - 96.5] | 96.3 | [95.0 - 97.3] | 96.8 | [95.3 - 97.8] | 0.02 |
| **'KL0501'** | **'KL0867'** | 98.4 | [97.3 - 99.1] | 96.3 | [94.9 - 97.3] | 99.0 | [98.3 - 99.4] | 0.03 |
| **'KL0497'** | **'KL0541'** | 96.2 | [94.2 - 97.5] | 96.2 | [94.8 - 97.2] | 97.6 | [96.4 - 98.4] | 0.06 |
| **'KL0501'** | **'KL0547'** | 97.7 | [96.2 - 98.6] | 96.2 | [94.9 - 97.2] | 98.5 | [97.7 - 99.1] | 0.03 |
| **'KL0475'** | **'KL0541'** | 94.4 | [91.9 - 96.1] | 96.2 | [94.8 - 97.2] | 96.4 | [94.9 - 97.5] | 0.04 |
| **'KL0541'** | **'KL0796'** | 98.7 | [97.8 - 99.3] | 96.2 | [94.9 - 97.2] | 99.2 | [98.6 - 99.5] | 0.01 |
| **'KL0475'** | **'KL0540'** | 94.1 | [91.6 - 95.9] | 96.0 | [94.6 - 97.1] | 96.3 | [94.6 - 97.4] | 0.04 |
| **'KL0483'** | **'KL0541'** | 97.1 | [95.4 - 98.1] | 95.7 | [94.3 - 96.8] | 98.1 | [97.1 - 98.8] | 0.03 |
| **'KL0497'** | **'KL0796'** | 98.2 | [96.9 - 98.9] | 95.6 | [94.1 - 96.7] | 98.8 | [98.0 - 99.3] | 0.05 |
| **'KL0497'** | **'KL0840'** | 99.6 | [99.1 - 99.8] | 95.5 | [94.0 - 96.6] | 99.6 | [99.3 - 99.8] | 0.01 |
| **'KL0497'** | **'KL0846'** | 99.5 | [99.0 - 99.7] | 95.4 | [93.9 - 96.6] | 99.6 | [99.2 - 99.8] | 0.01 |
| **'KL0497'** | **'KL0825'** | 98.9 | [98.1 - 99.4] | 95.4 | [93.9 - 96.6] | 99.2 | [98.7 - 99.6] | 0.01 |
| **'KL0497'** | **'KL0832'** | 99.6 | [99.2 - 99.8] | 95.4 | [93.9 - 96.5] | 99.7 | [99.4 - 99.8] | 0.0 |
| **'KL0501'** | **'KL0541'** | 95.2 | [93.0 - 96.8] | 95.4 | [93.9 - 96.6] | 96.9 | [95.5 - 97.9] | 0.06 |
| **'KL0483'** | **'KL0497'** | 97.4 | [95.8 - 98.4] | 95.4 | [93.8 - 96.5] | 98.3 | [97.3 - 98.9] | 0.09 |
| **'KL0501'** | **'KL0846'** | 99.6 | [99.1 - 99.8] | 95.3 | [93.8 - 96.5] | 99.6 | [99.3 - 99.8] | 0.01 |
| **'KL0475'** | **'KL0497'** | 95.9 | [93.9 - 97.3] | 95.3 | [93.7 - 96.4] | 97.4 | [96.1 - 98.2] | 0.02 |
| **'KL0501'** | **'KL0825'** | 99.1 | [98.3 - 99.5] | 95.2 | [93.6 - 96.4] | 99.3 | [98.8 - 99.6] | 0.0 |
| **'KL0501'** | **'KL0796'** | 96.8 | [95.0 - 98.0] | 95.2 | [93.6 - 96.4] | 97.9 | [96.8 - 98.6] | 0.04 |

| Query | Subject | $d_0$ | C.I. $d_0$ | $d_4$ | C.I. $d_4$ | $d_6$ | C.I. $d_6$ | Diff. G+C Percent |
|---|---|---|---|---|---|---|---|---|
| **'KL0501'** | **'KL0832'** | 99.7 | [99.4 - 99.9] | 95.2 | [93.7 - 96.4] | 99.7 | [99.5 - 99.8] | 0.0 |
| **'KL0501'** | **'KL0840'** | 99.7 | [99.3 - 99.8] | 95.2 | [93.7 - 96.4] | 99.7 | [99.4 - 99.8] | 0.0 |
| **'KL0483'** | **'KL0501'** | 97.5 | [95.9 - 98.4] | 95.2 | [93.7 - 96.4] | 98.3 | [97.4 - 98.9] | 0.09 |
| **'KL0475'** | **'KL0501'** | 94.2 | [91.7 - 96.0] | 94.9 | [93.3 - 96.2] | 96.2 | [94.5 - 97.3] | 0.01 |
| **'KL0497'** | **'KL0501'** | 99.1 | [98.4 - 99.5] | 94.3 | [92.6 - 95.6] | 99.3 | [98.8 - 99.6] | 0.0 |
| **'KL0853'** | **'KL0876'** | 96.9 | [95.2 - 98.0] | 93.1 | [91.2 - 94.7] | 97.8 | [96.6 - 98.6] | 0.13 |
| **'KL0870'** | **'KL0876'** | 96.2 | [94.2 - 97.5] | 91.9 | [89.8 - 93.6] | 97.2 | [95.9 - 98.1] | 0.09 |
| **'KL0515'** | **'KL0853'** | 96.5 | [94.7 - 97.8] | 90.9 | [88.7 - 92.7] | 97.3 | [96.0 - 98.2] | 0.08 |
| **'KL0853'** | **'KL0870'** | 96.5 | [94.7 - 97.8] | 90.6 | [88.3 - 92.4] | 97.3 | [96.0 - 98.2] | 0.04 |
| **'KL0497'** | **'KL0870'** | 96.5 | [94.6 - 97.7] | 90.6 | [88.3 - 92.4] | 97.3 | [96.0 - 98.2] | 0.06 |
| **'KL0785'** | **'KL0853'** | 97.6 | [96.1 - 98.5] | 90.4 | [88.1 - 92.2] | 98.0 | [96.9 - 98.7] | 0.04 |
| **'KL0818'** | **'KL0853'** | 97.6 | [96.1 - 98.5] | 90.3 | [88.1 - 92.2] | 98.0 | [96.9 - 98.7] | 0.05 |
| **'KL0497'** | **'KL0876'** | 98.0 | [96.7 - 98.8] | 89.7 | [87.4 - 91.7] | 98.3 | [97.3 - 98.9] | 0.03 |
| **'KL0846'** | **'KL0870'** | 96.5 | [94.6 - 97.7] | 89.4 | [87.0 - 91.4] | 97.2 | [95.8 - 98.1] | 0.05 |
| **'KL0475'** | **'KL0870'** | 94.9 | [92.6 - 96.5] | 89.4 | [87.0 - 91.4] | 96.1 | [94.4 - 97.2] | 0.05 |
| **'KL0840'** | **'KL0870'** | 96.6 | [94.8 - 97.8] | 89.3 | [86.9 - 91.3] | 97.3 | [95.9 - 98.2] | 0.06 |
| **'KL0832'** | **'KL0870'** | 96.8 | [95.0 - 97.9] | 89.3 | [86.9 - 91.3] | 97.3 | [96.0 - 98.2] | 0.06 |
| **'KL0825'** | **'KL0870'** | 97.8 | [96.5 - 98.7] | 89.2 | [86.8 - 91.2] | 98.1 | [97.0 - 98.8] | 0.06 |
| **'KL0846'** | **'KL0876'** | 98.2 | [96.9 - 98.9] | 89.1 | [86.7 - 91.1] | 98.3 | [97.3 - 98.9] | 0.04 |
| **'KL0818'** | **'KL0876'** | 97.1 | [95.4 - 98.2] | 89.1 | [86.7 - 91.1] | 97.6 | [96.3 - 98.4] | 0.08 |
| **'KL0501'** | **'KL0870'** | 96.6 | [94.8 - 97.8] | 89.1 | [86.7 - 91.1] | 97.2 | [95.9 - 98.1] | 0.06 |
| **'KL0785'** | *Corynebacterium ulcerans* NCTC 7910 | 97.0 | [95.3 - 98.1] | 89.1 | [86.7 - 91.2] | 97.5 | [96.3 - 98.3] | 0.05 |
| **'KL0785'** | **'KL0876'** | 97.1 | [95.4 - 98.2] | 89.1 | [86.7 - 91.1] | 97.6 | [96.3 - 98.4] | 0.09 |
| **'KL0515'** | *Corynebacterium ulcerans* NCTC 7910 | 98.0 | [96.8 - 98.8] | 89.1 | [86.7 - 91.1] | 98.2 | [97.2 - 98.9] | 0.01 |
| **'KL0475'** | **'KL0876'** | 91.5 | [88.4 - 93.8] | 89.1 | [86.7 - 91.1] | 93.5 | [91.3 - 95.2] | 0.05 |
| **'KL0818'** | *Corynebacterium ulcerans* NCTC 7910 | 97.0 | [95.3 - 98.1] | 89.1 | [86.7 - 91.1] | 97.5 | [96.3 - 98.3] | 0.04 |
| **'KL0832'** | **'KL0876'** | 98.4 | [97.3 - 99.1] | 89.0 | [86.6 - 91.0] | 98.4 | [97.5 - 99.0] | 0.03 |

| Query | Subject | $d_0$ | C.I. $d_0$ | $d_4$ | C.I. $d_4$ | $d_6$ | C.I. $d_6$ | Diff. G+C Percent |
|---|---|---|---|---|---|---|---|---|
| **'KL0796'** | **'KL0876'** | 94.5 | [92.1 - 96.2] | 89.0 | [86.6 - 91.1] | 95.7 | [94.0 - 97.0] | 0.08 |
| **'KL0515'** | **'KL0876'** | 98.1 | [96.8 - 98.9] | 89.0 | [86.6 - 91.1] | 98.3 | [97.3 - 98.9] | 0.05 |
| **'KL0825'** | **'KL0876'** | 97.3 | [95.8 - 98.3] | 89.0 | [86.6 - 91.0] | 97.7 | [96.5 - 98.5] | 0.04 |
| **'KL0483'** | **'KL0876'** | 95.3 | [93.1 - 96.8] | 89.0 | [86.6 - 91.0] | 96.3 | [94.7 - 97.4] | 0.12 |
| **'KL0840'** | **'KL0876'** | 98.3 | [97.1 - 99.0] | 89.0 | [86.6 - 91.0] | 98.4 | [97.5 - 99.0] | 0.04 |
| **'KL0541'** | *Corynebacterium ulcerans* NCTC 7910 | 93.3 | [90.6 - 95.2] | 88.9 | [86.5 - 91.0] | 94.8 | [92.9 - 96.3] | 0.05 |
| **'KL0796'** | **'KL0870'** | 97.2 | [95.6 - 98.3] | 88.9 | [86.5 - 90.9] | 97.6 | [96.4 - 98.4] | 0.02 |
| **'KL0867'** | *Corynebacterium ulcerans* NCTC 7910 | 96.8 | [95.0 - 97.9] | 88.9 | [86.5 - 91.0] | 97.3 | [96.0 - 98.2] | 0.03 |
| **'KL0547'** | *Corynebacterium ulcerans* NCTC 7910 | 95.9 | [93.8 - 97.3] | 88.9 | [86.5 - 91.0] | 96.7 | [95.2 - 97.7] | 0.03 |
| **'KL0547'** | **'KL0876'** | 95.2 | [93.0 - 96.8] | 88.8 | [86.3 - 90.8] | 96.2 | [94.6 - 97.4] | 0.07 |
| **'KL0541'** | **'KL0876'** | 92.6 | [89.7 - 94.7] | 88.8 | [86.4 - 90.9] | 94.3 | [92.2 - 95.8] | 0.09 |
| **'KL0497'** | **'KL0853'** | 96.0 | [94.0 - 97.4] | 88.7 | [86.2 - 90.7] | 96.8 | [95.3 - 97.8] | 0.1 |
| **'KL0867'** | **'KL0876'** | 96.2 | [94.3 - 97.5] | 88.7 | [86.3 - 90.8] | 96.9 | [95.5 - 97.9] | 0.06 |
| **'KL0540'** | *Corynebacterium ulcerans* NCTC 7910 | 97.9 | [96.6 - 98.8] | 88.7 | [86.3 - 90.8] | 98.1 | [97.1 - 98.8] | 0.05 |
| **'KL0540'** | **'KL0876'** | 97.4 | [95.9 - 98.4] | 88.6 | [86.2 - 90.7] | 97.7 | [96.6 - 98.5] | 0.08 |
| **'KL0497'** | *Corynebacterium ulcerans* NCTC 7910 | 97.8 | [96.4 - 98.6] | 88.6 | [86.1 - 90.6] | 98.0 | [96.9 - 98.7] | 0.01 |
| **'KL0876'** | *Corynebacterium ulcerans* NCTC 7910 | 97.7 | [96.2 - 98.6] | 88.5 | [86.0 - 90.6] | 97.9 | [96.8 - 98.6] | 0.04 |
| **'KL0853'** | *Corynebacterium ulcerans* NCTC 7910 | 95.3 | [93.1 - 96.8] | 88.5 | [86.1 - 90.6] | 96.2 | [94.6 - 97.4] | 0.1 |
| **'KL0483'** | **'KL0870'** | 97.4 | [95.9 - 98.4] | 88.4 | [86.0 - 90.5] | 97.7 | [96.5 - 98.5] | 0.03 |
| **'KL0475'** | *Corynebacterium ulcerans* NCTC 7910 | 91.2 | [88.1 - 93.5] | 88.4 | [85.9 - 90.5] | 93.2 | [90.9 - 95.0] | 0.01 |
| **'KL0515'** | **'KL0870'** | 94.8 | [92.5 - 96.5] | 88.4 | [86.0 - 90.5] | 95.9 | [94.2 - 97.1] | 0.04 |
| **'KL0840'** | *Corynebacterium ulcerans* NCTC 7910 | 98.1 | [96.9 - 98.9] | 88.3 | [85.8 - 90.4] | 98.2 | [97.2 - 98.8] | 0.0 |
| **'KL0846'** | *Corynebacterium ulcerans* NCTC 7910 | 98.0 | [96.7 - 98.8] | 88.3 | [85.8 - 90.4] | 98.1 | [97.1 - 98.8] | 0.0 |
| **'KL0796'** | *Corynebacterium ulcerans* NCTC 7910 | 94.3 | [91.8 - 96.0] | 88.2 | [85.7 - 90.3] | 95.4 | [93.6 - 96.8] | 0.04 |
| **'KL0825'** | *Corynebacterium ulcerans* NCTC 7910 | 97.2 | [95.5 - 98.2] | 88.2 | [85.7 - 90.3] | 97.5 | [96.3 - 98.3] | 0.0 |
| **'KL0832'** | *Corynebacterium ulcerans* NCTC 7910 | 98.2 | [97.0 - 98.9] | 88.2 | [85.7 - 90.3] | 98.3 | [97.3 - 98.9] | 0.0 |
| **'KL0541'** | **'KL0870'** | 95.3 | [93.1 - 96.8] | 88.2 | [85.7 - 90.3] | 96.2 | [94.5 - 97.3] | 0.0 |

| Query | Subject | $d_0$ | C.I. $d_0$ | $d_4$ | C.I. $d_4$ | $d_6$ | C.I. $d_6$ | Diff. G+C Percent |
|---|---|---|---|---|---|---|---|---|
| **'KL0870'** | *Corynebacterium ulcerans* NCTC 7910 | 94.7 | [92.3 - 96.3] | 88.1 | [85.6 - 90.2] | 95.7 | [94.0 - 97.0] | 0.06 |
| **'KL0483'** | *Corynebacterium ulcerans* NCTC 7910 | 95.2 | [92.9 - 96.7] | 88.0 | [85.5 - 90.1] | 96.1 | [94.4 - 97.3] | 0.08 |
| **'KL0501'** | **'KL0876'** | 97.8 | [96.3 - 98.6] | 88.0 | [85.5 - 90.1] | 97.9 | [96.8 - 98.6] | 0.03 |
| **'KL0785'** | **'KL0870'** | 96.2 | [94.2 - 97.5] | 87.9 | [85.4 - 90.1] | 96.8 | [95.3 - 97.8] | 0.0 |
| **'KL0818'** | **'KL0870'** | 96.3 | [94.3 - 97.6] | 87.9 | [85.4 - 90.0] | 96.8 | [95.4 - 97.9] | 0.01 |
| **'KL0846'** | **'KL0853'** | 96.2 | [94.3 - 97.5] | 87.9 | [85.3 - 90.0] | 96.8 | [95.3 - 97.8] | 0.09 |
| **'KL0840'** | **'KL0853'** | 96.4 | [94.5 - 97.6] | 87.9 | [85.4 - 90.0] | 96.9 | [95.5 - 97.9] | 0.1 |
| **'KL0556'** | *Corynebacterium ulcerans* NCTC 7910 | 98.1 | [96.9 - 98.9] | 87.9 | [85.3 - 90.0] | 98.2 | [97.2 - 98.8] | 0.01 |
| **'KL0832'** | **'KL0853'** | 96.5 | [94.6 - 97.7] | 87.8 | [85.3 - 89.9] | 97.0 | [95.6 - 98.0] | 0.1 |
| **'KL0501'** | **'KL0853'** | 96.4 | [94.4 - 97.6] | 87.8 | [85.2 - 89.9] | 96.9 | [95.5 - 97.9] | 0.1 |
| **'KL0501'** | *Corynebacterium ulcerans* NCTC 7910 | 97.9 | [96.6 - 98.7] | 87.7 | [85.2 - 89.9] | 98.0 | [96.9 - 98.7] | 0.0 |
| **'KL0825'** | **'KL0853'** | 95.3 | [93.1 - 96.9] | 87.7 | [85.2 - 89.9] | 96.2 | [94.5 - 97.3] | 0.1 |
| **'KL0497'** | **'KL0818'** | 96.8 | [95.0 - 98.0] | 87.6 | [85.0 - 89.7] | 97.2 | [95.9 - 98.1] | 0.05 |
| **'KL0497'** | **'KL0785'** | 96.7 | [94.9 - 97.9] | 87.6 | [85.0 - 89.7] | 97.1 | [95.8 - 98.1] | 0.06 |
| **'KL0556'** | **'KL0870'** | 95.4 | [93.2 - 96.9] | 87.6 | [85.0 - 89.7] | 96.2 | [94.5 - 97.3] | 0.05 |
| **'KL0497'** | **'KL0515'** | 97.8 | [96.4 - 98.7] | 87.4 | [84.8 - 89.6] | 97.9 | [96.8 - 98.6] | 0.02 |
| **'KL0475'** | **'KL0853'** | 92.6 | [89.8 - 94.7] | 87.3 | [84.8 - 89.5] | 94.1 | [92.0 - 95.7] | 0.09 |
| **'KL0540'** | **'KL0853'** | 95.7 | [93.6 - 97.2] | 87.3 | [84.7 - 89.5] | 96.4 | [94.8 - 97.5] | 0.05 |
| **'KL0556'** | **'KL0876'** | 97.6 | [96.1 - 98.5] | 87.2 | [84.6 - 89.4] | 97.7 | [96.5 - 98.5] | 0.05 |
| **'KL0867'** | **'KL0870'** | 94.8 | [92.5 - 96.5] | 87.2 | [84.6 - 89.4] | 95.7 | [94.0 - 97.0] | 0.03 |
| **'KL0547'** | **'KL0870'** | 94.0 | [91.4 - 95.8] | 87.2 | [84.6 - 89.4] | 95.1 | [93.2 - 96.5] | 0.03 |
| **'KL0853'** | **'KL0867'** | 95.5 | [93.3 - 97.0] | 87.2 | [84.6 - 89.4] | 96.2 | [94.6 - 97.4] | 0.07 |
| **'KL0541'** | **'KL0853'** | 94.0 | [91.5 - 95.8] | 87.2 | [84.6 - 89.4] | 95.2 | [93.3 - 96.5] | 0.04 |
| **'KL0818'** | **'KL0832'** | 97.2 | [95.6 - 98.3] | 87.1 | [84.5 - 89.3] | 97.4 | [96.2 - 98.3] | 0.05 |
| **'KL0785'** | **'KL0840'** | 97.0 | [95.3 - 98.1] | 87.1 | [84.5 - 89.3] | 97.3 | [96.0 - 98.2] | 0.05 |
| **'KL0515'** | **'KL0846'** | 97.9 | [96.5 - 98.7] | 87.1 | [84.5 - 89.3] | 97.9 | [96.8 - 98.6] | 0.01 |
| **'KL0796'** | **'KL0853'** | 95.5 | [93.4 - 97.0] | 87.1 | [84.5 - 89.3] | 96.2 | [94.6 - 97.4] | 0.06 |

| Query | Subject | $d_0$ | C.I. $d_0$ | $d_4$ | C.I. $d_4$ | $d_6$ | C.I. $d_6$ | Diff. G+C Percent |
|---|---|---|---|---|---|---|---|---|
| **'KL0785'** | **'KL0832'** | 97.1 | [95.5 - 98.2] | 87.1 | [84.5 - 89.3] | 97.4 | [96.1 - 98.2] | 0.05 |
| **'KL0515'** | **'KL0840'** | 98.0 | [96.7 - 98.8] | 87.1 | [84.5 - 89.3] | 98.0 | [97.0 - 98.7] | 0.01 |
| **'KL0818'** | **'KL0840'** | 97.1 | [95.4 - 98.2] | 87.1 | [84.5 - 89.3] | 97.4 | [96.1 - 98.2] | 0.05 |
| **'KL0785'** | **'KL0846'** | 96.8 | [95.1 - 98.0] | 87.1 | [84.6 - 89.3] | 97.2 | [95.8 - 98.1] | 0.05 |
| **'KL0818'** | **'KL0846'** | 96.9 | [95.2 - 98.1] | 87.1 | [84.6 - 89.3] | 97.3 | [95.9 - 98.2] | 0.04 |
| **'KL0515'** | **'KL0541'** | 92.9 | [90.1 - 94.9] | 87.0 | [84.4 - 89.2] | 94.3 | [92.2 - 95.8] | 0.04 |
| **'KL0483'** | **'KL0515'** | 94.9 | [92.6 - 96.6] | 87.0 | [84.4 - 89.2] | 95.8 | [94.1 - 97.0] | 0.07 |
| **'KL0541'** | **'KL0556'** | 94.1 | [91.6 - 95.9] | 87.0 | [84.4 - 89.2] | 95.2 | [93.3 - 96.6] | 0.04 |
| **'KL0515'** | **'KL0832'** | 98.1 | [96.9 - 98.9] | 87.0 | [84.4 - 89.2] | 98.1 | [97.1 - 98.8] | 0.02 |
| **'KL0475'** | **'KL0556'** | 92.3 | [89.4 - 94.5] | 87.0 | [84.4 - 89.2] | 93.9 | [91.7 - 95.5] | 0.0 |
| **'KL0515'** | **'KL0825'** | 97.0 | [95.3 - 98.1] | 87.0 | [84.4 - 89.2] | 97.3 | [96.0 - 98.2] | 0.01 |
| **'KL0540'** | **'KL0818'** | 96.7 | [94.8 - 97.9] | 86.9 | [84.3 - 89.1] | 97.0 | [95.6 - 98.0] | 0.0 |
| **'KL0540'** | **'KL0785'** | 96.6 | [94.7 - 97.8] | 86.9 | [84.3 - 89.1] | 96.9 | [95.5 - 97.9] | 0.0 |
| **'KL0515'** | **'KL0867'** | 96.4 | [94.5 - 97.7] | 86.9 | [84.3 - 89.1] | 96.8 | [95.4 - 97.8] | 0.01 |
| **'KL0475'** | **'KL0515'** | 91.1 | [88.0 - 93.5] | 86.9 | [84.4 - 89.2] | 93.0 | [90.6 - 94.7] | 0.0 |
| **'KL0515'** | **'KL0796'** | 94.2 | [91.7 - 96.0] | 86.9 | [84.3 - 89.1] | 95.2 | [93.4 - 96.6] | 0.03 |
| **'KL0556'** | **'KL0785'** | 96.8 | [95.1 - 98.0] | 86.9 | [84.3 - 89.1] | 97.1 | [95.8 - 98.1] | 0.04 |
| **'KL0818'** | **'KL0825'** | 96.1 | [94.1 - 97.5] | 86.9 | [84.3 - 89.1] | 96.6 | [95.1 - 97.7] | 0.05 |
| **'KL0785'** | **'KL0825'** | 96.0 | [94.0 - 97.4] | 86.9 | [84.3 - 89.1] | 96.6 | [95.0 - 97.6] | 0.05 |
| **'KL0556'** | **'KL0867'** | 97.5 | [95.9 - 98.4] | 86.9 | [84.3 - 89.1] | 97.6 | [96.4 - 98.4] | 0.02 |
| **'KL0515'** | **'KL0547'** | 95.5 | [93.3 - 97.0] | 86.9 | [84.3 - 89.1] | 96.2 | [94.5 - 97.3] | 0.02 |
| **'KL0515'** | **'KL0540'** | 97.5 | [96.0 - 98.5] | 86.9 | [84.3 - 89.1] | 97.6 | [96.4 - 98.4] | 0.03 |
| **'KL0497'** | **'KL0556'** | 98.3 | [97.1 - 99.0] | 86.9 | [84.3 - 89.1] | 98.2 | [97.2 - 98.8] | 0.02 |
| **'KL0515'** | **'KL0556'** | 97.9 | [96.5 - 98.7] | 86.8 | [84.2 - 89.0] | 97.9 | [96.8 - 98.6] | 0.0 |
| **'KL0483'** | **'KL0853'** | 97.1 | [95.5 - 98.2] | 86.8 | [84.2 - 89.1] | 97.4 | [96.1 - 98.2] | 0.01 |
| **'KL0556'** | **'KL0846'** | 98.7 | [97.7 - 99.3] | 86.8 | [84.2 - 89.0] | 98.5 | [97.6 - 99.1] | 0.01 |
| **'KL0540'** | **'KL0870'** | 95.9 | [93.9 - 97.3] | 86.8 | [84.2 - 89.0] | 96.5 | [94.9 - 97.6] | 0.01 |

| Query | Subject | $d_0$ | C.I. $d_0$ | $d_4$ | C.I. $d_4$ | $d_6$ | C.I. $d_6$ | Diff. G+C Percent |
|---|---|---|---|---|---|---|---|---|
| **'KL0547'** | **'KL0818'** | 94.5 | [92.1 - 96.2] | 86.8 | [84.2 - 89.1] | 95.5 | [93.6 - 96.8] | 0.01 |
| **'KL0547'** | **'KL0556'** | 96.6 | [94.8 - 97.8] | 86.8 | [84.2 - 89.0] | 97.0 | [95.6 - 98.0] | 0.02 |
| **'KL0547'** | **'KL0785'** | 94.4 | [91.9 - 96.1] | 86.8 | [84.2 - 89.0] | 95.4 | [93.5 - 96.7] | 0.02 |
| **'KL0540'** | **'KL0556'** | 98.5 | [97.4 - 99.1] | 86.8 | [84.2 - 89.0] | 98.4 | [97.4 - 99.0] | 0.04 |
| **'KL0547'** | **'KL0853'** | 94.7 | [92.4 - 96.4] | 86.8 | [84.2 - 89.1] | 95.6 | [93.8 - 96.9] | 0.07 |
| **'KL0556'** | **'KL0818'** | 96.9 | [95.2 - 98.0] | 86.8 | [84.2 - 89.1] | 97.2 | [95.9 - 98.1] | 0.04 |
| **'KL0556'** | **'KL0853'** | 95.8 | [93.8 - 97.2] | 86.8 | [84.2 - 89.0] | 96.4 | [94.8 - 97.5] | 0.09 |
| **'KL0556'** | **'KL0796'** | 95.3 | [93.0 - 96.8] | 86.7 | [84.1 - 89.0] | 96.0 | [94.3 - 97.2] | 0.03 |
| **'KL0483'** | **'KL0556'** | 96.0 | [94.0 - 97.4] | 86.7 | [84.1 - 88.9] | 96.5 | [95.0 - 97.6] | 0.08 |
| **'KL0556'** | **'KL0840'** | 98.8 | [97.9 - 99.3] | 86.7 | [84.1 - 89.0] | 98.6 | [97.7 - 99.1] | 0.01 |
| **'KL0785'** | **'KL0867'** | 95.5 | [93.3 - 97.0] | 86.7 | [84.1 - 88.9] | 96.2 | [94.5 - 97.3] | 0.03 |
| **'KL0818'** | **'KL0867'** | 95.6 | [93.5 - 97.1] | 86.7 | [84.1 - 88.9] | 96.2 | [94.6 - 97.4] | 0.02 |
| **'KL0556'** | **'KL0825'** | 98.0 | [96.6 - 98.8] | 86.6 | [84.0 - 88.8] | 97.9 | [96.8 - 98.7] | 0.01 |
| **'KL0556'** | **'KL0832'** | 98.9 | [98.0 - 99.4] | 86.6 | [84.0 - 88.9] | 98.6 | [97.8 - 99.2] | 0.01 |
| **'KL0501'** | **'KL0515'** | 98.2 | [97.0 - 98.9] | 86.5 | [83.9 - 88.8] | 98.1 | [97.1 - 98.8] | 0.02 |
| **'KL0501'** | **'KL0818'** | 97.3 | [95.7 - 98.3] | 86.5 | [83.9 - 88.8] | 97.5 | [96.2 - 98.3] | 0.05 |
| **'KL0501'** | **'KL0785'** | 97.2 | [95.6 - 98.2] | 86.5 | [83.9 - 88.7] | 97.4 | [96.1 - 98.3] | 0.06 |
| **'KL0501'** | **'KL0556'** | 98.9 | [98.0 - 99.4] | 86.5 | [83.9 - 88.8] | 98.6 | [97.8 - 99.2] | 0.01 |
| **'KL0785'** | **'KL0796'** | 95.6 | [93.5 - 97.1] | 86.4 | [83.7 - 88.6] | 96.2 | [94.6 - 97.4] | 0.01 |
| **'KL0796'** | **'KL0818'** | 95.7 | [93.6 - 97.2] | 86.4 | [83.7 - 88.6] | 96.3 | [94.7 - 97.4] | 0.01 |
| **'KL0541'** | **'KL0818'** | 94.6 | [92.2 - 96.3] | 86.3 | [83.7 - 88.6] | 95.4 | [93.6 - 96.8] | 0.01 |
| **'KL0475'** | **'KL0785'** | 92.8 | [90.0 - 94.9] | 86.3 | [83.7 - 88.6] | 94.1 | [92.0 - 95.7] | 0.04 |
| **'KL0475'** | **'KL0818'** | 93.0 | [90.2 - 95.0] | 86.3 | [83.7 - 88.6] | 94.2 | [92.2 - 95.8] | 0.04 |
| **'KL0541'** | **'KL0785'** | 94.4 | [92.0 - 96.2] | 86.3 | [83.7 - 88.6] | 95.4 | [93.5 - 96.7] | 0.0 |
| **'KL0483'** | **'KL0818'** | 96.6 | [94.7 - 97.8] | 86.3 | [83.6 - 88.5] | 96.9 | [95.4 - 97.9] | 0.04 |
| **'KL0483'** | **'KL0785'** | 96.4 | [94.6 - 97.7] | 86.2 | [83.6 - 88.5] | 96.8 | [95.3 - 97.8] | 0.03 |
| **'KL0515'** | *Corynebacterium silvaticum* KL0182 | 91.6 | [88.6 - 93.9] | 41.1 | [38.6 - 43.6] | 81.5 | [78.1 - 84.4] | 1.12 |

| Query | Subject | $d_0$ | C.I. $d_0$ | $d_4$ | C.I. $d_4$ | $d_6$ | C.I. $d_6$ | Diff. G+C Percent |
|---|---|---|---|---|---|---|---|---|
| **'KL0785'** | *Corynebacterium silvaticum* KL0182 | 90.3 | [87.1 - 92.8] | 41.1 | [38.6 - 43.6] | 80.3 | [76.9 - 83.4] | 1.08 |
| **'KL0876'** | *Corynebacterium silvaticum* KL0182 | 90.9 | [87.8 - 93.3] | 41.1 | [38.6 - 43.6] | 80.9 | [77.5 - 83.8] | 1.17 |
| **'KL0818'** | *Corynebacterium silvaticum* KL0182 | 90.3 | [87.1 - 92.8] | 41.1 | [38.6 - 43.7] | 80.4 | [77.0 - 83.4] | 1.09 |
| **'KL0853'** | *Corynebacterium silvaticum* KL0182 | 87.8 | [84.3 - 90.6] | 41.1 | [38.6 - 43.7] | 78.2 | [74.8 - 81.3] | 1.04 |
| **'KL0832'** | *Corynebacterium silvaticum* KL0182 | 91.6 | [88.6 - 93.9] | 40.9 | [38.4 - 43.5] | 81.4 | [78.0 - 84.3] | 1.14 |
| **'KL0547'** | *Corynebacterium silvaticum* KL0182 | 88.9 | [85.5 - 91.6] | 40.9 | [38.4 - 43.4] | 79.0 | [75.6 - 82.1] | 1.11 |
| **'KL0825'** | *Corynebacterium silvaticum* KL0182 | 90.1 | [86.8 - 92.6] | 40.9 | [38.4 - 43.5] | 80.1 | [76.6 - 83.1] | 1.14 |
| **'KL0501'** | *Corynebacterium silvaticum* KL0182 | 91.5 | [88.5 - 93.8] | 40.9 | [38.4 - 43.4] | 81.3 | [78.0 - 84.3] | 1.14 |
| **'KL0556'** | *Corynebacterium silvaticum* KL0182 | 92.7 | [89.9 - 94.8] | 40.9 | [38.5 - 43.5] | 82.4 | [79.1 - 85.3] | 1.13 |
| **'KL0867'** | *Corynebacterium silvaticum* KL0182 | 90.0 | [86.8 - 92.5] | 40.9 | [38.4 - 43.4] | 80.0 | [76.6 - 83.0] | 1.11 |
| **'KL0497'** | *Corynebacterium silvaticum* KL0182 | 91.4 | [88.3 - 93.7] | 40.9 | [38.4 - 43.4] | 81.2 | [77.8 - 84.1] | 1.14 |
| **'KL0846'** | *Corynebacterium silvaticum* KL0182 | 91.5 | [88.4 - 93.8] | 40.9 | [38.4 - 43.4] | 81.2 | [77.9 - 84.2] | 1.13 |
| **'KL0840'** | *Corynebacterium silvaticum* KL0182 | 91.6 | [88.6 - 93.9] | 40.9 | [38.4 - 43.4] | 81.4 | [78.0 - 84.3] | 1.14 |
| **'KL0870'** | *Corynebacterium silvaticum* KL0182 | 88.5 | [85.1 - 91.3] | 40.9 | [38.4 - 43.4] | 78.7 | [75.3 - 81.8] | 1.08 |
| **'KL0475'** | *Corynebacterium silvaticum* KL0182 | 83.5 | [79.7 - 86.7] | 40.8 | [38.3 - 43.4] | 74.6 | [71.1 - 77.7] | 1.13 |
| **'KL0540'** | *Corynebacterium silvaticum* KL0182 | 91.6 | [88.6 - 93.9] | 40.7 | [38.2 - 43.2] | 81.3 | [77.9 - 84.2] | 1.09 |
| **'KL0483'** | *Corynebacterium silvaticum* KL0182 | 88.2 | [84.7 - 91.0] | 40.7 | [38.2 - 43.3] | 78.4 | [74.9 - 81.4] | 1.05 |
| **'KL0541'** | *Corynebacterium silvaticum* KL0182 | 86.4 | [82.8 - 89.4] | 40.7 | [38.3 - 43.3] | 76.9 | [73.4 - 80.0] | 1.08 |
| **'KL0796'** | *Corynebacterium silvaticum* KL0182 | 87.8 | [84.3 - 90.6] | 40.7 | [38.3 - 43.3] | 78.1 | [74.6 - 81.1] | 1.1 |
| **'KL0832'** | *Corynebacterium pseudotuberculosis* ATCC 19410 | 86.8 | [83.2 - 89.7] | 27.6 | [25.2 - 30.1] | 67.9 | [64.5 - 71.1] | 1.13 |
| **'KL0515'** | *Corynebacterium pseudotuberculosis* ATCC 19410 | 86.7 | [83.1 - 89.6] | 27.6 | [25.2 - 30.0] | 67.8 | [64.3 - 71.0] | 1.14 |
| **'KL0497'** | *Corynebacterium pseudotuberculosis* ATCC 19410 | 86.5 | [82.8 - 89.4] | 27.6 | [25.2 - 30.0] | 67.6 | [64.2 - 70.8] | 1.12 |
| **'KL0840'** | *Corynebacterium pseudotuberculosis* DSM 20689 | 86.8 | [83.2 - 89.7] | 27.6 | [25.2 - 30.1] | 67.9 | [64.4 - 71.1] | 1.13 |
| **'KL0483'** | *Corynebacterium pseudotuberculosis* ATCC 19410 | 82.2 | [78.4 - 85.5] | 27.6 | [25.2 - 30.1] | 64.6 | [61.2 - 67.8] | 1.21 |
| **'KL0818'** | *Corynebacterium pseudotuberculosis* ATCC 19410 | 84.8 | [81.1 - 87.9] | 27.6 | [25.2 - 30.1] | 66.4 | [63.0 - 69.6] | 1.18 |

| Query | Subject | $d_0$ | C.I. $d_0$ | $d_4$ | C.I. $d_4$ | $d_6$ | C.I. $d_6$ | Diff. G+C Percent |
|---|---|---|---|---|---|---|---|---|
| **'KL0541'** | *Corynebacterium pseudotuberculosis* ATCC 19410 | 80.5 | [76.6 - 84.0] | 27.6 | [25.2 - 30.0] | 63.4 | [60.1 - 66.6] | 1.18 |
| **'KL0832'** | *Corynebacterium pseudotuberculosis* DSM 20689 | 86.8 | [83.2 - 89.7] | 27.6 | [25.2 - 30.1] | 67.9 | [64.5 - 71.1] | 1.12 |
| **'KL0483'** | *Corynebacterium pseudotuberculosis* DSM 20689 | 82.2 | [78.4 - 85.6] | 27.6 | [25.2 - 30.1] | 64.6 | [61.3 - 67.8] | 1.21 |
| **'KL0876'** | *Corynebacterium pseudotuberculosis* ATCC 19410 | 86.5 | [82.8 - 89.4] | 27.6 | [25.3 - 30.1] | 67.7 | [64.2 - 70.9] | 1.09 |
| **'KL0853'** | *Corynebacterium pseudotuberculosis* ATCC 19410 | 83.5 | [79.6 - 86.7] | 27.6 | [25.2 - 30.1] | 65.5 | [62.1 - 68.7] | 1.23 |
| **'KL0556'** | *Corynebacterium pseudotuberculosis* ATCC 19410 | 88.0 | [84.6 - 90.8] | 27.6 | [25.2 - 30.0] | 68.8 | [65.3 - 72.0] | 1.14 |
| **'KL0497'** | *Corynebacterium pseudotuberculosis* DSM 20689 | 86.5 | [82.9 - 89.4] | 27.6 | [25.2 - 30.1] | 67.6 | [64.2 - 70.9] | 1.12 |
| **'KL0853'** | *Corynebacterium pseudotuberculosis* DSM 20689 | 83.6 | [79.8 - 86.8] | 27.6 | [25.2 - 30.1] | 65.6 | [62.2 - 68.8] | 1.22 |
| **'KL0846'** | *Corynebacterium pseudotuberculosis* DSM 20689 | 86.6 | [83.0 - 89.6] | 27.6 | [25.2 - 30.1] | 67.7 | [64.3 - 71.0] | 1.13 |
| **'KL0515'** | *Corynebacterium pseudotuberculosis* DSM 20689 | 86.7 | [83.1 - 89.6] | 27.6 | [25.2 - 30.0] | 67.8 | [64.4 - 71.0] | 1.14 |
| **'KL0796'** | *Corynebacterium pseudotuberculosis* ATCC 19410 | 81.4 | [77.5 - 84.7] | 27.6 | [25.2 - 30.1] | 64.0 | [60.7 - 67.2] | 1.17 |
| **'KL0796'** | *Corynebacterium pseudotuberculosis* DSM 20689 | 81.4 | [77.5 - 84.7] | 27.6 | [25.2 - 30.1] | 64.0 | [60.7 - 67.2] | 1.17 |
| **'KL0870'** | *Corynebacterium pseudotuberculosis* DSM 20689 | 82.6 | [78.8 - 85.9] | 27.6 | [25.3 - 30.1] | 64.9 | [61.5 - 68.1] | 1.18 |
| **'KL0556'** | *Corynebacterium pseudotuberculosis* DSM 20689 | 88.0 | [84.6 - 90.8] | 27.6 | [25.2 - 30.1] | 68.8 | [65.4 - 72.0] | 1.14 |
| **'KL0846'** | *Corynebacterium pseudotuberculosis* ATCC 19410 | 86.6 | [83.0 - 89.6] | 27.6 | [25.2 - 30.1] | 67.7 | [64.3 - 71.0] | 1.13 |
| **'KL0840'** | *Corynebacterium pseudotuberculosis* ATCC 19410 | 86.8 | [83.2 - 89.7] | 27.6 | [25.2 - 30.1] | 67.8 | [64.4 - 71.1] | 1.13 |
| **'KL0825'** | *Corynebacterium pseudotuberculosis* DSM 20689 | 85.1 | [81.4 - 88.2] | 27.6 | [25.2 - 30.1] | 66.7 | [63.3 - 69.9] | 1.13 |
| **'KL0876'** | *Corynebacterium pseudotuberculosis* DSM 20689 | 86.5 | [82.9 - 89.4] | 27.6 | [25.3 - 30.1] | 67.7 | [64.3 - 70.9] | 1.09 |
| **'KL0825'** | *Corynebacterium pseudotuberculosis* ATCC 19410 | 85.1 | [81.4 - 88.2] | 27.6 | [25.2 - 30.1] | 66.7 | [63.3 - 69.9] | 1.13 |
| **'KL0541'** | *Corynebacterium pseudotuberculosis* DSM 20689 | 80.6 | [76.6 - 84.0] | 27.6 | [25.2 - 30.0] | 63.4 | [60.1 - 66.6] | 1.18 |
| **'KL0870'** | *Corynebacterium pseudotuberculosis* ATCC 19410 | 82.6 | [78.8 - 85.9] | 27.6 | [25.2 - 30.1] | 64.9 | [61.5 - 68.1] | 1.19 |

| Query | Subject | $d_0$ | C.I. $d_0$ | $d_4$ | C.I. $d_4$ | $d_6$ | C.I. $d_6$ | Diff. G+C Percent |
|---|---|---|---|---|---|---|---|---|
| **'KL0818'** | *Corynebacterium pseudotuberculosis* DSM 20689 | 84.9 | [81.2 - 88.0] | 27.6 | [25.2 - 30.1] | 66.5 | [63.1 - 69.7] | 1.17 |
| **'KL0867'** | *Corynebacterium pseudotuberculosis* DSM 20689 | 85.1 | [81.3 - 88.2] | 27.5 | [25.2 - 30.0] | 66.6 | [63.2 - 69.8] | 1.15 |
| **'KL0547'** | *Corynebacterium pseudotuberculosis* DSM 20689 | 83.7 | [79.9 - 86.9] | 27.5 | [25.2 - 30.0] | 65.6 | [62.2 - 68.8] | 1.16 |
| **'KL0867'** | *Corynebacterium pseudotuberculosis* ATCC 19410 | 85.0 | [81.3 - 88.1] | 27.5 | [25.2 - 30.0] | 66.5 | [63.2 - 69.8] | 1.16 |
| **'KL0785'** | *Corynebacterium pseudotuberculosis* ATCC 19410 | 85.3 | [81.6 - 88.4] | 27.5 | [25.2 - 30.0] | 66.7 | [63.3 - 70.0] | 1.18 |
| **'KL0540'** | *Corynebacterium pseudotuberculosis* ATCC 19410 | 86.6 | [83.0 - 89.6] | 27.5 | [25.2 - 30.0] | 67.7 | [64.3 - 70.9] | 1.18 |
| **'KL0785'** | *Corynebacterium pseudotuberculosis* DSM 20689 | 85.4 | [81.7 - 88.5] | 27.5 | [25.2 - 30.0] | 66.8 | [63.4 - 70.1] | 1.18 |
| **'KL0501'** | *Corynebacterium pseudotuberculosis* ATCC 19410 | 86.9 | [83.3 - 89.8] | 27.5 | [25.2 - 30.0] | 67.9 | [64.5 - 71.1] | 1.13 |
| **'KL0547'** | *Corynebacterium pseudotuberculosis* ATCC 19410 | 83.7 | [79.9 - 86.9] | 27.5 | [25.2 - 30.0] | 65.6 | [62.2 - 68.8] | 1.16 |
| **'KL0501'** | *Corynebacterium pseudotuberculosis* DSM 20689 | 86.9 | [83.3 - 89.8] | 27.5 | [25.2 - 30.0] | 67.9 | [64.5 - 71.2] | 1.12 |
| **'KL0540'** | *Corynebacterium pseudotuberculosis* DSM 20689 | 86.6 | [83.0 - 89.6] | 27.5 | [25.2 - 30.0] | 67.7 | [64.3 - 71.0] | 1.17 |
| **'KL0475'** | *Corynebacterium pseudotuberculosis* DSM 20689 | 77.8 | [73.8 - 81.3] | 27.4 | [25.1 - 29.9] | 61.4 | [58.2 - 64.6] | 1.14 |
| **'KL0475'** | *Corynebacterium pseudotuberculosis* ATCC 19410 | 77.8 | [73.8 - 81.3] | 27.4 | [25.1 - 29.9] | 61.4 | [58.1 - 64.6] | 1.14 |
| **'KL0870'** | *Corynebacterium mustelae* DSM 45274 | 12.9 | [10.2 - 16.2] | 25.2 | [22.9 - 27.7] | 13.3 | [11.0 - 16.1] | 0.8 |
| **'KL0876'** | *Corynebacterium mustelae* DSM 45274 | 13.0 | [10.2 - 16.2] | 25.2 | [22.8 - 27.7] | 13.3 | [11.0 - 16.1] | 0.71 |
| **'KL0846'** | *Corynebacterium mustelae* DSM 45274 | 13.0 | [10.2 - 16.2] | 25.2 | [22.8 - 27.6] | 13.3 | [11.0 - 16.1] | 0.74 |
| **'KL0867'** | *Corynebacterium mustelae* DSM 45274 | 13.0 | [10.2 - 16.2] | 25.2 | [22.8 - 27.6] | 13.3 | [11.0 - 16.1] | 0.77 |
| **'KL0540'** | *Corynebacterium mustelae* DSM 45274 | 13.0 | [10.2 - 16.2] | 25.2 | [22.9 - 27.7] | 13.3 | [11.0 - 16.1] | 0.79 |
| **'KL0541'** | *Corynebacterium mustelae* DSM 45274 | 12.9 | [10.2 - 16.2] | 25.2 | [22.8 - 27.6] | 13.3 | [11.0 - 16.1] | 0.79 |
| **'KL0547'** | *Corynebacterium mustelae* DSM 45274 | 12.9 | [10.2 - 16.2] | 25.2 | [22.8 - 27.6] | 13.3 | [11.0 - 16.1] | 0.77 |
| **'KL0853'** | *Corynebacterium mustelae* DSM 45274 | 12.9 | [10.2 - 16.2] | 25.2 | [22.9 - 27.7] | 13.3 | [11.0 - 16.1] | 0.84 |
| **'KL0832'** | *Corynebacterium mustelae* DSM 45274 | 13.0 | [10.2 - 16.2] | 25.2 | [22.9 - 27.7] | 13.3 | [11.0 - 16.1] | 0.74 |
| **'KL0556'** | *Corynebacterium mustelae* DSM 45274 | 13.0 | [10.3 - 16.2] | 25.2 | [22.9 - 27.7] | 13.3 | [11.0 - 16.1] | 0.75 |

| Query | Subject | $d_0$ | C.I. $d_0$ | $d_4$ | C.I. $d_4$ | $d_6$ | C.I. $d_6$ | Diff. G+C Percent |
|---|---|---|---|---|---|---|---|---|
| **'KL0501'** | *Corynebacterium mustelae* DSM 45274 | 13.0 | [10.2 - 16.2] | 25.2 | [22.9 - 27.7] | 13.3 | [11.0 - 16.1] | 0.74 |
| **'KL0483'** | *Corynebacterium mustelae* DSM 45274 | 12.9 | [10.2 - 16.2] | 25.2 | [22.9 - 27.7] | 13.3 | [11.0 - 16.1] | 0.83 |
| **'KL0825'** | *Corynebacterium mustelae* DSM 45274 | 13.0 | [10.2 - 16.2] | 25.2 | [22.9 - 27.7] | 13.3 | [11.0 - 16.1] | 0.74 |
| **'KL0840'** | *Corynebacterium mustelae* DSM 45274 | 13.0 | [10.2 - 16.2] | 25.2 | [22.9 - 27.7] | 13.3 | [11.0 - 16.1] | 0.74 |
| **'KL0497'** | *Corynebacterium mustelae* DSM 45274 | 13.0 | [10.2 - 16.2] | 25.1 | [22.8 - 27.6] | 13.3 | [11.0 - 16.1] | 0.73 |
| **'KL0796'** | *Corynebacterium mustelae* DSM 45274 | 12.9 | [10.2 - 16.2] | 25.1 | [22.8 - 27.6] | 13.3 | [11.0 - 16.1] | 0.78 |
| **'KL0853'** | *Corynebacterium vitaeruminis* DSM 20294 | 13.2 | [10.5 - 16.5] | 25.0 | [22.7 - 27.5] | 13.6 | [11.2 - 16.3] | 12.12 |
| **'KL0483'** | *Corynebacterium vitaeruminis* DSM 20294 | 13.2 | [10.5 - 16.5] | 25.0 | [22.6 - 27.4] | 13.6 | [11.2 - 16.4] | 12.13 |
| **'KL0876'** | *Corynebacterium vitaeruminis* DSM 20294 | 13.2 | [10.5 - 16.5] | 25.0 | [22.7 - 27.5] | 13.6 | [11.2 - 16.3] | 12.25 |
| **'KL0825'** | *Corynebacterium vitaeruminis* DSM 20294 | 13.2 | [10.5 - 16.5] | 24.9 | [22.6 - 27.4] | 13.6 | [11.2 - 16.4] | 12.21 |
| **'KL0541'** | *Corynebacterium vitaeruminis* DSM 20294 | 13.2 | [10.5 - 16.5] | 24.9 | [22.6 - 27.4] | 13.6 | [11.2 - 16.4] | 12.16 |
| **'KL0846'** | *Corynebacterium vitaeruminis* DSM 20294 | 13.2 | [10.5 - 16.5] | 24.9 | [22.6 - 27.4] | 13.6 | [11.2 - 16.4] | 12.21 |
| **'KL0840'** | *Corynebacterium vitaeruminis* DSM 20294 | 13.2 | [10.5 - 16.5] | 24.9 | [22.6 - 27.4] | 13.6 | [11.2 - 16.4] | 12.21 |
| **'KL0832'** | *Corynebacterium vitaeruminis* DSM 20294 | 13.2 | [10.5 - 16.5] | 24.9 | [22.6 - 27.4] | 13.6 | [11.2 - 16.4] | 12.22 |
| **'KL0870'** | *Corynebacterium pseudopelargi* CCM 8832 | 13.2 | [10.5 - 16.5] | 24.9 | [22.6 - 27.4] | 13.6 | [11.2 - 16.4] | 4.54 |
| **'KL0867'** | *Corynebacterium vitaeruminis* DSM 20294 | 13.2 | [10.5 - 16.5] | 24.8 | [22.5 - 27.3] | 13.6 | [11.2 - 16.4] | 12.19 |
| **'KL0501'** | *Corynebacterium vitaeruminis* DSM 20294 | 13.2 | [10.5 - 16.5] | 24.8 | [22.5 - 27.3] | 13.6 | [11.2 - 16.4] | 12.22 |
| **'KL0547'** | *Corynebacterium vitaeruminis* DSM 20294 | 13.2 | [10.5 - 16.5] | 24.8 | [22.5 - 27.3] | 13.6 | [11.2 - 16.4] | 12.18 |
| **'KL0853'** | *Corynebacterium pelargi* DSM 46737 | 13.2 | [10.5 - 16.5] | 24.7 | [22.4 - 27.1] | 13.6 | [11.2 - 16.3] | 4.76 |
| **'KL0876'** | *Corynebacterium pseudopelargi* CCM 8832 | 13.2 | [10.5 - 16.5] | 24.7 | [22.4 - 27.2] | 13.6 | [11.2 - 16.4] | 4.63 |
| **'KL0515'** | *Corynebacterium vitaeruminis* DSM 20294 | 13.2 | [10.5 - 16.5] | 24.7 | [22.3 - 27.1] | 13.6 | [11.2 - 16.4] | 12.2 |
| **'KL0870'** | *Corynebacterium vitaeruminis* DSM 20294 | 13.2 | [10.5 - 16.5] | 24.7 | [22.4 - 27.1] | 13.6 | [11.2 - 16.4] | 12.16 |
| **'KL0853'** | *Corynebacterium pseudopelargi* CCM 8832 | 13.2 | [10.5 - 16.5] | 24.6 | [22.3 - 27.1] | 13.6 | [11.2 - 16.4] | 4.5 |
| **'KL0556'** | *Corynebacterium vitaeruminis* DSM 20294 | 13.2 | [10.5 - 16.5] | 24.6 | [22.3 - 27.1] | 13.6 | [11.2 - 16.4] | 12.2 |
| **'KL0540'** | *Corynebacterium vitaeruminis* DSM 20294 | 13.2 | [10.5 - 16.5] | 24.6 | [22.3 - 27.0] | 13.6 | [11.2 - 16.4] | 12.17 |
| **'KL0796'** | *Corynebacterium vitaeruminis* DSM 20294 | 13.2 | [10.5 - 16.5] | 24.6 | [22.3 - 27.1] | 13.6 | [11.2 - 16.4] | 12.17 |
| **'KL0840'** | *Corynebacterium pseudopelargi* CCM 8832 | 13.3 | [10.5 - 16.6] | 24.5 | [22.1 - 26.9] | 13.6 | [11.2 - 16.4] | 4.59 |

| Query | Subject | $d_0$ | C.I. $d_0$ | $d_4$ | C.I. $d_4$ | $d_6$ | C.I. $d_6$ | Diff. G+C Percent |
|---|---|---|---|---|---|---|---|---|
| **'KL0547'** | *Corynebacterium pseudopelargi* CCM 8832 | 13.2 | [10.5 - 16.6] | 24.5 | [22.2 - 26.9] | 13.6 | [11.2 - 16.4] | 4.56 |
| **'KL0825'** | *Corynebacterium pseudopelargi* CCM 8832 | 13.2 | [10.5 - 16.6] | 24.5 | [22.1 - 26.9] | 13.6 | [11.2 - 16.4] | 4.59 |
| **'KL0501'** | *Corynebacterium pseudopelargi* CCM 8832 | 13.3 | [10.5 - 16.6] | 24.5 | [22.2 - 27.0] | 13.6 | [11.3 - 16.4] | 4.6 |
| **'KL0832'** | *Corynebacterium pseudopelargi* CCM 8832 | 13.3 | [10.5 - 16.6] | 24.5 | [22.1 - 26.9] | 13.6 | [11.2 - 16.4] | 4.6 |
| **'KL0501'** | *Corynebacterium pelargi* DSM 46737 | 13.3 | [10.5 - 16.6] | 24.5 | [22.2 - 27.0] | 13.6 | [11.3 - 16.4] | 4.86 |
| **'KL0483'** | *Corynebacterium pseudopelargi* CCM 8832 | 13.2 | [10.5 - 16.5] | 24.5 | [22.2 - 27.0] | 13.6 | [11.2 - 16.4] | 4.51 |
| **'KL0870'** | *Corynebacterium pelargi* DSM 46737 | 13.2 | [10.5 - 16.5] | 24.5 | [22.2 - 26.9] | 13.6 | [11.2 - 16.4] | 4.8 |
| **'KL0497'** | *Corynebacterium pelargi* DSM 46737 | 13.3 | [10.5 - 16.6] | 24.5 | [22.2 - 27.0] | 13.6 | [11.2 - 16.4] | 4.87 |
| **'KL0876'** | *Corynebacterium pelargi* DSM 46737 | 13.2 | [10.5 - 16.5] | 24.5 | [22.1 - 26.9] | 13.6 | [11.2 - 16.4] | 4.9 |
| **'KL0497'** | *Corynebacterium vitaeruminis* DSM 20294 | 13.2 | [10.5 - 16.6] | 24.5 | [22.2 - 27.0] | 13.6 | [11.2 - 16.4] | 12.22 |
| **'KL0541'** | *Corynebacterium pseudopelargi* CCM 8832 | 13.2 | [10.5 - 16.5] | 24.5 | [22.1 - 26.9] | 13.6 | [11.2 - 16.4] | 4.54 |
| **'KL0540'** | *Corynebacterium pseudopelargi* CCM 8832 | 13.3 | [10.5 - 16.6] | 24.5 | [22.2 - 26.9] | 13.6 | [11.2 - 16.4] | 4.55 |
| **'KL0867'** | *Corynebacterium pelargi* DSM 46737 | 13.3 | [10.5 - 16.6] | 24.4 | [22.1 - 26.9] | 13.6 | [11.2 - 16.4] | 4.83 |
| **'KL0785'** | *Corynebacterium vitaeruminis* DSM 20294 | 13.2 | [10.5 - 16.5] | 24.4 | [22.1 - 26.9] | 13.6 | [11.2 - 16.4] | 12.16 |
| **'KL0818'** | *Corynebacterium vitaeruminis* DSM 20294 | 13.2 | [10.5 - 16.5] | 24.4 | [22.1 - 26.9] | 13.6 | [11.2 - 16.4] | 12.17 |
| **'KL0825'** | *Corynebacterium pelargi* DSM 46737 | 13.3 | [10.5 - 16.6] | 24.4 | [22.1 - 26.9] | 13.6 | [11.2 - 16.4] | 4.86 |
| **'KL0840'** | *Corynebacterium pelargi* DSM 46737 | 13.3 | [10.5 - 16.6] | 24.4 | [22.1 - 26.9] | 13.6 | [11.3 - 16.4] | 4.86 |
| **'KL0475'** | *Corynebacterium mustelae* DSM 45274 | 12.8 | [10.1 - 16.1] | 24.4 | [22.1 - 26.9] | 13.2 | [10.8 - 15.9] | 0.75 |
| **'KL0540'** | *Corynebacterium pelargi* DSM 46737 | 13.3 | [10.5 - 16.6] | 24.4 | [22.1 - 26.9] | 13.6 | [11.3 - 16.4] | 4.81 |
| **'KL0846'** | *Corynebacterium pelargi* DSM 46737 | 13.3 | [10.5 - 16.6] | 24.4 | [22.1 - 26.9] | 13.6 | [11.3 - 16.4] | 4.86 |
| **'KL0483'** | *Corynebacterium pelargi* DSM 46737 | 13.3 | [10.5 - 16.6] | 24.4 | [22.1 - 26.9] | 13.6 | [11.2 - 16.4] | 4.78 |
| **'KL0796'** | *Corynebacterium pelargi* DSM 46737 | 13.2 | [10.5 - 16.6] | 24.4 | [22.1 - 26.9] | 13.6 | [11.2 - 16.4] | 4.82 |
| **'KL0541'** | *Corynebacterium pelargi* DSM 46737 | 13.2 | [10.5 - 16.6] | 24.4 | [22.1 - 26.9] | 13.6 | [11.2 - 16.4] | 4.81 |
| **'KL0556'** | *Corynebacterium kutscheri* DSM 20755 | 13.1 | [10.4 - 16.4] | 24.4 | [22.1 - 26.8] | 13.5 | [11.1 - 16.3] | 6.86 |
| **'KL0832'** | *Corynebacterium pelargi* DSM 46737 | 13.3 | [10.5 - 16.6] | 24.4 | [22.1 - 26.9] | 13.6 | [11.3 - 16.4] | 4.86 |
| **'KL0846'** | *Corynebacterium pseudopelargi* CCM 8832 | 13.3 | [10.5 - 16.6] | 24.4 | [22.1 - 26.9] | 13.6 | [11.2 - 16.4] | 4.59 |
| **'KL0547'** | *Corynebacterium pelargi* DSM 46737 | 13.3 | [10.5 - 16.6] | 24.4 | [22.1 - 26.9] | 13.6 | [11.2 - 16.4] | 4.83 |

| Query | Subject | $d_0$ | C.I. $d_0$ | $d_4$ | C.I. $d_4$ | $d_6$ | C.I. $d_6$ | Diff. G+C Percent |
|---|---|---|---|---|---|---|---|---|
| **'KL0796'** | *Corynebacterium pseudopelargi* CCM 8832 | 13.2 | [10.5 - 16.6] | 24.3 | [21.9 - 26.7] | 13.6 | [11.2 - 16.4] | 4.55 |
| **'KL0497'** | *Corynebacterium pseudopelargi* CCM 8832 | 13.3 | [10.5 - 16.6] | 24.3 | [22.0 - 26.8] | 13.6 | [11.3 - 16.4] | 4.6 |
| **'KL0867'** | *Corynebacterium pseudopelargi* CCM 8832 | 13.3 | [10.5 - 16.6] | 24.3 | [22.0 - 26.7] | 13.6 | [11.2 - 16.4] | 4.57 |
| **'KL0515'** | *Corynebacterium mustelae* DSM 45274 | 13.0 | [10.3 - 16.3] | 24.1 | [21.8 - 26.6] | 13.4 | [11.0 - 16.1] | 0.75 |
| **'KL0818'** | *Corynebacterium mustelae* DSM 45274 | 13.0 | [10.3 - 16.3] | 24.1 | [21.8 - 26.5] | 13.4 | [11.0 - 16.1] | 0.79 |
| **'KL0785'** | *Corynebacterium mustelae* DSM 45274 | 13.0 | [10.3 - 16.3] | 24.1 | [21.8 - 26.5] | 13.4 | [11.0 - 16.1] | 0.79 |
| **'KL0556'** | *Corynebacterium pseudopelargi* CCM 8832 | 13.2 | [10.5 - 16.6] | 24.1 | [21.8 - 26.6] | 13.6 | [11.2 - 16.4] | 4.59 |
| **'KL0515'** | *Corynebacterium pseudopelargi* CCM 8832 | 13.3 | [10.5 - 16.6] | 24.0 | [21.7 - 26.5] | 13.6 | [11.2 - 16.4] | 4.58 |
| **'KL0556'** | *Corynebacterium pelargi* DSM 46737 | 13.3 | [10.5 - 16.6] | 24.0 | [21.7 - 26.5] | 13.6 | [11.3 - 16.4] | 4.85 |
| **'KL0818'** | *Corynebacterium pseudopelargi* CCM 8832 | 13.2 | [10.5 - 16.6] | 24.0 | [21.7 - 26.4] | 13.6 | [11.2 - 16.4] | 4.55 |
| **'KL0785'** | *Corynebacterium pseudopelargi* CCM 8832 | 13.2 | [10.5 - 16.6] | 24.0 | [21.7 - 26.5] | 13.6 | [11.2 - 16.4] | 4.54 |
| **'KL0853'** | *Corynebacterium kutscheri* DSM 20755 | 13.2 | [10.5 - 16.5] | 24.0 | [21.7 - 26.5] | 13.6 | [11.2 - 16.3] | 6.94 |
| **'KL0825'** | *Corynebacterium kutscheri* DSM 20755 | 13.2 | [10.5 - 16.5] | 23.9 | [21.6 - 26.4] | 13.5 | [11.2 - 16.3] | 6.85 |
| **'KL0483'** | *Corynebacterium kutscheri* DSM 20755 | 13.2 | [10.4 - 16.5] | 23.9 | [21.6 - 26.4] | 13.5 | [11.2 - 16.3] | 6.93 |
| **'KL0832'** | *Corynebacterium kutscheri* DSM 20755 | 13.2 | [10.5 - 16.5] | 23.9 | [21.6 - 26.4] | 13.6 | [11.2 - 16.3] | 6.85 |
| **'KL0867'** | *Corynebacterium kutscheri* DSM 20755 | 13.2 | [10.5 - 16.5] | 23.9 | [21.6 - 26.4] | 13.5 | [11.2 - 16.3] | 6.88 |
| **'KL0497'** | *Corynebacterium kutscheri* DSM 20755 | 13.2 | [10.5 - 16.5] | 23.9 | [21.6 - 26.4] | 13.6 | [11.2 - 16.3] | 6.84 |
| **'KL0501'** | *Corynebacterium kutscheri* DSM 20755 | 13.2 | [10.5 - 16.5] | 23.9 | [21.6 - 26.4] | 13.6 | [11.2 - 16.3] | 6.84 |
| **'KL0540'** | *Corynebacterium kutscheri* DSM 20755 | 13.2 | [10.5 - 16.5] | 23.9 | [21.6 - 26.4] | 13.6 | [11.2 - 16.3] | 6.9 |
| **'KL0840'** | *Corynebacterium kutscheri* DSM 20755 | 13.2 | [10.5 - 16.5] | 23.9 | [21.6 - 26.4] | 13.6 | [11.2 - 16.3] | 6.85 |
| **'KL0796'** | *Corynebacterium kutscheri* DSM 20755 | 13.2 | [10.4 - 16.5] | 23.9 | [21.6 - 26.4] | 13.5 | [11.2 - 16.3] | 6.89 |
| **'KL0846'** | *Corynebacterium kutscheri* DSM 20755 | 13.2 | [10.5 - 16.5] | 23.9 | [21.6 - 26.4] | 13.6 | [11.2 - 16.3] | 6.85 |
| **'KL0870'** | *Corynebacterium kutscheri* DSM 20755 | 13.2 | [10.5 - 16.5] | 23.8 | [21.5 - 26.2] | 13.6 | [11.2 - 16.3] | 6.9 |
| **'KL0547'** | *Corynebacterium kutscheri* DSM 20755 | 13.2 | [10.5 - 16.5] | 23.7 | [21.4 - 26.2] | 13.6 | [11.2 - 16.4] | 6.88 |
| **'KL0475'** | *Corynebacterium phocae* DSM 44612 | 12.7 | [10.1 - 16.0] | 23.7 | [21.4 - 26.2] | 13.1 | [10.8 - 15.9] | 5.49 |
| **'KL0541'** | *Corynebacterium kutscheri* DSM 20755 | 13.2 | [10.5 - 16.5] | 23.7 | [21.4 - 26.2] | 13.6 | [11.2 - 16.4] | 6.9 |
| **'KL0515'** | *Corynebacterium kutscheri* DSM 20755 | 13.2 | [10.5 - 16.5] | 23.6 | [21.3 - 26.1] | 13.6 | [11.2 - 16.4] | 6.86 |

| Query | Subject | $d_0$ | C.I. $d_0$ | $d_4$ | C.I. $d_4$ | $d_6$ | C.I. $d_6$ | Diff. G+C Percent |
|---|---|---|---|---|---|---|---|---|
| **'KL0785'** | *Corynebacterium kutscheri* DSM 20755 | 13.2 | [10.5 - 16.5] | 23.6 | [21.3 - 26.1] | 13.6 | [11.2 - 16.3] | 6.9 |
| **'KL0876'** | *Corynebacterium kutscheri* DSM 20755 | 13.2 | [10.5 - 16.5] | 23.6 | [21.3 - 26.0] | 13.6 | [11.2 - 16.4] | 6.81 |
| **'KL0818'** | *Corynebacterium kutscheri* DSM 20755 | 13.2 | [10.5 - 16.5] | 23.6 | [21.3 - 26.1] | 13.6 | [11.2 - 16.3] | 6.89 |
| **'KL0556'** | *Corynebacterium phocae* DSM 44612 | 12.9 | [10.2 - 16.2] | 23.5 | [21.2 - 25.9] | 13.3 | [10.9 - 16.0] | 5.49 |
| **'KL0475'** | *Corynebacterium vitaeruminis* DSM 20294 | 13.0 | [10.3 - 16.3] | 23.4 | [21.1 - 25.8] | 13.4 | [11.0 - 16.2] | 12.2 |
| **'KL0475'** | *Corynebacterium pelargi* DSM 46737 | 13.0 | [10.3 - 16.3] | 23.4 | [21.1 - 25.8] | 13.4 | [11.0 - 16.2] | 4.85 |
| **'KL0785'** | *Corynebacterium pelargi* DSM 46737 | 13.3 | [10.6 - 16.6] | 23.3 | [21.0 - 25.7] | 13.6 | [11.3 - 16.4] | 4.81 |
| **'KL0515'** | *Corynebacterium pelargi* DSM 46737 | 13.3 | [10.6 - 16.6] | 23.3 | [21.0 - 25.7] | 13.7 | [11.3 - 16.4] | 4.85 |
| **'KL0876'** | *Corynebacterium phocae* DSM 44612 | 12.9 | [10.2 - 16.2] | 23.3 | [21.0 - 25.8] | 13.3 | [10.9 - 16.0] | 5.53 |
| **'KL0818'** | *Corynebacterium pelargi* DSM 46737 | 13.3 | [10.6 - 16.6] | 23.3 | [21.0 - 25.7] | 13.6 | [11.3 - 16.4] | 4.82 |
| **'KL0475'** | *Corynebacterium kutscheri* DSM 20755 | 12.9 | [10.2 - 16.2] | 23.3 | [21.0 - 25.8] | 13.3 | [11.0 - 16.1] | 6.86 |
| **'KL0853'** | *Corynebacterium phocae* DSM 44612 | 12.9 | [10.2 - 16.2] | 23.2 | [20.9 - 25.6] | 13.3 | [10.9 - 16.0] | 5.4 |
| **'KL0475'** | *Corynebacterium pseudopelargi* CCM 8832 | 13.0 | [10.3 - 16.3] | 23.2 | [20.9 - 25.6] | 13.4 | [11.0 - 16.2] | 4.59 |
| **'KL0785'** | *Corynebacterium phocae* DSM 44612 | 12.9 | [10.2 - 16.2] | 23.2 | [20.9 - 25.6] | 13.3 | [10.9 - 16.0] | 5.45 |
| **'KL0818'** | *Corynebacterium phocae* DSM 44612 | 12.9 | [10.2 - 16.2] | 23.2 | [20.9 - 25.6] | 13.3 | [10.9 - 16.0] | 5.45 |
| **'KL0853'** | *Corynebacterium diphtheriae* NCTC 11397 | 14.2 | [11.3 - 17.5] | 23.2 | [20.9 - 25.6] | 14.4 | [12.0 - 17.3] | 0.12 |
| **'KL0515'** | *Corynebacterium phocae* DSM 44612 | 12.9 | [10.2 - 16.2] | 23.2 | [20.9 - 25.6] | 13.3 | [10.9 - 16.0] | 5.49 |
| **'KL0818'** | *Corynebacterium diphtheriae* NCTC 11397 | 14.2 | [11.4 - 17.6] | 23.1 | [20.8 - 25.6] | 14.5 | [12.0 - 17.3] | 0.17 |
| **'KL0515'** | *Corynebacterium diphtheriae* NCTC 11397 | 14.2 | [11.4 - 17.6] | 23.1 | [20.8 - 25.6] | 14.5 | [12.0 - 17.3] | 0.2 |
| **'KL0785'** | *Corynebacterium diphtheriae* NCTC 11397 | 14.2 | [11.4 - 17.6] | 23.1 | [20.8 - 25.6] | 14.5 | [12.0 - 17.3] | 0.16 |
| **'KL0547'** | *Corynebacterium phocae* DSM 44612 | 12.9 | [10.2 - 16.2] | 23.0 | [20.7 - 25.5] | 13.3 | [10.9 - 16.0] | 5.47 |
| **'KL0540'** | *Corynebacterium diphtheriae* NCTC 11397 | 14.2 | [11.4 - 17.6] | 23.0 | [20.7 - 25.4] | 14.5 | [12.0 - 17.3] | 0.16 |
| **'KL0825'** | *Corynebacterium diphtheriae* NCTC 11397 | 14.2 | [11.4 - 17.6] | 23.0 | [20.7 - 25.4] | 14.5 | [12.0 - 17.3] | 0.21 |
| **'KL0846'** | *Corynebacterium diphtheriae* NCTC 11397 | 14.2 | [11.4 - 17.6] | 23.0 | [20.7 - 25.4] | 14.5 | [12.0 - 17.3] | 0.21 |
| **'KL0541'** | *Corynebacterium phocae* DSM 44612 | 12.9 | [10.2 - 16.2] | 23.0 | [20.7 - 25.5] | 13.3 | [10.9 - 16.0] | 5.44 |
| **'KL0832'** | *Corynebacterium diphtheriae* NCTC 11397 | 14.2 | [11.4 - 17.6] | 23.0 | [20.7 - 25.4] | 14.5 | [12.0 - 17.3] | 0.22 |
| **'KL0515'** | *Corynebacterium diphtheriae* subsp. *lausannense* CHUV2995 | 14.1 | [11.3 - 17.4] | 23.0 | [20.7 - 25.4] | 14.4 | [11.9 - 17.2] | 0.62 |

| Query | Subject | $d_0$ | C.I. $d_0$ | $d_4$ | C.I. $d_4$ | $d_6$ | C.I. $d_6$ | Diff. G+C Percent |
|---|---|---|---|---|---|---|---|---|
| **'KL0870'** | *Corynebacterium diphtheriae* NCTC 11397 | 14.2 | [11.4 - 17.6] | 23.0 | [20.7 - 25.5] | 14.5 | [12.0 - 17.3] | 0.16 |
| **'KL0840'** | *Corynebacterium diphtheriae* NCTC 11397 | 14.2 | [11.4 - 17.6] | 23.0 | [20.7 - 25.4] | 14.5 | [12.0 - 17.3] | 0.21 |
| **'KL0483'** | *Corynebacterium diphtheriae* NCTC 11397 | 14.2 | [11.3 - 17.5] | 23.0 | [20.7 - 25.4] | 14.4 | [12.0 - 17.3] | 0.13 |
| **'KL0556'** | *Corynebacterium diphtheriae* NCTC 11397 | 14.2 | [11.4 - 17.6] | 23.0 | [20.7 - 25.4] | 14.5 | [12.0 - 17.3] | 0.2 |
| **'KL0867'** | *Corynebacterium phocae* DSM 44612 | 12.9 | [10.2 - 16.2] | 23.0 | [20.7 - 25.5] | 13.3 | [10.9 - 16.0] | 5.47 |
| **'KL0796'** | *Corynebacterium diphtheriae* NCTC 11397 | 14.2 | [11.4 - 17.5] | 22.9 | [20.7 - 25.4] | 14.4 | [12.0 - 17.3] | 0.17 |
| **'KL0497'** | *Corynebacterium diphtheriae* NCTC 11397 | 14.2 | [11.4 - 17.6] | 22.9 | [20.6 - 25.4] | 14.5 | [12.0 - 17.3] | 0.22 |
| **'KL0501'** | *Corynebacterium diphtheriae* NCTC 11397 | 14.2 | [11.4 - 17.6] | 22.9 | [20.6 - 25.4] | 14.5 | [12.0 - 17.3] | 0.22 |
| **'KL0870'** | *Corynebacterium diphtheriae* subsp. *lausannense* CHUV2995 | 14.3 | [11.5 - 17.7] | 22.9 | [20.6 - 25.4] | 14.6 | [12.1 - 17.4] | 0.57 |
| **'KL0501'** | *Corynebacterium phocae* DSM 44612 | 12.9 | [10.2 - 16.2] | 22.9 | [20.7 - 25.4] | 13.3 | [10.9 - 16.0] | 5.5 |
| **'KL0785'** | *Corynebacterium diphtheriae* subsp. *lausannense* CHUV2995 | 14.1 | [11.3 - 17.4] | 22.9 | [20.6 - 25.4] | 14.4 | [11.9 - 17.2] | 0.58 |
| **'KL0870'** | *Corynebacterium phocae* DSM 44612 | 12.9 | [10.2 - 16.2] | 22.9 | [20.7 - 25.4] | 13.3 | [10.9 - 16.0] | 5.44 |
| **'KL0846'** | *Corynebacterium phocae* DSM 44612 | 12.9 | [10.2 - 16.2] | 22.9 | [20.6 - 25.4] | 13.3 | [10.9 - 16.0] | 5.5 |
| **'KL0870'** | *Corynebacterium rouxii* FRC0190 T | 14.5 | [11.7 - 17.9] | 22.9 | [20.6 - 25.4] | 14.7 | [12.3 - 17.6] | 0.14 |
| **'KL0818'** | *Corynebacterium diphtheriae* subsp. *lausannense* CHUV2995 | 14.1 | [11.3 - 17.4] | 22.9 | [20.6 - 25.4] | 14.4 | [11.9 - 17.2] | 0.58 |
| **'KL0547'** | *Corynebacterium diphtheriae* NCTC 11397 | 14.2 | [11.4 - 17.6] | 22.9 | [20.6 - 25.4] | 14.5 | [12.0 - 17.3] | 0.18 |
| **'KL0876'** | *Corynebacterium diphtheriae* NCTC 11397 | 14.2 | [11.4 - 17.6] | 22.9 | [20.7 - 25.4] | 14.5 | [12.1 - 17.4] | 0.25 |
| **'KL0497'** | *Corynebacterium phocae* DSM 44612 | 12.9 | [10.2 - 16.2] | 22.8 | [20.5 - 25.3] | 13.3 | [10.9 - 16.0] | 5.51 |
| **'KL0540'** | *Corynebacterium rouxii* FRC0190 T | 14.5 | [11.6 - 17.9] | 22.8 | [20.5 - 25.2] | 14.7 | [12.3 - 17.6] | 0.13 |
| **'KL0541'** | *Corynebacterium diphtheriae* NCTC 11397 | 14.1 | [11.3 - 17.5] | 22.8 | [20.5 - 25.3] | 14.4 | [12.0 - 17.3] | 0.16 |
| **'KL0867'** | *Corynebacterium diphtheriae* NCTC 11397 | 14.2 | [11.4 - 17.6] | 22.8 | [20.5 - 25.3] | 14.5 | [12.0 - 17.3] | 0.19 |
| **'KL0483'** | *Corynebacterium rouxii* FRC0190 T | 14.4 | [11.6 - 17.8] | 22.8 | [20.5 - 25.2] | 14.7 | [12.2 - 17.5] | 0.17 |
| **'KL0796'** | *Corynebacterium rouxii* FRC0190 T | 14.5 | [11.6 - 17.9] | 22.8 | [20.6 - 25.3] | 14.7 | [12.3 - 17.6] | 0.12 |
| **'KL0483'** | *Corynebacterium phocae* DSM 44612 | 12.9 | [10.2 - 16.2] | 22.8 | [20.5 - 25.2] | 13.3 | [10.9 - 16.0] | 5.41 |
| **'KL0540'** | *Corynebacterium phocae* DSM 44612 | 12.9 | [10.2 - 16.2] | 22.8 | [20.6 - 25.3] | 13.3 | [10.9 - 16.0] | 5.45 |
| **'KL0497'** | *Corynebacterium rouxii* FRC0190 T | 14.4 | [11.6 - 17.8] | 22.7 | [20.5 - 25.2] | 14.7 | [12.2 - 17.5] | 0.08 |

| Query | Subject | $d_0$ | C.I. $d_0$ | $d_4$ | C.I. $d_4$ | $d_6$ | C.I. $d_6$ | Diff. G+C Percent |
|---|---|---|---|---|---|---|---|---|
| **'KL0832'** | *Corynebacterium phocae* DSM 44612 | 12.9 | [10.2 - 16.2] | 22.7 | [20.4 - 25.2] | 13.3 | [10.9 - 16.1] | 5.5 |
| **'KL0556'** | *Corynebacterium rouxii* FRC0190 T | 14.3 | [11.5 - 17.7] | 22.7 | [20.4 - 25.1] | 14.6 | [12.1 - 17.4] | 0.09 |
| **'KL0796'** | *Corynebacterium phocae* DSM 44612 | 12.9 | [10.2 - 16.2] | 22.7 | [20.5 - 25.2] | 13.3 | [10.9 - 16.0] | 5.46 |
| **'KL0541'** | *Corynebacterium rouxii* FRC0190 T | 14.6 | [11.7 - 18.0] | 22.7 | [20.4 - 25.1] | 14.8 | [12.3 - 17.7] | 0.14 |
| **'KL0515'** | *Corynebacterium rouxii* FRC0190 T | 14.3 | [11.5 - 17.7] | 22.7 | [20.4 - 25.1] | 14.6 | [12.1 - 17.4] | 0.1 |
| **'KL0825'** | *Corynebacterium phocae* DSM 44612 | 12.9 | [10.2 - 16.2] | 22.7 | [20.5 - 25.2] | 13.3 | [10.9 - 16.0] | 5.5 |
| **'KL0785'** | *Corynebacterium rouxii* FRC0190 T | 14.3 | [11.5 - 17.7] | 22.7 | [20.5 - 25.2] | 14.5 | [12.1 - 17.4] | 0.14 |
| **'KL0818'** | *Corynebacterium rouxii* FRC0190 T | 14.3 | [11.5 - 17.7] | 22.7 | [20.4 - 25.2] | 14.5 | [12.1 - 17.4] | 0.13 |
| **'KL0840'** | *Corynebacterium phocae* DSM 44612 | 12.9 | [10.2 - 16.2] | 22.7 | [20.4 - 25.2] | 13.3 | [10.9 - 16.1] | 5.5 |
| **'KL0547'** | *Corynebacterium rouxii* FRC0190 T | 14.5 | [11.7 - 17.9] | 22.6 | [20.3 - 25.1] | 14.8 | [12.3 - 17.6] | 0.11 |
| **'KL0853'** | *Corynebacterium diphtheriae* subsp. *lausannense* CHUV2995 | 14.1 | [11.3 - 17.5] | 22.6 | [20.3 - 25.0] | 14.4 | [12.0 - 17.2] | 0.53 |
| **'KL0853'** | *Corynebacterium rouxii* FRC0190 T | 14.4 | [11.6 - 17.8] | 22.6 | [20.3 - 25.0] | 14.7 | [12.2 - 17.5] | 0.18 |
| **'KL0547'** | *Corynebacterium diphtheriae* subsp. *lausannense* CHUV2995 | 14.2 | [11.3 - 17.5] | 22.6 | [20.3 - 25.0] | 14.4 | [12.0 - 17.3] | 0.6 |
| **'KL0796'** | *Corynebacterium diphtheriae* subsp. *lausannense* CHUV2995 | 14.2 | [11.4 - 17.6] | 22.6 | [20.3 - 25.1] | 14.5 | [12.1 - 17.3] | 0.59 |
| **'KL0867'** | *Corynebacterium rouxii* FRC0190 T | 14.4 | [11.6 - 17.8] | 22.6 | [20.3 - 25.0] | 14.7 | [12.2 - 17.5] | 0.11 |
| **'KL0501'** | *Corynebacterium rouxii* FRC0190 T | 14.4 | [11.6 - 17.8] | 22.5 | [20.2 - 24.9] | 14.6 | [12.2 - 17.5] | 0.08 |
| **'KL0556'** | *Corynebacterium diphtheriae* subsp. *lausannense* CHUV2995 | 14.2 | [11.3 - 17.5] | 22.5 | [20.2 - 24.9] | 14.4 | [12.0 - 17.3] | 0.62 |
| **'KL0846'** | *Corynebacterium rouxii* FRC0190 T | 14.4 | [11.6 - 17.8] | 22.5 | [20.3 - 25.0] | 14.7 | [12.2 - 17.5] | 0.09 |
| **'KL0497'** | *Corynebacterium diphtheriae* subsp. *lausannense* CHUV2995 | 14.2 | [11.4 - 17.6] | 22.5 | [20.2 - 24.9] | 14.5 | [12.0 - 17.3] | 0.64 |
| **'KL0832'** | *Corynebacterium rouxii* FRC0190 T | 14.4 | [11.6 - 17.8] | 22.5 | [20.2 - 25.0] | 14.7 | [12.2 - 17.5] | 0.08 |
| **'KL0840'** | *Corynebacterium rouxii* FRC0190 T | 14.4 | [11.6 - 17.8] | 22.5 | [20.3 - 25.0] | 14.7 | [12.2 - 17.5] | 0.08 |
| **'KL0825'** | *Corynebacterium rouxii* FRC0190 T | 14.4 | [11.6 - 17.8] | 22.5 | [20.2 - 25.0] | 14.6 | [12.2 - 17.5] | 0.08 |
| **'KL0870'** | *Corynebacterium belfantii* FRC0043 | 14.3 | [11.5 - 17.7] | 22.5 | [20.2 - 25.0] | 14.6 | [12.1 - 17.4] | 0.26 |
| **'KL0867'** | *Corynebacterium diphtheriae* subsp. *lausannense* CHUV2995 | 14.2 | [11.3 - 17.5] | 22.5 | [20.2 - 24.9] | 14.4 | [12.0 - 17.3] | 0.6 |
| **'KL0501'** | *Corynebacterium diphtheriae* subsp. *lausannense* CHUV2995 | 14.2 | [11.3 - 17.5] | 22.4 | [20.1 - 24.8] | 14.4 | [12.0 - 17.3] | 0.63 |

| Query | Subject | $d_0$ | C.I. $d_0$ | $d_4$ | C.I. $d_4$ | $d_6$ | C.I. $d_6$ | Diff. G+C Percent |
|---|---|---|---|---|---|---|---|---|
| **'KL0515'** | *Corynebacterium belfantii* FRC0043 | 14.1 | [11.3 - 17.5] | 22.4 | [20.1 - 24.8] | 14.4 | [12.0 - 17.2] | 0.3 |
| **'KL0483'** | *Corynebacterium diphtheriae* subsp. *lausannense* CHUV2995 | 14.2 | [11.4 - 17.6] | 22.4 | [20.1 - 24.8] | 14.5 | [12.0 - 17.3] | 0.54 |
| **'KL0840'** | *Corynebacterium diphtheriae* subsp. *lausannense* CHUV2995 | 14.1 | [11.3 - 17.5] | 22.4 | [20.2 - 24.9] | 14.4 | [12.0 - 17.3] | 0.63 |
| **'KL0541'** | *Corynebacterium diphtheriae* subsp. *lausannense* CHUV2995 | 14.2 | [11.4 - 17.6] | 22.4 | [20.1 - 24.9] | 14.5 | [12.0 - 17.3] | 0.58 |
| **'KL0818'** | *Corynebacterium belfantii* FRC0043 | 14.1 | [11.3 - 17.5] | 22.4 | [20.1 - 24.8] | 14.4 | [12.0 - 17.2] | 0.27 |
| **'KL0825'** | *Corynebacterium diphtheriae* subsp. *lausannense* CHUV2995 | 14.1 | [11.3 - 17.5] | 22.4 | [20.2 - 24.9] | 14.4 | [12.0 - 17.2] | 0.63 |
| **'KL0832'** | *Corynebacterium diphtheriae* subsp. *lausannense* CHUV2995 | 14.1 | [11.3 - 17.5] | 22.4 | [20.2 - 24.9] | 14.4 | [12.0 - 17.3] | 0.63 |
| **'KL0846'** | *Corynebacterium diphtheriae* subsp. *lausannense* CHUV2995 | 14.2 | [11.3 - 17.5] | 22.4 | [20.2 - 24.9] | 14.4 | [12.0 - 17.3] | 0.63 |
| **'KL0785'** | *Corynebacterium belfantii* FRC0043 | 14.1 | [11.3 - 17.5] | 22.4 | [20.1 - 24.8] | 14.4 | [12.0 - 17.2] | 0.26 |
| **'KL0876'** | *Corynebacterium rouxii* FRC0190 T | 14.4 | [11.5 - 17.7] | 22.4 | [20.2 - 24.9] | 14.6 | [12.2 - 17.4] | 0.05 |
| **'KL0876'** | *Corynebacterium diphtheriae* subsp. *lausannense* CHUV2995 | 14.2 | [11.4 - 17.5] | 22.4 | [20.1 - 24.9] | 14.4 | [12.0 - 17.3] | 0.66 |
| **'KL0540'** | *Corynebacterium diphtheriae* subsp. *lausannense* CHUV2995 | 14.2 | [11.4 - 17.6] | 22.3 | [20.1 - 24.8] | 14.5 | [12.1 - 17.3] | 0.58 |
| **'KL0853'** | *Corynebacterium belfantii* FRC0043 | 14.2 | [11.3 - 17.5] | 22.2 | [19.9 - 24.6] | 14.4 | [12.0 - 17.3] | 0.22 |
| **'KL0796'** | *Corynebacterium belfantii* FRC0043 | 14.3 | [11.4 - 17.6] | 22.1 | [19.8 - 24.5] | 14.5 | [12.1 - 17.4] | 0.27 |
| **'KL0825'** | *Corynebacterium belfantii* FRC0043 | 14.2 | [11.3 - 17.5] | 22.0 | [19.7 - 24.4] | 14.4 | [12.0 - 17.3] | 0.31 |
| **'KL0547'** | *Corynebacterium belfantii* FRC0043 | 14.2 | [11.4 - 17.6] | 22.0 | [19.7 - 24.4] | 14.5 | [12.0 - 17.3] | 0.28 |
| **'KL0832'** | *Corynebacterium belfantii* FRC0043 | 14.2 | [11.4 - 17.6] | 22.0 | [19.7 - 24.4] | 14.4 | [12.0 - 17.3] | 0.32 |
| **'KL0876'** | *Corynebacterium belfantii* FRC0043 | 14.2 | [11.4 - 17.6] | 22.0 | [19.7 - 24.4] | 14.5 | [12.0 - 17.3] | 0.35 |
| **'KL0840'** | *Corynebacterium belfantii* FRC0043 | 14.2 | [11.4 - 17.6] | 22.0 | [19.7 - 24.4] | 14.4 | [12.0 - 17.3] | 0.31 |
| **'KL0846'** | *Corynebacterium belfantii* FRC0043 | 14.2 | [11.4 - 17.6] | 22.0 | [19.7 - 24.4] | 14.4 | [12.0 - 17.3] | 0.31 |
| **'KL0867'** | *Corynebacterium belfantii* FRC0043 | 14.2 | [11.4 - 17.6] | 21.9 | [19.7 - 24.4] | 14.5 | [12.0 - 17.3] | 0.29 |
| **'KL0501'** | *Corynebacterium belfantii* FRC0043 | 14.2 | [11.4 - 17.6] | 21.9 | [19.7 - 24.4] | 14.4 | [12.0 - 17.3] | 0.32 |
| **'KL0483'** | *Corynebacterium belfantii* FRC0043 | 14.2 | [11.4 - 17.6] | 21.9 | [19.7 - 24.4] | 14.5 | [12.0 - 17.3] | 0.23 |
| **'KL0541'** | *Corynebacterium belfantii* FRC0043 | 14.2 | [11.4 - 17.6] | 21.9 | [19.7 - 24.4] | 14.5 | [12.0 - 17.3] | 0.26 |
| **'KL0556'** | *Corynebacterium belfantii* FRC0043 | 14.2 | [11.4 - 17.6] | 21.9 | [19.7 - 24.4] | 14.5 | [12.0 - 17.3] | 0.3 |

| Query | Subject | $d_0$ | C.I. $d_0$ | $d_4$ | C.I. $d_4$ | $d_6$ | C.I. $d_6$ | Diff. G+C Percent |
|---|---|---|---|---|---|---|---|---|
| **'KL0475'** | *Corynebacterium diphtheriae* NCTC 11397 | 13.9 | [11.1 - 17.3] | 21.9 | [19.6 - 24.3] | 14.2 | [11.8 - 17.0] | 0.2 |
| **'KL0497'** | *Corynebacterium belfantii* FRC0043 | 14.2 | [11.4 - 17.6] | 21.9 | [19.7 - 24.4] | 14.5 | [12.1 - 17.3] | 0.32 |
| **'KL0540'** | *Corynebacterium belfantii* FRC0043 | 14.3 | [11.5 - 17.7] | 21.8 | [19.6 - 24.3] | 14.5 | [12.1 - 17.4] | 0.27 |
| **'KL0475'** | *Corynebacterium diphtheriae* subsp. *lausannense* CHUV2995 | 14.0 | [11.2 - 17.3] | 21.7 | [19.4 - 24.1] | 14.2 | [11.8 - 17.1] | 0.62 |
| **'KL0475'** | *Corynebacterium rouxii* FRC0190 T | 14.2 | [11.4 - 17.6] | 21.6 | [19.3 - 24.0] | 14.4 | [12.0 - 17.3] | 0.09 |
| **'KL0475'** | *Corynebacterium belfantii* FRC0043 | 14.0 | [11.2 - 17.3] | 21.0 | [18.7 - 23.4] | 14.2 | [11.8 - 17.1] | 0.31 |

| Strain | Authority | Other deposits | Synonyms | Base pairs | Percent G+C | No. proteins | Goldstamp | Bioproject accession | Biosample accession | Assembly accession | IMG OID |
|---|---|---|---|---|---|---|---|---|---|---|---|
| *Corynebacterium kutscheri* DSM 20755 | (Migula 1900) Bergey et al. 1925 emend. Nouioui et al. 2018 | CCUG 27535; ATCC 15677; NCTC 11138; JCM 9385; IFO 15288; NBRC 15288; CIP 103423 | *Bacterium kutscheri*; *Corynebacterium kutscheri* | 2354 065 | 46.5 | 2047 | Gp0110293 | PRJNA276037 | SAMN03365283 | GCA_000980835 | |
| *Corynebacterium mustelae* DSM 45274 | Funke et al. 2010 emend. Nouioui et al. 2018 | 3105; CCUG 57279 | *Corynebacterium mustelae* | 3474 226 | 52.6 | 3110 | Gp0114696 | PRJNA282348 | SAMN03568800 | GCA_001020985 | |
| *Corynebacterium diphtheriae* NCTC 11397 | (Kruse 1886) Lehmann and Neumann 1896 emend. Nouioui et al. 2018 | DSM 44123; ATCC 27010; CIP 100721 | *Bacillus diphtheriae*; *Corynebacterium diphtheriae*; *Corynebacterium diphtheriae* subsp. *diphtheriae* | 2463 666 | 53.5 | 2337 | Gp0132011 | PRJEB6403 | SAMEA2517360 | GCA_001457455 | |
| KL0475 | | | | 2472 849 | 53.3 | 2279 | | | | | |
| KL0483 | | | | 2577 295 | 53.4 | 2395 | | | | | |
| KL0497 | | | | 2477 500 | 53.3 | 2261 | | | | | |
| KL0501 | | | | 2446 525 | 53.3 | 2221 | | | | | |
| KL0515 | | | | 2449 403 | 53.3 | 2212 | | | | | |
| KL0540 | | | | 2474 507 | 53.4 | 2254 | | | | | |
| KL0541 | | | | 2640 162 | 53.4 | 2485 | | | | | |
| KL0547 | | | | 2549 371 | 53.3 | 2364 | | | | | |
| KL0556 | | | | 2421 288 | 53.3 | 2169 | | | | | |
| KL0785 | | | | 2470 695 | 53.4 | 2237 | | | | | |

| Strain | Authority | Other deposits | Synonyms | Base pairs | Percent G+C | No. proteins | Goldstamp | Bioproject accession | Biosample accession | Assembly accession | IMG OID |
|---|---|---|---|---|---|---|---|---|---|---|---|
| KL0796 | | | | 2603 053 | 53.4 | 2435 | | | | | |
| KL0818 | | | | 2474 549 | 53.4 | 2245 | | | | | |
| KL0825 | | | | 2499 718 | 53.3 | 2289 | | | | | |
| KL0832 | | | | 2452 482 | 53.3 | 2217 | | | | | |
| KL0840 | | | | 2449 355 | 53.3 | 2212 | | | | | |
| KL0846 | | | | 2443 613 | 53.3 | 2205 | | | | | |
| KL0853 | | | | 2557 514 | 53.4 | 2343 | | | | | |
| KL0867 | | | | 2514 118 | 53.3 | 2319 | | | | | |
| KL0870 | | | | 2575 028 | 53.4 | 2385 | | | | | |
| KL0876 | | | | 2469 953 | 53.3 | 2231 | | | | | |

#### Results

##### Type-based species and subspecies clustering

The resulting species and subspecies clusters are listed in Table 4, whereas the taxonomic identification of the query strains is found in Table 1. Briefly, the clustering yielded 12 species clusters and the provided query strains were assigned to 1 of these. Moreover, user strains were located in 1 of 12 subspecies clusters.

##### Figure caption genome tree

**Figure 1.** Tree inferred with FastME 2.1.6.1 [6] from GBDP distances calculated from genome sequences. The branch lengths are scaled in terms of GBDP distance formula $d_5$. The numbers above branches are GBDP pseudo-bootstrap support values > 60 % from 100 replications, with an average branch support of 26.8 %. The tree was rooted at the midpoint [7].

##### Figure caption SSU tree

**Figure 2.** Tree inferred with FastME 2.1.6.1 [6] from GBDP distances calculated from 16S rDNA gene sequences. The branch lengths are scaled in terms of GBDP distance formula $d_5$. The numbers above branches are GBDP pseudo-bootstrap support values > 60 % from 100 replications, with an average branch support of 43.4 %. The tree was rooted at the midpoint [7].
