## Supplementary File 3 for "Taxonomic classification of strain PO100/5 shows a broader geographic distribution and genetic markers of the recently described *Corynebacterium silvaticum*": Cul_public_p1_report.pdf

### TYPE (STRAIN) GENOME SERVER

### JOB SUMMARY

PRINT DATE: 2020-06-11 22:44:46 +0200

JOB ID: 8293e2dc-bebd-425a-ab0d-63c32b106f85

RESULT PAGE: https://tygs.dsmz.de/user_results/show?guid=8293e2dc-bebd-425a-ab0d-63c32b106f85

#### Table 1: Phylogenies

**Publication-ready versions** of both the genome-scale GBDP tree and the 16S rRNA gene sequence tree can be customized and exported either in SVG (vector graphic) or PNG format from within the phylogeny viewers in your TYGS result page. For publications the **SVG format is recommended** because it is lossless, always keeps its high resolution and can also be easily converted to other popular formats such as PDF or EPS. Please follow the link provided above!

#### Table 2: Identification

The below list contains the result of the TYGS species identification routine.

Explanation of remarks that might occur in the below table:

**remark [R1]:** The TYGS type strain database is automatically updated on an almost daily basis. However, if a particular type strain genome is not available in the TYGS database, this can have several reasons which are detailed in the FAQ. You can request an extended 16S rRNA gene analysis via the 16S tree viewer found in your result page to detect **not yet genome-sequenced** type strains relevant for your study.

**remark [R2]:** > 70% dDDH value (formula $d_4$) and (almost) minimal dDDH values for gene-content formulae $d_0$ and $d_6$ indicate a potentially unreliable identification result and should thus be checked via the 16S rRNA gene sequence similarity. Such strong deviations can, in principle, be caused by sequence contamination.

**remark [R3]:** G+C content difference of > 1 % indicates a potentially unreliable identification result because within species G+C content varies no more than 1 %, if computed from genome sequences (PMID: 24505073).

| Strain | Conclusion | Identification result | Remark |
|---|---|---|---|
| **'04-13'** | belongs to known species | *Corynebacterium silvaticum* | |
| **'05-13'** | belongs to known species | *Corynebacterium silvaticum* | |
| **'02-13'** | belongs to known species | *Corynebacterium ulcerans* | |
| **'04-15'** | belongs to known species | *Corynebacterium ulcerans* | |
| **'04-3911'** | belongs to known species | *Corynebacterium ulcerans* | |
| **'06-16'** | belongs to known species | *Corynebacterium ulcerans* | |
| **'06-19'** | belongs to known species | *Corynebacterium ulcerans* | |
| **'0102'** | belongs to known species | *Corynebacterium ulcerans* | |
| **'0211'** | belongs to known species | *Corynebacterium ulcerans* | |
| **'809'** | belongs to known species | *Corynebacterium ulcerans* | |
| **'2590'** | belongs to known species | *Corynebacterium ulcerans* | |
| **'4940'** | belongs to known species | *Corynebacterium ulcerans* | |
| **'05146'** | belongs to known species | *Corynebacterium ulcerans* | |
| **'131001'** | belongs to known species | *Corynebacterium ulcerans* | |
| **'210931'** | belongs to known species | *Corynebacterium ulcerans* | |

| Strain | Conclusion | Identification result | Remark |
|---|---|---|---|
| **'210932'** | belongs to known species | *Corynebacterium ulcerans* | |
| **'BR-AD22'** | belongs to known species | *Corynebacterium ulcerans* | |
| **'03-8664'** | potential new species | | see [R1] |
| **'04-7514'** | potential new species | | see [R1] |
| **'131002'** | potential new species | | see [R1] |

#### Table 3: Pairwise comparisons of user genomes vs. type-strain genomes

The following table contains the pairwise dDDH values between your user genomes and the selected type-strain genomes. The dDDH values are provided along with their confidence intervals (C.I.) for the three different GBDP formulas:

• formula $d_0$ (a.k.a. GGDC formula 1): length of all HSPs divided by total genome length
• formula $d_4$ (a.k.a. GGDC formula 2): sum of all identities found in HSPs divided by overall HSP length
• formula $d_6$ (a.k.a. GGDC formula 3): sum of all identities found in HSPs divided by total genome length

**Note:** Formula $d_4$ is independent of genome length and is thus robust against the use of incomplete draft genomes. For other reasons for preferring formula $d_4$, see the FAQ.

| Query | Subject | $d_0$ | C.I. $d_0$ | $d_4$ | C.I. $d_4$ | $d_6$ | C.I. $d_6$ | Diff. G+C Percent |
|---|---|---|---|---|---|---|---|---|
| **'04-15'** | **'06-19'** | 100.0 | [100.0 - 100.0] | 100.0 | [100.0 - 100.0] | 100.0 | [100.0 - 100.0] | 0.0 |
| **'04-13'** | *Corynebacterium silvaticum* KL0182 | 100.0 | [100.0 - 100.0] | 99.9 | [99.8 - 99.9] | 100.0 | [100.0 - 100.0] | 0.01 |
| **'0102'** | **'0211'** | 100.0 | [100.0 - 100.0] | 99.9 | [99.8 - 99.9] | 100.0 | [100.0 - 100.0] | 0.0 |
| **'131001'** | **'210932'** | 100.0 | [100.0 - 100.0] | 99.8 | [99.7 - 99.9] | 100.0 | [100.0 - 100.0] | 0.0 |
| **'02-13'** | **'06-16'** | 99.5 | [99.1 - 99.8] | 99.6 | [99.3 - 99.7] | 99.8 | [99.6 - 99.9] | 0.03 |
| **'04-3911'** | **'0211'** | 98.9 | [98.0 - 99.4] | 99.5 | [99.1 - 99.7] | 99.5 | [99.0 - 99.7] | 0.04 |
| **'04-3911'** | **'0102'** | 98.9 | [98.0 - 99.4] | 99.5 | [99.1 - 99.7] | 99.5 | [99.0 - 99.7] | 0.04 |
| **'04-13'** | **'05-13'** | 99.7 | [99.4 - 99.9] | 98.6 | [97.9 - 99.1] | 99.9 | [99.7 - 99.9] | 0.02 |
| **'05-13'** | *Corynebacterium silvaticum* KL0182 | 99.7 | [99.4 - 99.9] | 98.6 | [97.9 - 99.0] | 99.8 | [99.7 - 99.9] | 0.03 |
| **'06-16'** | **'210931'** | 99.4 | [98.8 - 99.7] | 98.6 | [98.0 - 99.1] | 99.7 | [99.4 - 99.8] | 0.07 |
| **'02-13'** | **'210931'** | 98.4 | [97.2 - 99.0] | 98.4 | [97.7 - 98.9] | 99.1 | [98.5 - 99.5] | 0.04 |
| **'06-19'** | **'BR-AD22'** | 98.0 | [96.7 - 98.8] | 98.0 | [97.2 - 98.6] | 98.9 | [98.2 - 99.3] | 0.07 |
| **'04-15'** | **'BR-AD22'** | 98.1 | [96.8 - 98.8] | 97.9 | [97.0 - 98.5] | 98.9 | [98.2 - 99.3] | 0.08 |
| **'06-16'** | **'05146'** | 98.7 | [97.7 - 99.3] | 96.7 | [95.5 - 97.6] | 99.2 | [98.6 - 99.5] | 0.07 |
| **'02-13'** | **'05146'** | 98.7 | [97.7 - 99.3] | 96.5 | [95.2 - 97.4] | 99.2 | [98.6 - 99.5] | 0.03 |
| **'05146'** | **'210931'** | 99.2 | [98.5 - 99.6] | 95.6 | [94.2 - 96.8] | 99.4 | [99.0 - 99.7] | 0.0 |
| **'04-15'** | **'04-3911'** | 98.5 | [97.4 - 99.1] | 91.9 | [89.8 - 93.6] | 98.7 | [97.9 - 99.2] | 0.0 |
| **'04-3911'** | **'06-19'** | 98.4 | [97.3 - 99.1] | 91.9 | [89.8 - 93.6] | 98.7 | [97.9 - 99.2] | 0.01 |
| **'04-3911'** | **'BR-AD22'** | 95.7 | [93.6 - 97.2] | 91.4 | [89.2 - 93.1] | 96.8 | [95.4 - 97.8] | 0.08 |
| **'04-15'** | **'0102'** | 96.8 | [95.1 - 98.0] | 91.3 | [89.2 - 93.1] | 97.6 | [96.3 - 98.4] | 0.03 |
| **'04-15'** | **'0211'** | 96.8 | [95.1 - 98.0] | 91.3 | [89.2 - 93.1] | 97.6 | [96.3 - 98.4] | 0.03 |
| **'06-19'** | **'0211'** | 96.8 | [95.0 - 97.9] | 91.3 | [89.2 - 93.1] | 97.6 | [96.3 - 98.4] | 0.03 |

| Query | Subject | $d_0$ | C.I. $d_0$ | $d_4$ | C.I. $d_4$ | $d_6$ | C.I. $d_6$ | Diff. G+C Percent |
|---|---|---|---|---|---|---|---|---|
| **'06-19'** | **'0102'** | 96.8 | [95.0 - 97.9] | 91.3 | [89.1 - 93.1] | 97.6 | [96.3 - 98.4] | 0.03 |
| **'2590'** | **'4940'** | 96.9 | [95.2 - 98.0] | 90.5 | [88.3 - 92.4] | 97.6 | [96.3 - 98.4] | 0.05 |
| **'04-3911'** | **'05146'** | 98.8 | [97.8 - 99.3] | 90.3 | [88.0 - 92.2] | 98.8 | [98.1 - 99.3] | 0.02 |
| **'2590'** | *Corynebacterium ulcerans* NCTC 7910 | 96.8 | [95.0 - 97.9] | 90.1 | [87.8 - 92.0] | 97.4 | [96.1 - 98.3] | 0.02 |
| **'0211'** | **'BR-AD22'** | 97.8 | [96.4 - 98.7] | 90.1 | [87.8 - 92.0] | 98.1 | [97.1 - 98.8] | 0.04 |
| **'0102'** | **'BR-AD22'** | 97.8 | [96.4 - 98.7] | 90.1 | [87.8 - 92.0] | 98.1 | [97.1 - 98.8] | 0.04 |
| **'809'** | **'4940'** | 98.7 | [97.7 - 99.2] | 89.8 | [87.4 - 91.7] | 98.7 | [97.9 - 99.2] | 0.04 |
| **'0211'** | **'05146'** | 96.8 | [95.0 - 97.9] | 89.3 | [86.9 - 91.3] | 97.3 | [96.0 - 98.2] | 0.05 |
| **'0102'** | **'05146'** | 96.8 | [95.0 - 97.9] | 89.3 | [86.9 - 91.3] | 97.3 | [96.0 - 98.2] | 0.05 |
| **'809'** | **'2590'** | 95.5 | [93.3 - 97.0] | 89.2 | [86.8 - 91.2] | 96.4 | [94.9 - 97.5] | 0.01 |
| **'4940'** | *Corynebacterium ulcerans* NCTC 7910 | 98.9 | [98.0 - 99.4] | 89.1 | [86.7 - 91.1] | 98.8 | [98.1 - 99.3] | 0.03 |
| **'02-13'** | *Corynebacterium ulcerans* NCTC 7910 | 96.8 | [95.1 - 98.0] | 88.9 | [86.5 - 91.0] | 97.4 | [96.1 - 98.2] | 0.03 |
| **'06-16'** | **'809'** | 96.4 | [94.5 - 97.7] | 88.8 | [86.4 - 90.9] | 97.0 | [95.6 - 98.0] | 0.07 |
| **'06-19'** | **'2590'** | 96.3 | [94.4 - 97.6] | 88.8 | [86.4 - 90.9] | 97.0 | [95.6 - 97.9] | 0.03 |
| **'04-15'** | **'2590'** | 96.3 | [94.4 - 97.6] | 88.8 | [86.4 - 90.9] | 97.0 | [95.6 - 98.0] | 0.03 |
| **'210931'** | *Corynebacterium ulcerans* NCTC 7910 | 97.8 | [96.3 - 98.6] | 88.7 | [86.2 - 90.7] | 98.0 | [96.9 - 98.7] | 0.01 |
| **'06-16'** | *Corynebacterium ulcerans* NCTC 7910 | 97.0 | [95.3 - 98.1] | 88.7 | [86.3 - 90.8] | 97.5 | [96.2 - 98.3] | 0.06 |
| **'809'** | **'210931'** | 97.4 | [95.9 - 98.4] | 88.6 | [86.1 - 90.6] | 97.7 | [96.5 - 98.5] | 0.0 |
| **'809'** | *Corynebacterium ulcerans* NCTC 7910 | 98.3 | [97.1 - 99.0] | 88.5 | [86.0 - 90.6] | 98.3 | [97.4 - 99.0] | 0.01 |
| **'04-3911'** | **'210932'** | 96.7 | [94.9 - 97.9] | 88.5 | [86.0 - 90.5] | 97.2 | [95.9 - 98.1] | 0.01 |
| **'04-15'** | *Corynebacterium ulcerans* NCTC 7910 | 98.2 | [97.0 - 98.9] | 88.5 | [86.0 - 90.6] | 98.3 | [97.3 - 98.9] | 0.01 |
| **'04-3911'** | **'131001'** | 96.7 | [94.9 - 97.9] | 88.5 | [86.0 - 90.5] | 97.2 | [95.9 - 98.1] | 0.01 |
| **'06-19'** | *Corynebacterium ulcerans* NCTC 7910 | 98.2 | [97.0 - 98.9] | 88.5 | [86.0 - 90.5] | 98.3 | [97.3 - 98.9] | 0.02 |
| **'02-13'** | **'809'** | 96.5 | [94.6 - 97.7] | 88.4 | [85.9 - 90.5] | 97.0 | [95.7 - 98.0] | 0.04 |
| **'06-19'** | **'4940'** | 98.8 | [97.9 - 99.3] | 88.4 | [85.9 - 90.5] | 98.7 | [97.9 - 99.2] | 0.01 |
| **'04-15'** | **'4940'** | 98.9 | [98.0 - 99.4] | 88.4 | [85.9 - 90.5] | 98.7 | [98.0 - 99.2] | 0.02 |
| **'2590'** | **'BR-AD22'** | 94.7 | [92.3 - 96.4] | 88.3 | [85.8 - 90.4] | 95.8 | [94.0 - 97.0] | 0.11 |

| Query | Subject | $d_0$ | C.I. $d_0$ | $d_4$ | C.I. $d_4$ | $d_6$ | C.I. $d_6$ | Diff. G+C Percent |
|---|---|---|---|---|---|---|---|---|
| **'4940'** | **'BR-AD22'** | 95.5 | [93.3 - 97.0] | 88.3 | [85.9 - 90.4] | 96.3 | [94.7 - 97.5] | 0.06 |
| **'04-3911'** | *Corynebacterium ulcerans* NCTC 7910 | 97.2 | [95.5 - 98.2] | 88.3 | [85.9 - 90.4] | 97.5 | [96.3 - 98.4] | 0.01 |
| **'809'** | **'BR-AD22'** | 96.7 | [95.0 - 97.9] | 88.2 | [85.7 - 90.3] | 97.2 | [95.9 - 98.1] | 0.1 |
| **'05146'** | *Corynebacterium ulcerans* NCTC 7910 | 98.4 | [97.2 - 99.0] | 88.2 | [85.7 - 90.3] | 98.4 | [97.4 - 99.0] | 0.01 |
| **'02-13'** | **'04-3911'** | 96.5 | [94.6 - 97.7] | 88.2 | [85.7 - 90.3] | 97.0 | [95.6 - 98.0] | 0.02 |
| **'BR-AD22'** | *Corynebacterium ulcerans* NCTC 7910 | 94.8 | [92.5 - 96.5] | 88.2 | [85.7 - 90.3] | 95.8 | [94.1 - 97.1] | 0.09 |
| **'04-15'** | **'809'** | 98.1 | [96.8 - 98.9] | 88.2 | [85.7 - 90.3] | 98.2 | [97.2 - 98.8] | 0.02 |
| **'06-19'** | **'809'** | 98.1 | [96.8 - 98.8] | 88.1 | [85.6 - 90.3] | 98.1 | [97.1 - 98.8] | 0.03 |
| **'2590'** | **'210932'** | 95.6 | [93.5 - 97.1] | 88.0 | [85.4 - 90.1] | 96.4 | [94.8 - 97.5] | 0.02 |
| **'0211'** | *Corynebacterium ulcerans* NCTC 7910 | 95.1 | [92.8 - 96.7] | 88.0 | [85.5 - 90.1] | 96.0 | [94.4 - 97.2] | 0.04 |
| **'131001'** | *Corynebacterium ulcerans* NCTC 7910 | 97.2 | [95.6 - 98.3] | 88.0 | [85.5 - 90.1] | 97.5 | [96.3 - 98.4] | 0.0 |
| **'2590'** | **'131001'** | 95.7 | [93.5 - 97.1] | 88.0 | [85.5 - 90.1] | 96.4 | [94.8 - 97.5] | 0.02 |
| **'0102'** | *Corynebacterium ulcerans* NCTC 7910 | 95.1 | [92.8 - 96.7] | 88.0 | [85.5 - 90.1] | 96.0 | [94.4 - 97.2] | 0.05 |
| **'04-3911'** | **'2590'** | 95.9 | [93.8 - 97.3] | 87.9 | [85.4 - 90.1] | 96.6 | [95.0 - 97.6] | 0.03 |
| **'809'** | **'05146'** | 97.9 | [96.6 - 98.8] | 87.9 | [85.3 - 90.0] | 98.0 | [97.0 - 98.7] | 0.0 |
| **'04-3911'** | **'06-16'** | 96.9 | [95.1 - 98.0] | 87.9 | [85.4 - 90.0] | 97.3 | [96.0 - 98.2] | 0.05 |
| **'04-3911'** | **'210931'** | 97.7 | [96.3 - 98.6] | 87.9 | [85.4 - 90.0] | 97.9 | [96.8 - 98.6] | 0.02 |
| **'210932'** | *Corynebacterium ulcerans* NCTC 7910 | 97.3 | [95.6 - 98.3] | 87.9 | [85.4 - 90.1] | 97.6 | [96.3 - 98.4] | 0.0 |
| **'0102'** | **'131001'** | 96.9 | [95.2 - 98.0] | 87.8 | [85.3 - 89.9] | 97.3 | [96.0 - 98.2] | 0.05 |
| **'0102'** | **'210932'** | 96.9 | [95.2 - 98.0] | 87.8 | [85.3 - 89.9] | 97.3 | [96.0 - 98.2] | 0.04 |
| **'0211'** | **'210932'** | 96.9 | [95.2 - 98.0] | 87.8 | [85.3 - 89.9] | 97.3 | [96.0 - 98.2] | 0.04 |
| **'0211'** | **'131001'** | 96.9 | [95.2 - 98.0] | 87.8 | [85.3 - 89.9] | 97.3 | [96.0 - 98.2] | 0.05 |
| **'02-13'** | **'2590'** | 95.0 | [92.7 - 96.6] | 87.7 | [85.2 - 89.9] | 95.9 | [94.2 - 97.2] | 0.04 |
| **'4940'** | **'210931'** | 98.4 | [97.2 - 99.0] | 87.6 | [85.1 - 89.8] | 98.3 | [97.4 - 98.9] | 0.04 |
| **'04-15'** | **'06-16'** | 97.3 | [95.7 - 98.3] | 87.6 | [85.1 - 89.8] | 97.5 | [96.3 - 98.4] | 0.05 |
| **'06-19'** | **'05146'** | 98.8 | [97.9 - 99.3] | 87.6 | [85.1 - 89.8] | 98.7 | [97.8 - 99.2] | 0.02 |
| **'04-15'** | **'210931'** | 97.8 | [96.5 - 98.7] | 87.6 | [85.1 - 89.8] | 97.9 | [96.8 - 98.7] | 0.02 |

| Query | Subject | $d_0$ | C.I. $d_0$ | $d_4$ | C.I. $d_4$ | $d_6$ | C.I. $d_6$ | Diff. G+C Percent |
|---|---|---|---|---|---|---|---|---|
| **'06-16'** | **'06-19'** | 97.2 | [95.6 - 98.3] | 87.6 | [85.1 - 89.8] | 97.5 | [96.3 - 98.4] | 0.05 |
| **'04-15'** | **'05146'** | 98.8 | [97.9 - 99.3] | 87.6 | [85.1 - 89.8] | 98.7 | [97.9 - 99.2] | 0.02 |
| **'06-19'** | **'210931'** | 97.8 | [96.4 - 98.7] | 87.6 | [85.1 - 89.8] | 97.9 | [96.8 - 98.6] | 0.03 |
| **'02-13'** | **'04-15'** | 97.3 | [95.7 - 98.3] | 87.5 | [84.9 - 89.6] | 97.5 | [96.3 - 98.4] | 0.01 |
| **'05146'** | **'BR-AD22'** | 95.5 | [93.4 - 97.0] | 87.5 | [85.0 - 89.7] | 96.3 | [94.7 - 97.4] | 0.1 |
| **'02-13'** | **'06-19'** | 97.2 | [95.6 - 98.3] | 87.5 | [84.9 - 89.7] | 97.5 | [96.2 - 98.3] | 0.01 |
| **'04-3911'** | **'809'** | 97.1 | [95.5 - 98.2] | 87.5 | [84.9 - 89.6] | 97.4 | [96.1 - 98.3] | 0.02 |
| **'0102'** | **'2590'** | 93.8 | [91.2 - 95.7] | 87.5 | [85.0 - 89.7] | 95.0 | [93.1 - 96.4] | 0.06 |
| **'2590'** | **'05146'** | 96.7 | [94.9 - 97.9] | 87.5 | [84.9 - 89.6] | 97.1 | [95.8 - 98.1] | 0.01 |
| **'0211'** | **'2590'** | 93.8 | [91.2 - 95.7] | 87.5 | [85.0 - 89.7] | 95.0 | [93.1 - 96.4] | 0.06 |
| **'06-16'** | **'4940'** | 97.5 | [96.1 - 98.5] | 87.4 | [84.8 - 89.6] | 97.7 | [96.5 - 98.5] | 0.03 |
| **'02-13'** | **'4940'** | 97.5 | [96.0 - 98.5] | 87.4 | [84.8 - 89.6] | 97.7 | [96.5 - 98.5] | 0.0 |
| **'04-3911'** | **'4940'** | 98.1 | [96.8 - 98.9] | 87.4 | [84.9 - 89.6] | 98.1 | [97.1 - 98.8] | 0.02 |
| **'4940'** | **'131001'** | 98.0 | [96.6 - 98.8] | 87.3 | [84.8 - 89.5] | 98.0 | [96.9 - 98.7] | 0.03 |
| **'4940'** | **'210932'** | 98.0 | [96.6 - 98.8] | 87.3 | [84.7 - 89.5] | 98.0 | [96.9 - 98.7] | 0.03 |
| **'2590'** | **'210931'** | 97.0 | [95.3 - 98.1] | 87.3 | [84.8 - 89.5] | 97.3 | [96.0 - 98.2] | 0.01 |
| **'0102'** | **'4940'** | 95.7 | [93.6 - 97.1] | 87.2 | [84.7 - 89.4] | 96.4 | [94.8 - 97.5] | 0.01 |
| **'0211'** | **'4940'** | 95.7 | [93.6 - 97.1] | 87.2 | [84.6 - 89.4] | 96.4 | [94.8 - 97.5] | 0.01 |
| **'06-16'** | **'2590'** | 96.7 | [94.9 - 97.9] | 87.2 | [84.7 - 89.4] | 97.1 | [95.7 - 98.0] | 0.08 |
| **'02-13'** | **'0211'** | 94.8 | [92.5 - 96.5] | 87.1 | [84.5 - 89.3] | 95.7 | [93.9 - 97.0] | 0.02 |
| **'210931'** | **'BR-AD22'** | 96.4 | [94.5 - 97.7] | 87.1 | [84.5 - 89.3] | 96.9 | [95.4 - 97.9] | 0.1 |
| **'02-13'** | **'BR-AD22'** | 94.1 | [91.6 - 95.9] | 87.1 | [84.6 - 89.3] | 95.2 | [93.3 - 96.6] | 0.06 |
| **'02-13'** | **'0102'** | 94.8 | [92.5 - 96.5] | 87.1 | [84.5 - 89.3] | 95.7 | [93.9 - 97.0] | 0.02 |
| **'06-16'** | **'BR-AD22'** | 95.8 | [93.7 - 97.2] | 87.0 | [84.5 - 89.2] | 96.4 | [94.8 - 97.5] | 0.03 |
| **'06-16'** | **'0102'** | 95.1 | [92.9 - 96.7] | 87.0 | [84.4 - 89.2] | 95.9 | [94.2 - 97.1] | 0.02 |
| **'06-16'** | **'0211'** | 95.1 | [92.9 - 96.7] | 87.0 | [84.4 - 89.2] | 95.9 | [94.2 - 97.1] | 0.02 |
| **'0211'** | **'210931'** | 95.7 | [93.6 - 97.1] | 86.9 | [84.3 - 89.1] | 96.3 | [94.7 - 97.5] | 0.06 |

| Query | Subject | $d_0$ | C.I. $d_0$ | $d_4$ | C.I. $d_4$ | $d_6$ | C.I. $d_6$ | Diff. G+C Percent |
|---|---|---|---|---|---|---|---|---|
| **'0102'** | **'210931'** | 95.7 | [93.6 - 97.1] | 86.9 | [84.3 - 89.1] | 96.3 | [94.7 - 97.5] | 0.06 |
| **'0211'** | **'809'** | 97.2 | [95.6 - 98.2] | 86.8 | [84.2 - 89.1] | 97.4 | [96.1 - 98.3] | 0.05 |
| **'4940'** | **'05146'** | 99.1 | [98.4 - 99.5] | 86.8 | [84.2 - 89.0] | 98.8 | [98.1 - 99.3] | 0.04 |
| **'06-19'** | **'131001'** | 97.6 | [96.2 - 98.5] | 86.8 | [84.2 - 89.0] | 97.7 | [96.5 - 98.5] | 0.02 |
| **'06-16'** | **'131001'** | 96.7 | [94.9 - 97.9] | 86.8 | [84.2 - 89.1] | 97.1 | [95.7 - 98.0] | 0.06 |
| **'0102'** | **'809'** | 97.2 | [95.6 - 98.2] | 86.8 | [84.2 - 89.0] | 97.4 | [96.1 - 98.3] | 0.05 |
| **'06-16'** | **'210932'** | 96.7 | [94.9 - 97.9] | 86.8 | [84.2 - 89.0] | 97.1 | [95.7 - 98.0] | 0.06 |
| **'04-15'** | **'131001'** | 97.7 | [96.2 - 98.6] | 86.8 | [84.2 - 89.0] | 97.7 | [96.6 - 98.5] | 0.01 |
| **'04-15'** | **'210932'** | 97.7 | [96.2 - 98.6] | 86.7 | [84.1 - 88.9] | 97.7 | [96.6 - 98.5] | 0.01 |
| **'06-19'** | **'210932'** | 97.7 | [96.2 - 98.6] | 86.7 | [84.1 - 88.9] | 97.7 | [96.6 - 98.5] | 0.01 |
| **'131001'** | **'BR-AD22'** | 96.0 | [94.0 - 97.4] | 86.6 | [84.0 - 88.8] | 96.5 | [95.0 - 97.6] | 0.09 |
| **'210932'** | **'BR-AD22'** | 96.0 | [94.0 - 97.4] | 86.6 | [84.0 - 88.8] | 96.5 | [95.0 - 97.6] | 0.08 |
| **'131001'** | **'210931'** | 97.2 | [95.6 - 98.3] | 86.6 | [84.0 - 88.9] | 97.4 | [96.1 - 98.3] | 0.01 |
| **'210931'** | **'210932'** | 97.2 | [95.6 - 98.2] | 86.6 | [84.0 - 88.9] | 97.4 | [96.1 - 98.3] | 0.01 |
| **'05146'** | **'131001'** | 98.2 | [97.0 - 98.9] | 86.5 | [83.9 - 88.8] | 98.1 | [97.1 - 98.8] | 0.01 |
| **'02-13'** | **'131001'** | 96.8 | [95.0 - 97.9] | 86.5 | [83.9 - 88.7] | 97.1 | [95.7 - 98.0] | 0.03 |
| **'05146'** | **'210932'** | 98.2 | [97.0 - 98.9] | 86.5 | [83.8 - 88.7] | 98.1 | [97.1 - 98.8] | 0.01 |
| **'02-13'** | **'210932'** | 96.8 | [95.0 - 97.9] | 86.4 | [83.8 - 88.7] | 97.1 | [95.7 - 98.0] | 0.02 |
| **'809'** | **'131001'** | 98.5 | [97.4 - 99.1] | 86.3 | [83.6 - 88.5] | 98.3 | [97.3 - 98.9] | 0.01 |
| **'809'** | **'210932'** | 98.5 | [97.4 - 99.1] | 86.2 | [83.6 - 88.5] | 98.3 | [97.4 - 98.9] | 0.01 |
| **'03-8664'** | **'131002'** | 96.4 | [94.4 - 97.6] | 81.1 | [78.2 - 83.7] | 96.2 | [94.5 - 97.3] | 0.25 |
| **'03-8664'** | **'04-7514'** | 96.2 | [94.2 - 97.5] | 78.8 | [75.9 - 81.5] | 95.7 | [93.9 - 97.0] | 0.16 |
| **'04-7514'** | **'131002'** | 96.4 | [94.5 - 97.7] | 75.9 | [72.9 - 78.6] | 95.5 | [93.7 - 96.8] | 0.09 |
| **'03-8664'** | **'0102'** | 95.0 | [92.6 - 96.6] | 68.2 | [65.2 - 71.0] | 93.1 | [90.7 - 94.8] | 0.27 |
| **'03-8664'** | **'04-3911'** | 97.2 | [95.6 - 98.3] | 68.1 | [65.1 - 71.0] | 95.0 | [93.1 - 96.4] | 0.3 |
| **'03-8664'** | **'05146'** | 96.7 | [94.9 - 97.9] | 68.1 | [65.1 - 71.0] | 94.6 | [92.5 - 96.1] | 0.32 |
| **'03-8664'** | **'0211'** | 95.0 | [92.7 - 96.6] | 68.1 | [65.2 - 71.0] | 93.1 | [90.7 - 94.8] | 0.27 |

| Query | Subject | $d_0$ | C.I. $d_0$ | $d_4$ | C.I. $d_4$ | $d_6$ | C.I. $d_6$ | Diff. G+C Percent |
|---|---|---|---|---|---|---|---|---|
| **'03-8664'** | **'210932'** | 95.6 | [93.4 - 97.1] | 68.0 | [65.0 - 70.9] | 93.6 | [91.3 - 95.2] | 0.31 |
| **'03-8664'** | **'131001'** | 95.6 | [93.4 - 97.1] | 68.0 | [65.0 - 70.9] | 93.6 | [91.4 - 95.2] | 0.31 |
| **'03-8664'** | *Corynebacterium ulcerans* NCTC 7910 | 96.2 | [94.3 - 97.5] | 67.8 | [64.8 - 70.6] | 94.1 | [92.0 - 95.7] | 0.31 |
| **'03-8664'** | **'809'** | 95.3 | [93.0 - 96.8] | 67.7 | [64.7 - 70.5] | 93.2 | [90.9 - 95.0] | 0.32 |
| **'03-8664'** | **'06-19'** | 96.3 | [94.3 - 97.6] | 67.7 | [64.7 - 70.5] | 94.1 | [92.0 - 95.7] | 0.3 |
| **'03-8664'** | **'2590'** | 94.7 | [92.3 - 96.4] | 67.7 | [64.7 - 70.5] | 92.7 | [90.4 - 94.6] | 0.33 |
| **'03-8664'** | **'BR-AD22'** | 92.7 | [89.9 - 94.8] | 67.7 | [64.7 - 70.5] | 91.1 | [88.5 - 93.1] | 0.23 |
| **'03-8664'** | **'04-15'** | 96.3 | [94.3 - 97.6] | 67.7 | [64.7 - 70.5] | 94.1 | [92.0 - 95.7] | 0.3 |
| **'03-8664'** | **'210931'** | 95.9 | [93.9 - 97.3] | 67.6 | [64.6 - 70.4] | 93.8 | [91.6 - 95.4] | 0.32 |
| **'02-13'** | **'03-8664'** | 94.9 | [92.5 - 96.5] | 67.6 | [64.6 - 70.4] | 92.9 | [90.5 - 94.7] | 0.29 |
| **'03-8664'** | **'4940'** | 96.7 | [94.9 - 97.9] | 67.5 | [64.5 - 70.4] | 94.4 | [92.4 - 96.0] | 0.28 |
| **'03-8664'** | **'06-16'** | 95.3 | [93.1 - 96.8] | 67.4 | [64.5 - 70.3] | 93.2 | [90.9 - 94.9] | 0.25 |
| **'04-7514'** | **'0102'** | 97.8 | [96.4 - 98.7] | 63.7 | [60.8 - 66.5] | 94.8 | [92.9 - 96.3] | 0.11 |
| **'04-7514'** | **'0211'** | 97.8 | [96.4 - 98.7] | 63.7 | [60.8 - 66.5] | 94.8 | [92.9 - 96.3] | 0.11 |
| **'05146'** | **'131002'** | 98.4 | [97.3 - 99.1] | 63.7 | [60.8 - 66.5] | 95.5 | [93.7 - 96.8] | 0.07 |
| **'131001'** | **'131002'** | 97.6 | [96.2 - 98.5] | 63.5 | [60.6 - 66.4] | 94.6 | [92.6 - 96.1] | 0.07 |
| **'131002'** | **'210932'** | 97.6 | [96.1 - 98.5] | 63.5 | [60.6 - 66.4] | 94.6 | [92.6 - 96.1] | 0.06 |
| **'131002'** | *Corynebacterium ulcerans* NCTC 7910 | 98.3 | [97.1 - 99.0] | 63.4 | [60.5 - 66.2] | 95.2 | [93.4 - 96.6] | 0.06 |
| **'131002'** | **'210931'** | 97.3 | [95.8 - 98.3] | 63.3 | [60.4 - 66.1] | 94.3 | [92.2 - 95.8] | 0.08 |
| **'04-3911'** | **'131002'** | 96.8 | [95.0 - 98.0] | 63.3 | [60.4 - 66.1] | 93.8 | [91.6 - 95.4] | 0.06 |
| **'04-7514'** | *Corynebacterium ulcerans* NCTC 7910 | 96.2 | [94.2 - 97.5] | 63.3 | [60.4 - 66.2] | 93.2 | [90.9 - 94.9] | 0.16 |
| **'04-7514'** | **'131001'** | 95.3 | [93.0 - 96.8] | 63.3 | [60.4 - 66.1] | 92.4 | [89.9 - 94.2] | 0.16 |
| **'04-3911'** | **'04-7514'** | 97.7 | [96.2 - 98.6] | 63.3 | [60.4 - 66.1] | 94.6 | [92.6 - 96.1] | 0.15 |
| **'04-7514'** | **'210932'** | 95.3 | [93.1 - 96.8] | 63.3 | [60.4 - 66.1] | 92.4 | [89.9 - 94.2] | 0.15 |
| **'0102'** | **'131002'** | 94.6 | [92.1 - 96.3] | 63.2 | [60.3 - 66.0] | 91.7 | [89.2 - 93.7] | 0.02 |
| **'2590'** | **'131002'** | 96.8 | [95.0 - 97.9] | 63.2 | [60.3 - 66.1] | 93.7 | [91.5 - 95.4] | 0.08 |
| **'0211'** | **'131002'** | 94.6 | [92.2 - 96.3] | 63.2 | [60.3 - 66.0] | 91.7 | [89.2 - 93.7] | 0.02 |

| Query | Subject | $d_0$ | C.I. $d_0$ | $d_4$ | C.I. $d_4$ | $d_6$ | C.I. $d_6$ | Diff. G+C Percent |
|---|---|---|---|---|---|---|---|---|
| **'04-7514'** | **'05146'** | 96.8 | [95.0 - 98.0] | 63.2 | [60.3 - 66.0] | 93.7 | [91.6 - 95.4] | 0.16 |
| **'06-16'** | **'131002'** | 96.9 | [95.2 - 98.0] | 63.1 | [60.2 - 65.9] | 93.8 | [91.7 - 95.5] | 0.0 |
| **'809'** | **'131002'** | 97.1 | [95.5 - 98.2] | 63.1 | [60.2 - 65.9] | 94.0 | [91.9 - 95.6] | 0.07 |
| **'02-13'** | **'131002'** | 96.9 | [95.1 - 98.0] | 63.1 | [60.2 - 65.9] | 93.8 | [91.6 - 95.4] | 0.04 |
| **'04-7514'** | **'809'** | 95.7 | [93.6 - 97.1] | 63.0 | [60.1 - 65.8] | 92.7 | [90.3 - 94.5] | 0.17 |
| **'131002'** | **'BR-AD22'** | 94.0 | [91.4 - 95.8] | 63.0 | [60.1 - 65.8] | 91.2 | [88.6 - 93.2] | 0.02 |
| **'04-7514'** | **'2590'** | 94.7 | [92.4 - 96.4] | 62.9 | [60.0 - 65.7] | 91.8 | [89.3 - 93.8] | 0.17 |
| **'4940'** | **'131002'** | 98.5 | [97.5 - 99.1] | 62.9 | [60.0 - 65.7] | 95.4 | [93.6 - 96.7] | 0.03 |
| **'04-7514'** | **'4940'** | 97.3 | [95.8 - 98.3] | 62.8 | [59.9 - 65.6] | 94.2 | [92.1 - 95.8] | 0.12 |
| **'06-19'** | **'131002'** | 97.8 | [96.4 - 98.7] | 62.8 | [59.9 - 65.6] | 94.6 | [92.6 - 96.1] | 0.05 |
| **'04-15'** | **'131002'** | 97.8 | [96.4 - 98.7] | 62.8 | [59.9 - 65.6] | 94.7 | [92.7 - 96.2] | 0.05 |
| **'04-7514'** | **'210931'** | 96.5 | [94.7 - 97.8] | 62.7 | [59.8 - 65.5] | 93.4 | [91.1 - 95.1] | 0.17 |
| **'04-7514'** | **'BR-AD22'** | 95.4 | [93.2 - 96.9] | 62.6 | [59.7 - 65.4] | 92.4 | [89.9 - 94.2] | 0.07 |
| **'04-15'** | **'04-7514'** | 96.1 | [94.2 - 97.5] | 62.6 | [59.7 - 65.4] | 93.0 | [90.7 - 94.8] | 0.14 |
| **'04-7514'** | **'06-19'** | 96.1 | [94.1 - 97.4] | 62.6 | [59.7 - 65.4] | 93.0 | [90.6 - 94.7] | 0.14 |
| **'02-13'** | **'04-7514'** | 95.3 | [93.0 - 96.8] | 62.5 | [59.7 - 65.3] | 92.2 | [89.7 - 94.1] | 0.13 |
| **'04-7514'** | **'06-16'** | 95.7 | [93.6 - 97.2] | 62.4 | [59.5 - 65.2] | 92.6 | [90.2 - 94.4] | 0.09 |
| **'03-8664'** | **'04-13'** | 90.2 | [87.0 - 92.7] | 42.7 | [40.2 - 45.3] | 81.1 | [77.7 - 84.0] | 0.81 |
| **'03-8664'** | **'05-13'** | 90.1 | [86.9 - 92.6] | 42.7 | [40.2 - 45.3] | 81.0 | [77.6 - 83.9] | 0.79 |
| **'03-8664'** | *Corynebacterium silvaticum* KL0182 | 90.0 | [86.8 - 92.5] | 42.7 | [40.2 - 45.2] | 80.9 | [77.5 - 83.9] | 0.82 |
| **'05-13'** | **'131002'** | 92.4 | [89.5 - 94.5] | 41.6 | [39.1 - 44.2] | 82.4 | [79.1 - 85.3] | 1.04 |
| **'131002'** | *Corynebacterium silvaticum* KL0182 | 92.3 | [89.4 - 94.4] | 41.6 | [39.1 - 44.1] | 82.3 | [79.0 - 85.2] | 1.07 |
| **'04-13'** | **'131002'** | 92.5 | [89.6 - 94.6] | 41.6 | [39.1 - 44.1] | 82.5 | [79.2 - 85.4] | 1.06 |
| **'04-7514'** | **'05-13'** | 91.1 | [88.0 - 93.5] | 41.2 | [38.7 - 43.7] | 81.1 | [77.7 - 84.1] | 0.95 |
| **'04-13'** | **'04-7514'** | 91.2 | [88.1 - 93.6] | 41.2 | [38.7 - 43.7] | 81.2 | [77.8 - 84.1] | 0.97 |
| **'05-13'** | **'06-19'** | 91.6 | [88.6 - 93.9] | 41.2 | [38.7 - 43.7] | 81.5 | [78.2 - 84.5] | 1.09 |
| **'04-7514'** | *Corynebacterium silvaticum* KL0182 | 91.0 | [87.9 - 93.4] | 41.2 | [38.7 - 43.7] | 81.0 | [77.6 - 84.0] | 0.98 |

| Query | Subject | $d_0$ | C.I. $d_0$ | $d_4$ | C.I. $d_4$ | $d_6$ | C.I. $d_6$ | Diff. G+C Percent |
|---|---|---|---|---|---|---|---|---|
| **'06-19'** | *Corynebacterium silvaticum* KL0182 | 91.5 | [88.5 - 93.8] | 41.1 | [38.6 - 43.7] | 81.4 | [78.1 - 84.4] | 1.12 |
| **'04-13'** | **'04-15'** | 91.8 | [88.9 - 94.1] | 41.1 | [38.6 - 43.6] | 81.7 | [78.3 - 84.6] | 1.11 |
| **'04-15'** | **'05-13'** | 91.7 | [88.7 - 94.0] | 41.1 | [38.6 - 43.7] | 81.6 | [78.3 - 84.6] | 1.09 |
| **'04-13'** | **'06-19'** | 91.7 | [88.7 - 94.0] | 41.1 | [38.6 - 43.7] | 81.6 | [78.2 - 84.5] | 1.11 |
| **'04-15'** | *Corynebacterium silvaticum* KL0182 | 91.6 | [88.6 - 93.9] | 41.1 | [38.6 - 43.7] | 81.5 | [78.2 - 84.5] | 1.12 |
| **'4940'** | *Corynebacterium silvaticum* KL0182 | 92.6 | [89.7 - 94.7] | 41.0 | [38.5 - 43.5] | 82.3 | [79.0 - 85.2] | 1.1 |
| **'05-13'** | *Corynebacterium ulcerans* NCTC 7910 | 91.8 | [88.8 - 94.0] | 41.0 | [38.5 - 43.5] | 81.6 | [78.2 - 84.5] | 1.11 |
| **'04-3911'** | *Corynebacterium silvaticum* KL0182 | 91.9 | [89.0 - 94.2] | 41.0 | [38.5 - 43.5] | 81.7 | [78.4 - 84.7] | 1.13 |
| **'05-13'** | **'210932'** | 91.5 | [88.5 - 93.8] | 41.0 | [38.5 - 43.5] | 81.4 | [78.0 - 84.3] | 1.1 |
| **'210932'** | *Corynebacterium silvaticum* KL0182 | 91.4 | [88.3 - 93.7] | 41.0 | [38.5 - 43.5] | 81.2 | [77.9 - 84.2] | 1.13 |
| **'05-13'** | **'131001'** | 91.5 | [88.5 - 93.8] | 41.0 | [38.5 - 43.5] | 81.4 | [78.0 - 84.3] | 1.11 |
| **'04-13'** | **'4940'** | 92.8 | [90.0 - 94.8] | 41.0 | [38.5 - 43.5] | 82.5 | [79.1 - 85.4] | 1.09 |
| **'04-3911'** | **'05-13'** | 92.0 | [89.1 - 94.2] | 41.0 | [38.5 - 43.5] | 81.8 | [78.4 - 84.7] | 1.1 |
| **'05-13'** | **'0102'** | 89.2 | [85.8 - 91.8] | 41.0 | [38.5 - 43.5] | 79.3 | [75.9 - 82.3] | 1.06 |
| **'131001'** | *Corynebacterium silvaticum* KL0182 | 91.4 | [88.4 - 93.7] | 41.0 | [38.5 - 43.5] | 81.3 | [77.9 - 84.2] | 1.14 |
| **'809'** | *Corynebacterium silvaticum* KL0182 | 90.7 | [87.6 - 93.1] | 41.0 | [38.5 - 43.6] | 80.7 | [77.3 - 83.7] | 1.14 |
| **'04-13'** | **'210932'** | 91.6 | [88.6 - 93.9] | 41.0 | [38.5 - 43.5] | 81.4 | [78.1 - 84.4] | 1.12 |
| **'05-13'** | **'809'** | 90.8 | [87.7 - 93.2] | 41.0 | [38.5 - 43.6] | 80.8 | [77.4 - 83.7] | 1.11 |
| **'05-13'** | **'4940'** | 92.7 | [89.8 - 94.7] | 41.0 | [38.5 - 43.5] | 82.4 | [79.1 - 85.3] | 1.07 |
| **'05-13'** | **'0211'** | 89.2 | [85.8 - 91.8] | 41.0 | [38.5 - 43.5] | 79.3 | [75.9 - 82.4] | 1.06 |
| **'04-13'** | **'04-3911'** | 92.1 | [89.2 - 94.3] | 41.0 | [38.5 - 43.5] | 81.9 | [78.5 - 84.8] | 1.11 |
| **'04-13'** | **'809'** | 90.9 | [87.8 - 93.3] | 41.0 | [38.5 - 43.5] | 80.8 | [77.5 - 83.8] | 1.13 |
| **'04-13'** | **'131001'** | 91.6 | [88.6 - 93.9] | 41.0 | [38.5 - 43.5] | 81.4 | [78.1 - 84.4] | 1.12 |
| **'02-13'** | **'04-13'** | 90.2 | [86.9 - 92.7] | 40.9 | [38.4 - 43.5] | 80.1 | [76.7 - 83.1] | 1.1 |
| **'04-13'** | **'BR-AD22'** | 88.1 | [84.6 - 90.9] | 40.9 | [38.4 - 43.5] | 78.4 | [74.9 - 81.4] | 1.04 |
| **'04-13'** | **'0211'** | 89.3 | [86.0 - 91.9] | 40.9 | [38.4 - 43.5] | 79.4 | [76.0 - 82.4] | 1.08 |
| **'02-13'** | *Corynebacterium silvaticum* KL0182 | 90.0 | [86.7 - 92.5] | 40.9 | [38.4 - 43.5] | 80.0 | [76.5 - 83.0] | 1.11 |

| Query | Subject | $d_0$ | C.I. $d_0$ | $d_4$ | C.I. $d_4$ | $d_6$ | C.I. $d_6$ | Diff. G+C Percent |
|---|---|---|---|---|---|---|---|---|
| **'0102'** | *Corynebacterium silvaticum* KL0182 | 89.1 | [85.7 - 91.7] | 40.9 | [38.4 - 43.5] | 79.2 | [75.8 - 82.3] | 1.09 |
| **'04-13'** | *Corynebacterium ulcerans* NCTC 7910 | 91.9 | [88.9 - 94.1] | 40.9 | [38.4 - 43.5] | 81.7 | [78.3 - 84.6] | 1.12 |
| **'0211'** | *Corynebacterium silvaticum* KL0182 | 89.1 | [85.8 - 91.7] | 40.9 | [38.4 - 43.5] | 79.2 | [75.8 - 82.3] | 1.09 |
| **'04-13'** | **'0102'** | 89.3 | [85.9 - 91.9] | 40.9 | [38.4 - 43.5] | 79.4 | [76.0 - 82.4] | 1.08 |
| **'04-13'** | **'05146'** | 92.2 | [89.3 - 94.4] | 40.9 | [38.4 - 43.5] | 81.9 | [78.6 - 84.9] | 1.13 |
| **'05-13'** | **'05146'** | 92.2 | [89.3 - 94.4] | 40.9 | [38.4 - 43.5] | 81.9 | [78.6 - 84.8] | 1.11 |
| **'05146'** | *Corynebacterium silvaticum* KL0182 | 92.0 | [89.1 - 94.2] | 40.9 | [38.4 - 43.5] | 81.7 | [78.4 - 84.7] | 1.14 |
| **'02-13'** | **'05-13'** | 90.2 | [87.0 - 92.7] | 40.9 | [38.4 - 43.4] | 80.1 | [76.7 - 83.2] | 1.08 |
| **'BR-AD22'** | *Corynebacterium silvaticum* KL0182 | 87.9 | [84.4 - 90.7] | 40.9 | [38.4 - 43.5] | 78.2 | [74.7 - 81.3] | 1.05 |
| **'05-13'** | **'BR-AD22'** | 88.1 | [84.7 - 90.9] | 40.9 | [38.4 - 43.4] | 78.4 | [74.9 - 81.5] | 1.02 |
| **'04-13'** | **'210931'** | 91.0 | [87.9 - 93.4] | 40.8 | [38.3 - 43.3] | 80.8 | [77.4 - 83.8] | 1.13 |
| **'210931'** | *Corynebacterium silvaticum* KL0182 | 90.8 | [87.7 - 93.2] | 40.8 | [38.3 - 43.3] | 80.6 | [77.2 - 83.6] | 1.15 |
| **'04-13'** | **'2590'** | 90.7 | [87.5 - 93.1] | 40.8 | [38.3 - 43.3] | 80.5 | [77.1 - 83.5] | 1.14 |
| **'2590'** | *Corynebacterium silvaticum* KL0182 | 90.5 | [87.3 - 92.9] | 40.8 | [38.3 - 43.3] | 80.3 | [76.9 - 83.3] | 1.15 |
| **'05-13'** | **'2590'** | 90.7 | [87.6 - 93.1] | 40.7 | [38.2 - 43.3] | 80.5 | [77.1 - 83.5] | 1.12 |
| **'06-16'** | *Corynebacterium silvaticum* KL0182 | 90.4 | [87.2 - 92.8] | 40.7 | [38.2 - 43.2] | 80.2 | [76.8 - 83.2] | 1.07 |
| **'05-13'** | **'210931'** | 91.2 | [88.1 - 93.5] | 40.7 | [38.2 - 43.2] | 80.9 | [77.5 - 83.9] | 1.12 |
| **'04-13'** | **'06-16'** | 90.6 | [87.4 - 93.0] | 40.7 | [38.2 - 43.2] | 80.4 | [77.0 - 83.4] | 1.06 |
| **'05-13'** | **'06-16'** | 90.7 | [87.6 - 93.1] | 40.6 | [38.2 - 43.2] | 80.5 | [77.1 - 83.5] | 1.04 |
| **'03-8664'** | *Corynebacterium pseudotuberculosis* ATCC 19410 | 76.4 | [72.4 - 80.0] | 28.6 | [26.2 - 31.1] | 61.5 | [58.2 - 64.6] | 1.44 |
| **'03-8664'** | *Corynebacterium pseudotuberculosis* DSM 20689 | 76.4 | [72.4 - 80.0] | 28.6 | [26.2 - 31.1] | 61.5 | [58.2 - 64.7] | 1.44 |
| **'05-13'** | *Corynebacterium pseudotuberculosis* ATCC 19410 | 82.9 | [79.0 - 86.1] | 28.6 | [26.2 - 31.1] | 65.9 | [62.5 - 69.1] | 2.24 |
| **'05-13'** | *Corynebacterium pseudotuberculosis* DSM 20689 | 83.0 | [79.1 - 86.2] | 28.6 | [26.2 - 31.1] | 66.0 | [62.6 - 69.2] | 2.23 |
| **'04-13'** | *Corynebacterium pseudotuberculosis* DSM 20689 | 83.1 | [79.3 - 86.3] | 28.5 | [26.2 - 31.0] | 66.1 | [62.7 - 69.3] | 2.25 |
| **'04-13'** | *Corynebacterium pseudotuberculosis* ATCC 19410 | 83.0 | [79.2 - 86.2] | 28.5 | [26.2 - 31.0] | 66.0 | [62.6 - 69.2] | 2.25 |

| Query | Subject | $d_0$ | C.I. $d_0$ | $d_4$ | C.I. $d_4$ | $d_6$ | C.I. $d_6$ | Diff. G+C Percent |
|---|---|---|---|---|---|---|---|---|
| **'0102'** | *Corynebacterium pseudotuberculosis* ATCC 19410 | 83.2 | [79.3 - 86.4] | 27.8 | [25.4 - 30.3] | 65.5 | [62.1 - 68.7] | 1.18 |
| **'0211'** | *Corynebacterium pseudotuberculosis* DSM 20689 | 83.2 | [79.4 - 86.4] | 27.8 | [25.4 - 30.3] | 65.5 | [62.1 - 68.7] | 1.17 |
| **'0102'** | *Corynebacterium pseudotuberculosis* DSM 20689 | 83.2 | [79.4 - 86.4] | 27.8 | [25.4 - 30.3] | 65.5 | [62.1 - 68.7] | 1.17 |
| **'04-3911'** | *Corynebacterium pseudotuberculosis* ATCC 19410 | 85.6 | [81.9 - 88.6] | 27.8 | [25.4 - 30.3] | 67.2 | [63.8 - 70.4] | 1.14 |
| **'BR-AD22'** | *Corynebacterium pseudotuberculosis* ATCC 19410 | 82.9 | [79.1 - 86.2] | 27.8 | [25.4 - 30.3] | 65.2 | [61.9 - 68.5] | 1.22 |
| **'BR-AD22'** | *Corynebacterium pseudotuberculosis* DSM 20689 | 82.9 | [79.1 - 86.2] | 27.8 | [25.4 - 30.3] | 65.3 | [61.9 - 68.5] | 1.21 |
| **'04-3911'** | *Corynebacterium pseudotuberculosis* DSM 20689 | 85.6 | [81.9 - 88.6] | 27.8 | [25.4 - 30.3] | 67.2 | [63.8 - 70.4] | 1.14 |
| **'0211'** | *Corynebacterium pseudotuberculosis* ATCC 19410 | 83.2 | [79.4 - 86.4] | 27.8 | [25.4 - 30.3] | 65.5 | [62.1 - 68.7] | 1.17 |
| **'210931'** | *Corynebacterium pseudotuberculosis* ATCC 19410 | 86.0 | [82.3 - 89.0] | 27.7 | [25.4 - 30.2] | 67.4 | [64.0 - 70.7] | 1.12 |
| **'05146'** | *Corynebacterium pseudotuberculosis* ATCC 19410 | 87.4 | [83.8 - 90.2] | 27.7 | [25.4 - 30.2] | 68.4 | [65.0 - 71.7] | 1.12 |
| **'131001'** | *Corynebacterium pseudotuberculosis* DSM 20689 | 86.7 | [83.1 - 89.6] | 27.7 | [25.4 - 30.2] | 67.9 | [64.5 - 71.2] | 1.13 |
| **'210931'** | *Corynebacterium pseudotuberculosis* DSM 20689 | 86.0 | [82.4 - 89.0] | 27.7 | [25.4 - 30.2] | 67.4 | [64.0 - 70.7] | 1.12 |
| **'809'** | *Corynebacterium pseudotuberculosis* DSM 20689 | 86.1 | [82.5 - 89.1] | 27.7 | [25.4 - 30.2] | 67.5 | [64.1 - 70.8] | 1.12 |
| **'05146'** | *Corynebacterium pseudotuberculosis* DSM 20689 | 87.4 | [83.8 - 90.2] | 27.7 | [25.4 - 30.2] | 68.5 | [65.0 - 71.7] | 1.12 |
| **'131002'** | *Corynebacterium pseudotuberculosis* DSM 20689 | 88.6 | [85.2 - 91.3] | 27.7 | [25.4 - 30.2] | 69.4 | [66.0 - 72.7] | 1.19 |
| **'131002'** | *Corynebacterium pseudotuberculosis* ATCC 19410 | 88.6 | [85.2 - 91.3] | 27.7 | [25.4 - 30.2] | 69.4 | [66.0 - 72.6] | 1.19 |
| **'210932'** | *Corynebacterium pseudotuberculosis* DSM 20689 | 86.7 | [83.1 - 89.6] | 27.7 | [25.4 - 30.2] | 67.9 | [64.5 - 71.2] | 1.13 |
| **'809'** | *Corynebacterium pseudotuberculosis* ATCC 19410 | 86.1 | [82.5 - 89.1] | 27.7 | [25.4 - 30.2] | 67.5 | [64.1 - 70.7] | 1.12 |
| **'210932'** | *Corynebacterium pseudotuberculosis* ATCC 19410 | 86.7 | [83.1 - 89.6] | 27.7 | [25.4 - 30.2] | 67.9 | [64.5 - 71.2] | 1.13 |
| **'131001'** | *Corynebacterium pseudotuberculosis* ATCC 19410 | 86.7 | [83.1 - 89.6] | 27.7 | [25.4 - 30.2] | 67.9 | [64.5 - 71.2] | 1.13 |
| **'04-7514'** | *Corynebacterium pseudotuberculosis* DSM 20689 | 85.5 | [81.8 - 88.6] | 27.6 | [25.3 - 30.1] | 67.0 | [63.6 - 70.2] | 1.28 |

| Query | Subject | $d_0$ | C.I. $d_0$ | $d_4$ | C.I. $d_4$ | $d_6$ | C.I. $d_6$ | Diff. G+C Percent |
|---|---|---|---|---|---|---|---|---|
| **'4940'** | *Corynebacterium pseudotuberculosis* ATCC 19410 | 87.6 | [84.1 - 90.5] | 27.6 | [25.2 - 30.0] | 68.5 | [65.0 - 71.7] | 1.16 |
| **'04-15'** | *Corynebacterium pseudotuberculosis* DSM 20689 | 87.9 | [84.4 - 90.7] | 27.6 | [25.2 - 30.1] | 68.7 | [65.3 - 71.9] | 1.14 |
| **'4940'** | *Corynebacterium pseudotuberculosis* DSM 20689 | 87.7 | [84.1 - 90.5] | 27.6 | [25.2 - 30.0] | 68.5 | [65.1 - 71.7] | 1.16 |
| **'02-13'** | *Corynebacterium pseudotuberculosis* ATCC 19410 | 85.0 | [81.3 - 88.1] | 27.6 | [25.2 - 30.0] | 66.5 | [63.2 - 69.8] | 1.16 |
| **'06-19'** | *Corynebacterium pseudotuberculosis* ATCC 19410 | 87.6 | [84.0 - 90.4] | 27.6 | [25.3 - 30.1] | 68.5 | [65.0 - 71.7] | 1.15 |
| **'02-13'** | *Corynebacterium pseudotuberculosis* DSM 20689 | 85.0 | [81.3 - 88.1] | 27.6 | [25.2 - 30.0] | 66.6 | [63.2 - 69.8] | 1.15 |
| **'04-7514'** | *Corynebacterium pseudotuberculosis* ATCC 19410 | 85.5 | [81.8 - 88.5] | 27.6 | [25.3 - 30.1] | 66.9 | [63.5 - 70.2] | 1.29 |
| **'06-19'** | *Corynebacterium pseudotuberculosis* DSM 20689 | 87.7 | [84.2 - 90.5] | 27.6 | [25.2 - 30.1] | 68.6 | [65.1 - 71.8] | 1.14 |
| **'04-15'** | *Corynebacterium pseudotuberculosis* ATCC 19410 | 87.9 | [84.4 - 90.7] | 27.6 | [25.2 - 30.1] | 68.7 | [65.3 - 71.9] | 1.14 |
| **'06-16'** | *Corynebacterium pseudotuberculosis* DSM 20689 | 85.3 | [81.5 - 88.3] | 27.5 | [25.2 - 30.0] | 66.7 | [63.3 - 69.9] | 1.19 |
| **'2590'** | *Corynebacterium pseudotuberculosis* DSM 20689 | 85.8 | [82.1 - 88.8] | 27.5 | [25.2 - 30.0] | 67.1 | [63.7 - 70.3] | 1.11 |
| **'2590'** | *Corynebacterium pseudotuberculosis* ATCC 19410 | 85.6 | [82.0 - 88.7] | 27.5 | [25.2 - 30.0] | 67.0 | [63.6 - 70.2] | 1.11 |
| **'06-16'** | *Corynebacterium pseudotuberculosis* ATCC 19410 | 85.2 | [81.5 - 88.3] | 27.5 | [25.2 - 30.0] | 66.7 | [63.3 - 69.9] | 1.19 |
| **'03-8664'** | *Corynebacterium mustelae* DSM 45274 | 13.0 | [10.3 - 16.3] | 26.7 | [24.3 - 29.2] | 13.4 | [11.0 - 16.1] | 1.05 |
| **'05-13'** | *Corynebacterium resistens* DSM 45100 | 13.1 | [10.4 - 16.4] | 26.4 | [24.1 - 28.9] | 13.5 | [11.1 - 16.2] | 2.67 |
| **'131002'** | *Corynebacterium mustelae* DSM 45274 | 13.0 | [10.3 - 16.3] | 26.2 | [23.8 - 28.7] | 13.4 | [11.0 - 16.2] | 0.81 |
| **'04-7514'** | *Corynebacterium mustelae* DSM 45274 | 13.0 | [10.3 - 16.3] | 26.2 | [23.8 - 28.7] | 13.4 | [11.0 - 16.1] | 0.9 |
| **'210931'** | *Corynebacterium mustelae* DSM 45274 | 13.0 | [10.3 - 16.3] | 26.1 | [23.8 - 28.6] | 13.4 | [11.0 - 16.2] | 0.73 |
| **'05146'** | *Corynebacterium mustelae* DSM 45274 | 13.0 | [10.3 - 16.3] | 26.1 | [23.8 - 28.6] | 13.4 | [11.0 - 16.2] | 0.73 |
| **'210932'** | *Corynebacterium mustelae* DSM 45274 | 13.0 | [10.3 - 16.3] | 26.1 | [23.7 - 28.6] | 13.4 | [11.0 - 16.2] | 0.74 |
| **'04-3911'** | *Corynebacterium mustelae* DSM 45274 | 13.0 | [10.3 - 16.3] | 26.1 | [23.8 - 28.6] | 13.4 | [11.0 - 16.2] | 0.75 |
| **'809'** | *Corynebacterium mustelae* DSM 45274 | 13.0 | [10.3 - 16.3] | 26.1 | [23.7 - 28.6] | 13.4 | [11.0 - 16.2] | 0.73 |
| **'BR-AD22'** | *Corynebacterium mustelae* DSM 45274 | 13.0 | [10.3 - 16.3] | 26.1 | [23.8 - 28.6] | 13.4 | [11.0 - 16.1] | 0.83 |

| Query | Subject | $d_0$ | C.I. $d_0$ | $d_4$ | C.I. $d_4$ | $d_6$ | C.I. $d_6$ | Diff. G+C Percent |
|---|---|---|---|---|---|---|---|---|
| **'0211'** | *Corynebacterium mustelae* DSM 45274 | 13.0 | [10.3 - 16.3] | 26.1 | [23.8 - 28.6] | 13.4 | [11.0 - 16.2] | 0.79 |
| **'131001'** | *Corynebacterium mustelae* DSM 45274 | 13.0 | [10.3 - 16.3] | 26.1 | [23.8 - 28.6] | 13.4 | [11.0 - 16.2] | 0.74 |
| **'0102'** | *Corynebacterium mustelae* DSM 45274 | 13.0 | [10.3 - 16.3] | 26.1 | [23.8 - 28.6] | 13.4 | [11.0 - 16.2] | 0.79 |
| **'04-7514'** | *Corynebacterium vitaeruminis* DSM 20294 | 13.2 | [10.5 - 16.6] | 26.0 | [23.7 - 28.5] | 13.6 | [11.2 - 16.4] | 12.06 |
| **'BR-AD22'** | *Corynebacterium vitaeruminis* DSM 20294 | 13.2 | [10.5 - 16.5] | 26.0 | [23.6 - 28.4] | 13.6 | [11.2 - 16.4] | 12.13 |
| **'05146'** | *Corynebacterium vitaeruminis* DSM 20294 | 13.3 | [10.5 - 16.6] | 25.8 | [23.4 - 28.3] | 13.6 | [11.3 - 16.4] | 12.22 |
| **'809'** | *Corynebacterium vitaeruminis* DSM 20294 | 13.2 | [10.5 - 16.6] | 25.8 | [23.4 - 28.2] | 13.6 | [11.2 - 16.4] | 12.22 |
| **'04-3911'** | *Corynebacterium vitaeruminis* DSM 20294 | 13.3 | [10.5 - 16.6] | 25.8 | [23.5 - 28.3] | 13.6 | [11.3 - 16.4] | 12.2 |
| **'0102'** | *Corynebacterium pseudopelargi* CCM 8832 | 13.3 | [10.5 - 16.6] | 25.7 | [23.4 - 28.2] | 13.6 | [11.3 - 16.4] | 4.55 |
| **'0211'** | *Corynebacterium pseudopelargi* CCM 8832 | 13.3 | [10.5 - 16.6] | 25.7 | [23.4 - 28.2] | 13.6 | [11.3 - 16.4] | 4.55 |
| **'04-3911'** | *Corynebacterium pseudopelargi* CCM 8832 | 13.3 | [10.5 - 16.6] | 25.7 | [23.4 - 28.2] | 13.6 | [11.3 - 16.4] | 4.59 |
| **'131002'** | *Corynebacterium vitaeruminis* DSM 20294 | 13.3 | [10.5 - 16.6] | 25.7 | [23.4 - 28.2] | 13.6 | [11.3 - 16.4] | 12.15 |
| **'0211'** | *Corynebacterium vitaeruminis* DSM 20294 | 13.3 | [10.5 - 16.6] | 25.6 | [23.2 - 28.0] | 13.6 | [11.3 - 16.4] | 12.17 |
| **'0102'** | *Corynebacterium vitaeruminis* DSM 20294 | 13.3 | [10.5 - 16.6] | 25.6 | [23.2 - 28.1] | 13.6 | [11.3 - 16.4] | 12.17 |
| **'210931'** | *Corynebacterium vitaeruminis* DSM 20294 | 13.3 | [10.5 - 16.6] | 25.6 | [23.3 - 28.1] | 13.6 | [11.3 - 16.4] | 12.22 |
| **'210932'** | *Corynebacterium vitaeruminis* DSM 20294 | 13.3 | [10.5 - 16.6] | 25.5 | [23.2 - 28.0] | 13.6 | [11.3 - 16.4] | 12.21 |
| **'BR-AD22'** | *Corynebacterium pelargi* DSM 46737 | 13.2 | [10.5 - 16.6] | 25.5 | [23.2 - 28.0] | 13.6 | [11.2 - 16.4] | 4.77 |
| **'131001'** | *Corynebacterium vitaeruminis* DSM 20294 | 13.3 | [10.5 - 16.6] | 25.5 | [23.2 - 28.0] | 13.6 | [11.3 - 16.4] | 12.21 |
| **'05146'** | *Corynebacterium resistens* DSM 45100 | 12.9 | [10.2 - 16.2] | 25.5 | [23.1 - 27.9] | 13.3 | [10.9 - 16.1] | 3.79 |
| **'0102'** | *Corynebacterium pelargi* DSM 46737 | 13.3 | [10.5 - 16.6] | 25.4 | [23.0 - 27.8] | 13.6 | [11.3 - 16.4] | 4.82 |
| **'BR-AD22'** | *Corynebacterium pseudopelargi* CCM 8832 | 13.2 | [10.5 - 16.6] | 25.4 | [23.1 - 27.9] | 13.6 | [11.2 - 16.4] | 4.51 |
| **'0211'** | *Corynebacterium pelargi* DSM 46737 | 13.3 | [10.5 - 16.6] | 25.4 | [23.0 - 27.8] | 13.6 | [11.3 - 16.4] | 4.82 |
| **'131002'** | *Corynebacterium kutscheri* DSM 20755 | 13.2 | [10.5 - 16.5] | 25.4 | [23.0 - 27.8] | 13.6 | [11.2 - 16.4] | 6.91 |
| **'BR-AD22'** | *Corynebacterium resistens* DSM 45100 | 12.9 | [10.2 - 16.2] | 25.4 | [23.0 - 27.8] | 13.3 | [10.9 - 16.1] | 3.69 |
| **'210932'** | *Corynebacterium kutscheri* DSM 20755 | 13.2 | [10.5 - 16.6] | 25.3 | [22.9 - 27.7] | 13.6 | [11.2 - 16.4] | 6.85 |
| **'04-3911'** | *Corynebacterium pelargi* DSM 46737 | 13.3 | [10.5 - 16.6] | 25.3 | [23.0 - 27.8] | 13.7 | [11.3 - 16.4] | 4.85 |
| **'2590'** | *Corynebacterium mustelae* DSM 45274 | 12.9 | [10.2 - 16.2] | 25.3 | [22.9 - 27.8] | 13.3 | [11.0 - 16.1] | 0.72 |

| Query | Subject | $d_0$ | C.I. $d_0$ | $d_4$ | C.I. $d_4$ | $d_6$ | C.I. $d_6$ | Diff. G+C Percent |
|---|---|---|---|---|---|---|---|---|
| **'04-7514'** | *Corynebacterium pelargi* DSM 46737 | 13.3 | [10.6 - 16.6] | 25.3 | [22.9 - 27.7] | 13.7 | [11.3 - 16.5] | 4.7 |
| **'05146'** | *Corynebacterium pseudopelargi* CCM 8832 | 13.3 | [10.6 - 16.6] | 25.2 | [22.9 - 27.7] | 13.7 | [11.3 - 16.5] | 4.6 |
| **'05-13'** | *Corynebacterium mustelae* DSM 45274 | 12.9 | [10.2 - 16.2] | 25.2 | [22.9 - 27.7] | 13.3 | [11.0 - 16.1] | 1.85 |
| **'06-19'** | *Corynebacterium mustelae* DSM 45274 | 13.0 | [10.2 - 16.2] | 25.2 | [22.9 - 27.7] | 13.3 | [11.0 - 16.1] | 0.76 |
| **'06-16'** | *Corynebacterium mustelae* DSM 45274 | 13.0 | [10.2 - 16.2] | 25.2 | [22.9 - 27.7] | 13.3 | [11.0 - 16.1] | 0.8 |
| **'05146'** | *Corynebacterium pelargi* DSM 46737 | 13.3 | [10.6 - 16.6] | 25.2 | [22.9 - 27.7] | 13.7 | [11.3 - 16.5] | 4.87 |
| **'02-13'** | *Corynebacterium mustelae* DSM 45274 | 13.0 | [10.2 - 16.2] | 25.2 | [22.8 - 27.6] | 13.3 | [11.0 - 16.1] | 0.77 |
| **'210931'** | *Corynebacterium pseudopelargi* CCM 8832 | 13.3 | [10.6 - 16.6] | 25.2 | [22.9 - 27.7] | 13.7 | [11.3 - 16.5] | 4.61 |
| **'210931'** | *Corynebacterium pelargi* DSM 46737 | 13.3 | [10.6 - 16.6] | 25.2 | [22.9 - 27.7] | 13.7 | [11.3 - 16.5] | 4.87 |
| **'03-8664'** | *Corynebacterium resistens* DSM 45100 | 12.9 | [10.2 - 16.2] | 25.2 | [22.9 - 27.7] | 13.3 | [11.0 - 16.1] | 3.47 |
| **'131002'** | *Corynebacterium humireducens* DSM 45392 | 13.1 | [10.3 - 16.4] | 25.2 | [22.9 - 27.7] | 13.4 | [11.1 - 16.2] | 15.19 |
| **'04-15'** | *Corynebacterium mustelae* DSM 45274 | 13.0 | [10.2 - 16.2] | 25.2 | [22.9 - 27.7] | 13.3 | [11.0 - 16.1] | 0.75 |
| **'04-7514'** | *Corynebacterium resistens* DSM 45100 | 12.9 | [10.2 - 16.2] | 25.1 | [22.8 - 27.6] | 13.3 | [10.9 - 16.1] | 3.62 |
| **'809'** | *Corynebacterium humireducens* DSM 45392 | 13.1 | [10.3 - 16.4] | 25.1 | [22.7 - 27.5] | 13.4 | [11.1 - 16.2] | 15.27 |
| **'131001'** | *Corynebacterium kutscheri* DSM 20755 | 13.3 | [10.5 - 16.6] | 25.1 | [22.8 - 27.6] | 13.6 | [11.2 - 16.4] | 6.85 |
| **'04-7514'** | *Corynebacterium pseudopelargi* CCM 8832 | 13.3 | [10.6 - 16.6] | 25.1 | [22.8 - 27.6] | 13.7 | [11.3 - 16.4] | 4.44 |
| **'03-8664'** | *Corynebacterium humireducens* DSM 45392 | 13.1 | [10.4 - 16.4] | 25.1 | [22.8 - 27.6] | 13.5 | [11.1 - 16.2] | 14.95 |
| **'4940'** | *Corynebacterium mustelae* DSM 45274 | 13.0 | [10.3 - 16.3] | 25.1 | [22.8 - 27.6] | 13.3 | [11.0 - 16.1] | 0.77 |
| **'210931'** | *Corynebacterium humireducens* DSM 45392 | 13.1 | [10.3 - 16.4] | 25.1 | [22.7 - 27.5] | 13.4 | [11.1 - 16.2] | 15.27 |
| **'03-8664'** | *Corynebacterium vitaeruminis* DSM 20294 | 13.3 | [10.6 - 16.6] | 25.1 | [22.8 - 27.6] | 13.7 | [11.3 - 16.5] | 11.9 |
| **'04-13'** | *Corynebacterium mustelae* DSM 45274 | 12.9 | [10.2 - 16.2] | 25.1 | [22.8 - 27.6] | 13.3 | [11.0 - 16.1] | 1.86 |
| **'05146'** | *Corynebacterium kutscheri* DSM 20755 | 13.2 | [10.5 - 16.6] | 25.0 | [22.7 - 27.5] | 13.6 | [11.2 - 16.4] | 6.84 |
| **'210931'** | *Corynebacterium kutscheri* DSM 20755 | 13.2 | [10.5 - 16.6] | 25.0 | [22.7 - 27.5] | 13.6 | [11.2 - 16.4] | 6.84 |
| **'04-7514'** | *Corynebacterium humireducens* DSM 45392 | 13.1 | [10.4 - 16.4] | 25.0 | [22.7 - 27.5] | 13.4 | [11.1 - 16.2] | 15.1 |
| **'809'** | *Corynebacterium resistens* DSM 45100 | 12.9 | [10.2 - 16.2] | 25.0 | [22.7 - 27.5] | 13.3 | [10.9 - 16.1] | 3.79 |

| Query | Subject | $d_0$ | C.I. $d_0$ | $d_4$ | C.I. $d_4$ | $d_6$ | C.I. $d_6$ | Diff. G+C Percent |
|---|---|---|---|---|---|---|---|---|
| **'05146'** | *Corynebacterium humireducens* DSM 45392 | 13.1 | [10.3 - 16.4] | 25.0 | [22.7 - 27.5] | 13.4 | [11.1 - 16.2] | 15.27 |
| **'210932'** | *Corynebacterium pseudopelargi* CCM 8832 | 13.3 | [10.6 - 16.6] | 24.9 | [22.6 - 27.4] | 13.7 | [11.3 - 16.5] | 4.59 |
| **'BR-AD22'** | *Corynebacterium humireducens* DSM 45392 | 13.0 | [10.3 - 16.3] | 24.9 | [22.6 - 27.4] | 13.4 | [11.1 - 16.2] | 15.17 |
| **'131001'** | *Corynebacterium pseudopelargi* CCM 8832 | 13.3 | [10.6 - 16.6] | 24.9 | [22.6 - 27.4] | 13.7 | [11.3 - 16.5] | 4.6 |
| **'04-3911'** | *Corynebacterium kutscheri* DSM 20755 | 13.3 | [10.5 - 16.6] | 24.8 | [22.5 - 27.3] | 13.6 | [11.3 - 16.4] | 6.86 |
| **'0102'** | *Corynebacterium resistens* DSM 45100 | 12.9 | [10.2 - 16.2] | 24.8 | [22.4 - 27.2] | 13.3 | [11.0 - 16.1] | 3.73 |
| **'131001'** | *Corynebacterium resistens* DSM 45100 | 12.9 | [10.2 - 16.2] | 24.8 | [22.5 - 27.3] | 13.3 | [11.0 - 16.1] | 3.78 |
| **'04-13'** | *Corynebacterium resistens* DSM 45100 | 13.1 | [10.4 - 16.4] | 24.8 | [22.5 - 27.3] | 13.5 | [11.1 - 16.3] | 2.66 |
| **'4940'** | *Corynebacterium pseudopelargi* CCM 8832 | 13.2 | [10.5 - 16.5] | 24.8 | [22.5 - 27.3] | 13.6 | [11.2 - 16.4] | 4.56 |
| **'06-19'** | *Corynebacterium vitaeruminis* DSM 20294 | 13.2 | [10.5 - 16.5] | 24.8 | [22.4 - 27.2] | 13.6 | [11.2 - 16.4] | 12.2 |
| **'06-16'** | *Corynebacterium vitaeruminis* DSM 20294 | 13.2 | [10.5 - 16.5] | 24.8 | [22.5 - 27.3] | 13.6 | [11.2 - 16.4] | 12.15 |
| **'0211'** | *Corynebacterium resistens* DSM 45100 | 12.9 | [10.2 - 16.2] | 24.8 | [22.4 - 27.2] | 13.3 | [11.0 - 16.1] | 3.73 |
| **'04-15'** | *Corynebacterium vitaeruminis* DSM 20294 | 13.2 | [10.5 - 16.5] | 24.8 | [22.4 - 27.2] | 13.6 | [11.2 - 16.4] | 12.2 |
| **'03-8664'** | *Corynebacterium pseudopelargi* CCM 8832 | 13.3 | [10.6 - 16.7] | 24.8 | [22.5 - 27.3] | 13.7 | [11.3 - 16.5] | 4.28 |
| **'4940'** | *Corynebacterium vitaeruminis* DSM 20294 | 13.2 | [10.5 - 16.5] | 24.8 | [22.5 - 27.3] | 13.6 | [11.2 - 16.4] | 12.18 |
| **'131001'** | *Corynebacterium pelargi* DSM 46737 | 13.3 | [10.6 - 16.7] | 24.8 | [22.5 - 27.2] | 13.7 | [11.3 - 16.5] | 4.86 |
| **'02-13'** | *Corynebacterium vitaeruminis* DSM 20294 | 13.2 | [10.5 - 16.5] | 24.8 | [22.5 - 27.3] | 13.6 | [11.2 - 16.4] | 12.19 |
| **'210932'** | *Corynebacterium pelargi* DSM 46737 | 13.3 | [10.6 - 16.7] | 24.8 | [22.5 - 27.3] | 13.7 | [11.3 - 16.5] | 4.86 |
| **'04-3911'** | *Corynebacterium humireducens* DSM 45392 | 13.1 | [10.3 - 16.3] | 24.8 | [22.4 - 27.2] | 13.4 | [11.1 - 16.2] | 15.25 |
| **'03-8664'** | *Corynebacterium kutscheri* DSM 20755 | 13.2 | [10.5 - 16.6] | 24.8 | [22.4 - 27.2] | 13.6 | [11.2 - 16.4] | 7.16 |
| **'809'** | *Corynebacterium pseudopelargi* CCM 8832 | 13.3 | [10.6 - 16.7] | 24.8 | [22.5 - 27.3] | 13.7 | [11.3 - 16.5] | 4.6 |
| **'210932'** | *Corynebacterium resistens* DSM 45100 | 12.9 | [10.2 - 16.2] | 24.8 | [22.4 - 27.2] | 13.3 | [11.0 - 16.1] | 3.78 |
| **'04-7514'** | *Corynebacterium kutscheri* DSM 20755 | 13.3 | [10.5 - 16.6] | 24.7 | [22.4 - 27.2] | 13.6 | [11.2 - 16.4] | 7.0 |
| **'809'** | *Corynebacterium phocae* DSM 44612 | 12.9 | [10.2 - 16.2] | 24.7 | [22.4 - 27.2] | 13.3 | [11.0 - 16.1] | 5.51 |
| **'03-8664'** | *Corynebacterium pelargi* DSM 46737 | 13.4 | [10.7 - 16.7] | 24.7 | [22.4 - 27.2] | 13.8 | [11.4 - 16.6] | 4.55 |
| **'BR-AD22'** | *Corynebacterium kutscheri* DSM 20755 | 13.3 | [10.6 - 16.6] | 24.7 | [22.3 - 27.1] | 13.7 | [11.3 - 16.4] | 6.94 |

| Query | Subject | $d_0$ | C.I. $d_0$ | $d_4$ | C.I. $d_4$ | $d_6$ | C.I. $d_6$ | Diff. G+C Percent |
|---|---|---|---|---|---|---|---|---|
| **'04-3911'** | *Corynebacterium resistens* DSM 45100 | 12.9 | [10.2 - 16.2] | 24.7 | [22.4 - 27.2] | 13.3 | [11.0 - 16.1] | 3.77 |
| **'0102'** | *Corynebacterium humireducens* DSM 45392 | 13.1 | [10.3 - 16.3] | 24.7 | [22.4 - 27.1] | 13.4 | [11.1 - 16.2] | 15.21 |
| **'0211'** | *Corynebacterium humireducens* DSM 45392 | 13.1 | [10.3 - 16.3] | 24.7 | [22.4 - 27.2] | 13.4 | [11.1 - 16.2] | 15.21 |
| **'04-15'** | *Corynebacterium pelargi* DSM 46737 | 13.2 | [10.5 - 16.5] | 24.7 | [22.4 - 27.2] | 13.6 | [11.2 - 16.4] | 4.85 |
| **'06-19'** | *Corynebacterium pelargi* DSM 46737 | 13.2 | [10.5 - 16.5] | 24.7 | [22.4 - 27.2] | 13.6 | [11.2 - 16.4] | 4.84 |
| **'210932'** | *Corynebacterium phocae* DSM 44612 | 12.9 | [10.2 - 16.2] | 24.6 | [22.3 - 27.1] | 13.3 | [11.0 - 16.1] | 5.5 |
| **'04-15'** | *Corynebacterium pseudopelargi* CCM 8832 | 13.2 | [10.5 - 16.5] | 24.6 | [22.3 - 27.1] | 13.6 | [11.2 - 16.4] | 4.58 |
| **'131001'** | *Corynebacterium phocae* DSM 44612 | 12.9 | [10.2 - 16.2] | 24.6 | [22.3 - 27.1] | 13.3 | [11.0 - 16.1] | 5.5 |
| **'06-19'** | *Corynebacterium pseudopelargi* CCM 8832 | 13.2 | [10.5 - 16.5] | 24.6 | [22.3 - 27.1] | 13.6 | [11.2 - 16.4] | 4.58 |
| **'2590'** | *Corynebacterium vitaeruminis* DSM 20294 | 13.2 | [10.5 - 16.5] | 24.6 | [22.3 - 27.0] | 13.6 | [11.2 - 16.4] | 12.23 |
| **'0102'** | *Corynebacterium kutscheri* DSM 20755 | 13.3 | [10.6 - 16.6] | 24.5 | [22.2 - 27.0] | 13.7 | [11.3 - 16.5] | 6.89 |
| **'03-8664'** | *Corynebacterium phocae* DSM 44612 | 13.0 | [10.2 - 16.2] | 24.5 | [22.2 - 27.0] | 13.3 | [11.0 - 16.1] | 5.19 |
| **'0211'** | *Corynebacterium kutscheri* DSM 20755 | 13.3 | [10.6 - 16.6] | 24.5 | [22.2 - 27.0] | 13.7 | [11.3 - 16.5] | 6.89 |
| **'02-13'** | *Corynebacterium pseudopelargi* CCM 8832 | 13.2 | [10.5 - 16.6] | 24.5 | [22.2 - 26.9] | 13.6 | [11.2 - 16.4] | 4.57 |
| **'210932'** | *Corynebacterium humireducens* DSM 45392 | 13.1 | [10.4 - 16.4] | 24.5 | [22.2 - 27.0] | 13.5 | [11.1 - 16.2] | 15.26 |
| **'131001'** | *Corynebacterium humireducens* DSM 45392 | 13.1 | [10.4 - 16.4] | 24.5 | [22.2 - 27.0] | 13.5 | [11.1 - 16.2] | 15.26 |
| **'06-16'** | *Corynebacterium pelargi* DSM 46737 | 13.3 | [10.5 - 16.6] | 24.4 | [22.1 - 26.9] | 13.6 | [11.2 - 16.4] | 4.8 |
| **'02-13'** | *Corynebacterium pelargi* DSM 46737 | 13.3 | [10.5 - 16.6] | 24.4 | [22.1 - 26.9] | 13.6 | [11.2 - 16.4] | 4.83 |
| **'05-13'** | *Corynebacterium humireducens* DSM 45392 | 13.0 | [10.3 - 16.3] | 24.4 | [22.1 - 26.8] | 13.4 | [11.0 - 16.2] | 14.15 |
| **'2590'** | *Corynebacterium pseudopelargi* CCM 8832 | 13.2 | [10.5 - 16.5] | 24.4 | [22.1 - 26.9] | 13.6 | [11.2 - 16.4] | 4.61 |
| **'131002'** | *Corynebacterium pseudopelargi* CCM 8832 | 13.3 | [10.6 - 16.7] | 24.4 | [22.1 - 26.9] | 13.7 | [11.3 - 16.5] | 4.53 |
| **'131002'** | *Corynebacterium pelargi* DSM 46737 | 13.4 | [10.6 - 16.7] | 24.4 | [22.1 - 26.9] | 13.7 | [11.4 - 16.5] | 4.8 |
| **'04-13'** | *Corynebacterium vitaeruminis* DSM 20294 | 13.4 | [10.7 - 16.8] | 24.4 | [22.1 - 26.9] | 13.8 | [11.4 - 16.6] | 11.09 |
| **'06-19'** | *Corynebacterium kutscheri* DSM 20755 | 13.2 | [10.5 - 16.5] | 24.3 | [22.0 - 26.8] | 13.6 | [11.2 - 16.3] | 6.87 |
| **'04-13'** | *Corynebacterium humireducens* DSM 45392 | 13.0 | [10.3 - 16.3] | 24.3 | [22.0 - 26.7] | 13.4 | [11.0 - 16.2] | 14.14 |

| Query | Subject | $d_0$ | C.I. $d_0$ | $d_4$ | C.I. $d_4$ | $d_6$ | C.I. $d_6$ | Diff. G+C Percent |
|---|---|---|---|---|---|---|---|---|
| **'04-15'** | *Corynebacterium kutscheri* DSM 20755 | 13.2 | [10.5 - 16.5] | 24.3 | [22.0 - 26.8] | 13.6 | [11.2 - 16.3] | 6.86 |
| **'06-16'** | *Corynebacterium pseudopelargi* CCM 8832 | 13.3 | [10.5 - 16.6] | 24.3 | [22.0 - 26.8] | 13.6 | [11.2 - 16.4] | 4.53 |
| **'210931'** | *Corynebacterium resistens* DSM 45100 | 12.9 | [10.2 - 16.2] | 24.3 | [22.0 - 26.8] | 13.3 | [11.0 - 16.1] | 3.79 |
| **'131002'** | *Corynebacterium resistens* DSM 45100 | 12.9 | [10.2 - 16.2] | 24.3 | [22.0 - 26.7] | 13.3 | [11.0 - 16.1] | 3.72 |
| **'809'** | *Corynebacterium kutscheri* DSM 20755 | 13.3 | [10.6 - 16.7] | 24.3 | [22.0 - 26.8] | 13.7 | [11.3 - 16.5] | 6.84 |
| **'BR-AD22'** | *Corynebacterium phocae* DSM 44612 | 12.9 | [10.2 - 16.2] | 24.3 | [22.0 - 26.8] | 13.3 | [11.0 - 16.1] | 5.41 |
| **'05-13'** | *Corynebacterium pseudopelargi* CCM 8832 | 13.2 | [10.5 - 16.5] | 24.2 | [21.9 - 26.7] | 13.6 | [11.2 - 16.4] | 3.49 |
| **'04-13'** | *Corynebacterium pseudopelargi* CCM 8832 | 13.2 | [10.5 - 16.5] | 24.2 | [21.8 - 26.6] | 13.6 | [11.2 - 16.4] | 3.47 |
| **'04-7514'** | *Corynebacterium phocae* DSM 44612 | 13.0 | [10.3 - 16.3] | 24.2 | [21.9 - 26.7] | 13.3 | [11.0 - 16.1] | 5.34 |
| **'0102'** | *Corynebacterium phocae* DSM 44612 | 12.9 | [10.2 - 16.2] | 24.1 | [21.8 - 26.6] | 13.3 | [11.0 - 16.1] | 5.45 |
| **'0211'** | *Corynebacterium phocae* DSM 44612 | 12.9 | [10.2 - 16.2] | 24.1 | [21.8 - 26.6] | 13.3 | [11.0 - 16.1] | 5.45 |
| **'06-16'** | *Corynebacterium humireducens* DSM 45392 | 13.0 | [10.3 - 16.3] | 24.1 | [21.8 - 26.6] | 13.4 | [11.0 - 16.2] | 15.2 |
| **'210931'** | *Corynebacterium phocae* DSM 44612 | 13.0 | [10.2 - 16.2] | 24.1 | [21.8 - 26.6] | 13.3 | [11.0 - 16.1] | 5.51 |
| **'2590'** | *Corynebacterium humireducens* DSM 45392 | 13.0 | [10.3 - 16.3] | 24.1 | [21.8 - 26.6] | 13.4 | [11.0 - 16.1] | 15.28 |
| **'05-13'** | *Corynebacterium kutscheri* DSM 20755 | 13.1 | [10.4 - 16.4] | 24.1 | [21.8 - 26.6] | 13.5 | [11.1 - 16.2] | 7.95 |
| **'02-13'** | *Corynebacterium humireducens* DSM 45392 | 13.0 | [10.3 - 16.3] | 24.1 | [21.8 - 26.6] | 13.4 | [11.0 - 16.2] | 15.23 |
| **'131002'** | *Corynebacterium phocae* DSM 44612 | 13.0 | [10.2 - 16.2] | 24.1 | [21.8 - 26.6] | 13.3 | [11.0 - 16.1] | 5.43 |
| **'04-3911'** | *Corynebacterium phocae* DSM 44612 | 12.9 | [10.2 - 16.2] | 24.1 | [21.8 - 26.6] | 13.3 | [11.0 - 16.1] | 5.49 |
| **'04-13'** | *Corynebacterium kutscheri* DSM 20755 | 13.1 | [10.4 - 16.4] | 24.0 | [21.7 - 26.4] | 13.5 | [11.1 - 16.2] | 7.97 |
| **'06-16'** | *Corynebacterium kutscheri* DSM 20755 | 13.2 | [10.5 - 16.5] | 24.0 | [21.7 - 26.4] | 13.5 | [11.2 - 16.3] | 6.91 |
| **'809'** | *Corynebacterium pelargi* DSM 46737 | 13.4 | [10.6 - 16.7] | 24.0 | [21.7 - 26.5] | 13.7 | [11.4 - 16.5] | 4.87 |
| **'03-8664'** | *Corynebacterium rouxii* FRC0190 T | 14.5 | [11.6 - 17.9] | 24.0 | [21.6 - 26.4] | 14.7 | [12.3 - 17.6] | 0.4 |
| **'02-13'** | *Corynebacterium kutscheri* DSM 20755 | 13.2 | [10.5 - 16.5] | 23.9 | [21.6 - 26.4] | 13.5 | [11.2 - 16.3] | 6.88 |
| **'06-19'** | *Corynebacterium humireducens* DSM 45392 | 13.0 | [10.3 - 16.3] | 23.9 | [21.6 - 26.4] | 13.4 | [11.0 - 16.1] | 15.24 |
| **'04-15'** | *Corynebacterium humireducens* DSM 45392 | 13.0 | [10.3 - 16.3] | 23.9 | [21.6 - 26.4] | 13.4 | [11.0 - 16.1] | 15.25 |

| Query | Subject | $d_0$ | C.I. $d_0$ | $d_4$ | C.I. $d_4$ | $d_6$ | C.I. $d_6$ | Diff. G+C Percent |
|---|---|---|---|---|---|---|---|---|
| **'03-8664'** | *Corynebacterium diphtheriae* subsp. *lausannense* CHUV2995 | 14.3 | [11.4 - 17.6] | 23.9 | [21.6 - 26.4] | 14.5 | [12.1 - 17.4] | 0.32 |
| **'4940'** | *Corynebacterium pelargi* DSM 46737 | 13.3 | [10.5 - 16.6] | 23.9 | [21.6 - 26.4] | 13.6 | [11.3 - 16.4] | 4.83 |
| **'03-8664'** | *Corynebacterium diphtheriae* NCTC 11397 | 14.2 | [11.4 - 17.6] | 23.9 | [21.6 - 26.4] | 14.5 | [12.0 - 17.3] | 0.1 |
| **'05146'** | *Corynebacterium phocae* DSM 44612 | 13.0 | [10.3 - 16.2] | 23.9 | [21.6 - 26.3] | 13.3 | [11.0 - 16.1] | 5.51 |
| **'04-15'** | *Corynebacterium resistens* DSM 45100 | 12.9 | [10.2 - 16.2] | 23.9 | [21.6 - 26.4] | 13.3 | [10.9 - 16.0] | 3.77 |
| **'06-19'** | *Corynebacterium resistens* DSM 45100 | 12.9 | [10.2 - 16.2] | 23.9 | [21.6 - 26.4] | 13.3 | [10.9 - 16.0] | 3.76 |
| **'2590'** | *Corynebacterium kutscheri* DSM 20755 | 13.2 | [10.5 - 16.5] | 23.9 | [21.6 - 26.4] | 13.6 | [11.2 - 16.4] | 6.83 |
| **'04-7514'** | *Corynebacterium diphtheriae* NCTC 11397 | 14.2 | [11.4 - 17.6] | 23.8 | [21.5 - 26.3] | 14.5 | [12.0 - 17.3] | 0.06 |
| **'05-13'** | *Corynebacterium diphtheriae* subsp. *lausannense* CHUV2995 | 14.3 | [11.5 - 17.7] | 23.8 | [21.5 - 26.3] | 14.6 | [12.2 - 17.5] | 0.48 |
| **'04-13'** | *Corynebacterium diphtheriae* subsp. *lausannense* CHUV2995 | 14.4 | [11.5 - 17.7] | 23.7 | [21.4 - 26.2] | 14.6 | [12.2 - 17.5] | 0.49 |
| **'4940'** | *Corynebacterium humireducens* DSM 45392 | 13.0 | [10.3 - 16.3] | 23.7 | [21.4 - 26.1] | 13.4 | [11.0 - 16.1] | 15.23 |
| **'2590'** | *Corynebacterium pelargi* DSM 46737 | 13.3 | [10.6 - 16.6] | 23.6 | [21.3 - 26.0] | 13.7 | [11.3 - 16.4] | 4.88 |
| **'05-13'** | *Corynebacterium vitaeruminis* DSM 20294 | 13.5 | [10.8 - 16.9] | 23.6 | [21.3 - 26.0] | 13.9 | [11.5 - 16.7] | 11.11 |
| **'04-7514'** | *Corynebacterium diphtheriae* subsp. *lausannense* CHUV2995 | 14.2 | [11.4 - 17.6] | 23.6 | [21.3 - 26.1] | 14.5 | [12.1 - 17.4] | 0.47 |
| **'2590'** | *Corynebacterium resistens* DSM 45100 | 12.9 | [10.2 - 16.1] | 23.6 | [21.3 - 26.1] | 13.2 | [10.9 - 16.0] | 3.8 |
| **'BR-AD22'** | *Corynebacterium diphtheriae* NCTC 11397 | 14.2 | [11.4 - 17.6] | 23.6 | [21.3 - 26.1] | 14.5 | [12.1 - 17.3] | 0.12 |
| **'210931'** | *Corynebacterium diphtheriae* NCTC 11397 | 14.3 | [11.4 - 17.6] | 23.5 | [21.2 - 25.9] | 14.5 | [12.1 - 17.4] | 0.22 |
| **'4940'** | *Corynebacterium kutscheri* DSM 20755 | 13.2 | [10.5 - 16.5] | 23.5 | [21.2 - 26.0] | 13.6 | [11.2 - 16.4] | 6.88 |
| **'0211'** | *Corynebacterium diphtheriae* NCTC 11397 | 14.3 | [11.4 - 17.7] | 23.5 | [21.2 - 26.0] | 14.5 | [12.1 - 17.4] | 0.17 |
| **'0102'** | *Corynebacterium diphtheriae* NCTC 11397 | 14.3 | [11.4 - 17.7] | 23.5 | [21.2 - 26.0] | 14.5 | [12.1 - 17.4] | 0.17 |
| **'04-3911'** | *Corynebacterium diphtheriae* NCTC 11397 | 14.3 | [11.5 - 17.7] | 23.5 | [21.2 - 25.9] | 14.6 | [12.1 - 17.4] | 0.2 |
| **'04-3911'** | *Corynebacterium rouxii* FRC0190 T | 14.6 | [11.7 - 18.0] | 23.5 | [21.2 - 26.0] | 14.8 | [12.4 - 17.7] | 0.09 |
| **'131001'** | *Corynebacterium diphtheriae* NCTC 11397 | 14.2 | [11.4 - 17.6] | 23.4 | [21.2 - 25.9] | 14.5 | [12.1 - 17.4] | 0.21 |
| **'05146'** | *Corynebacterium diphtheriae* NCTC 11397 | 14.3 | [11.4 - 17.6] | 23.4 | [21.1 - 25.9] | 14.5 | [12.1 - 17.4] | 0.22 |
| **'210932'** | *Corynebacterium diphtheriae* NCTC 11397 | 14.2 | [11.4 - 17.6] | 23.4 | [21.1 - 25.9] | 14.5 | [12.1 - 17.4] | 0.21 |
| **'0211'** | *Corynebacterium rouxii* FRC0190 T | 14.7 | [11.8 - 18.1] | 23.3 | [21.0 - 25.8] | 14.9 | [12.5 - 17.8] | 0.13 |

| Query | Subject | $d_0$ | C.I. $d_0$ | $d_4$ | C.I. $d_4$ | $d_6$ | C.I. $d_6$ | Diff. G+C Percent |
|---|---|---|---|---|---|---|---|---|
| **'809'** | *Corynebacterium diphtheriae* NCTC 11397 | 14.3 | [11.4 - 17.6] | 23.3 | [21.0 - 25.8] | 14.5 | [12.1 - 17.4] | 0.22 |
| **'04-7514'** | *Corynebacterium rouxii* FRC0190 T | 14.5 | [11.7 - 17.9] | 23.3 | [21.0 - 25.8] | 14.8 | [12.3 - 17.7] | 0.24 |
| **'05-13'** | *Corynebacterium rouxii* FRC0190 T | 14.3 | [11.5 - 17.7] | 23.3 | [21.0 - 25.7] | 14.6 | [12.1 - 17.4] | 1.19 |
| **'04-7514'** | *Corynebacterium belfantii* FRC0043 | 14.3 | [11.4 - 17.6] | 23.3 | [21.0 - 25.7] | 14.5 | [12.1 - 17.4] | 0.16 |
| **'06-16'** | *Corynebacterium resistens* DSM 45100 | 12.9 | [10.2 - 16.2] | 23.3 | [21.1 - 25.8] | 13.3 | [10.9 - 16.0] | 3.72 |
| **'4940'** | *Corynebacterium phocae* DSM 44612 | 12.9 | [10.2 - 16.2] | 23.3 | [21.0 - 25.8] | 13.3 | [10.9 - 16.0] | 5.47 |
| **'0102'** | *Corynebacterium rouxii* FRC0190 T | 14.7 | [11.8 - 18.1] | 23.3 | [21.0 - 25.8] | 14.9 | [12.5 - 17.8] | 0.13 |
| **'04-15'** | *Corynebacterium diphtheriae* NCTC 11397 | 14.2 | [11.4 - 17.6] | 23.2 | [20.9 - 25.6] | 14.5 | [12.0 - 17.3] | 0.2 |
| **'04-3911'** | *Corynebacterium diphtheriae* subsp. *lausannense* CHUV2995 | 14.4 | [11.6 - 17.8] | 23.2 | [20.9 - 25.7] | 14.7 | [12.2 - 17.5] | 0.62 |
| **'04-15'** | *Corynebacterium phocae* DSM 44612 | 12.9 | [10.2 - 16.2] | 23.2 | [20.9 - 25.7] | 13.3 | [10.9 - 16.0] | 5.49 |
| **'04-13'** | *Corynebacterium rouxii* FRC0190 T | 14.3 | [11.5 - 17.7] | 23.2 | [20.9 - 25.7] | 14.6 | [12.1 - 17.4] | 1.21 |
| **'06-19'** | *Corynebacterium phocae* DSM 44612 | 12.9 | [10.2 - 16.2] | 23.2 | [20.9 - 25.7] | 13.3 | [10.9 - 16.0] | 5.48 |
| **'210931'** | *Corynebacterium rouxii* FRC0190 T | 14.5 | [11.7 - 17.9] | 23.2 | [20.9 - 25.6] | 14.8 | [12.3 - 17.6] | 0.07 |
| **'03-8664'** | *Corynebacterium belfantii* FRC0043 | 14.3 | [11.5 - 17.7] | 23.2 | [21.0 - 25.7] | 14.6 | [12.1 - 17.4] | 0.0 |
| **'06-19'** | *Corynebacterium diphtheriae* NCTC 11397 | 14.2 | [11.4 - 17.6] | 23.2 | [20.9 - 25.6] | 14.5 | [12.0 - 17.3] | 0.2 |
| **'131001'** | *Corynebacterium rouxii* FRC0190 T | 14.4 | [11.6 - 17.8] | 23.2 | [20.9 - 25.6] | 14.7 | [12.2 - 17.6] | 0.08 |
| **'02-13'** | *Corynebacterium resistens* DSM 45100 | 12.9 | [10.2 - 16.2] | 23.2 | [20.9 - 25.7] | 13.3 | [10.9 - 16.0] | 3.75 |
| **'0211'** | *Corynebacterium diphtheriae* subsp. *lausannense* CHUV2995 | 14.4 | [11.6 - 17.8] | 23.1 | [20.8 - 25.5] | 14.6 | [12.2 - 17.5] | 0.58 |
| **'2590'** | *Corynebacterium phocae* DSM 44612 | 12.9 | [10.2 - 16.2] | 23.1 | [20.8 - 25.6] | 13.3 | [10.9 - 16.0] | 5.52 |
| **'0102'** | *Corynebacterium diphtheriae* subsp. *lausannense* CHUV2995 | 14.4 | [11.6 - 17.8] | 23.1 | [20.8 - 25.5] | 14.6 | [12.2 - 17.5] | 0.58 |
| **'BR-AD22'** | *Corynebacterium rouxii* FRC0190 T | 14.7 | [11.8 - 18.1] | 23.1 | [20.8 - 25.6] | 14.9 | [12.4 - 17.8] | 0.17 |
| **'131002'** | *Corynebacterium diphtheriae* NCTC 11397 | 14.3 | [11.5 - 17.7] | 23.1 | [20.8 - 25.5] | 14.6 | [12.1 - 17.4] | 0.15 |
| **'210932'** | *Corynebacterium rouxii* FRC0190 T | 14.5 | [11.6 - 17.9] | 23.1 | [20.8 - 25.5] | 14.7 | [12.3 - 17.6] | 0.09 |
| **'BR-AD22'** | *Corynebacterium diphtheriae* subsp. *lausannense* CHUV2995 | 14.3 | [11.4 - 17.7] | 23.0 | [20.7 - 25.4] | 14.5 | [12.1 - 17.4] | 0.54 |
| **'06-16'** | *Corynebacterium diphtheriae* NCTC 11397 | 14.2 | [11.4 - 17.6] | 23.0 | [20.7 - 25.4] | 14.5 | [12.0 - 17.3] | 0.15 |
| **'809'** | *Corynebacterium diphtheriae* subsp. *lausannense* CHUV2995 | 14.2 | [11.3 - 17.5] | 23.0 | [20.7 - 25.5] | 14.4 | [12.0 - 17.3] | 0.64 |

| Query | Subject | $d_0$ | C.I. $d_0$ | $d_4$ | C.I. $d_4$ | $d_6$ | C.I. $d_6$ | Diff. G+C Percent |
|---|---|---|---|---|---|---|---|---|
| **'05-13'** | *Corynebacterium phocae* DSM 44612 | 12.9 | [10.2 - 16.2] | 23.0 | [20.7 - 25.5] | 13.3 | [10.9 - 16.1] | 4.39 |
| **'2590'** | *Corynebacterium diphtheriae* NCTC 11397 | 14.2 | [11.4 - 17.6] | 23.0 | [20.7 - 25.4] | 14.5 | [12.0 - 17.3] | 0.23 |
| **'05-13'** | *Corynebacterium pelargi* DSM 46737 | 13.3 | [10.6 - 16.7] | 23.0 | [20.7 - 25.5] | 13.7 | [11.3 - 16.5] | 3.76 |
| **'02-13'** | *Corynebacterium phocae* DSM 44612 | 12.9 | [10.2 - 16.2] | 22.9 | [20.6 - 25.4] | 13.3 | [10.9 - 16.0] | 5.47 |
| **'210932'** | *Corynebacterium diphtheriae* subsp. *lausannense* CHUV2995 | 14.2 | [11.4 - 17.6] | 22.9 | [20.6 - 25.4] | 14.5 | [12.0 - 17.3] | 0.63 |
| **'04-3911'** | *Corynebacterium belfantii* FRC0043 | 14.4 | [11.6 - 17.8] | 22.9 | [20.6 - 25.3] | 14.7 | [12.2 - 17.5] | 0.31 |
| **'06-16'** | *Corynebacterium phocae* DSM 44612 | 12.9 | [10.2 - 16.2] | 22.9 | [20.6 - 25.4] | 13.3 | [10.9 - 16.0] | 5.44 |
| **'131001'** | *Corynebacterium diphtheriae* subsp. *lausannense* CHUV2995 | 14.2 | [11.4 - 17.6] | 22.9 | [20.6 - 25.4] | 14.5 | [12.0 - 17.3] | 0.63 |
| **'05146'** | *Corynebacterium rouxii* FRC0190 T | 14.5 | [11.6 - 17.9] | 22.9 | [20.6 - 25.4] | 14.7 | [12.3 - 17.6] | 0.08 |
| **'04-13'** | *Corynebacterium pelargi* DSM 46737 | 13.4 | [10.6 - 16.7] | 22.9 | [20.6 - 25.3] | 13.7 | [11.3 - 16.5] | 3.74 |
| **'04-13'** | *Corynebacterium phocae* DSM 44612 | 12.9 | [10.2 - 16.2] | 22.9 | [20.7 - 25.4] | 13.3 | [10.9 - 16.0] | 4.38 |
| **'4940'** | *Corynebacterium diphtheriae* NCTC 11397 | 14.2 | [11.4 - 17.6] | 22.9 | [20.6 - 25.4] | 14.5 | [12.0 - 17.3] | 0.18 |
| **'02-13'** | *Corynebacterium diphtheriae* NCTC 11397 | 14.2 | [11.4 - 17.6] | 22.9 | [20.7 - 25.4] | 14.5 | [12.0 - 17.3] | 0.19 |
| **'809'** | *Corynebacterium rouxii* FRC0190 T | 14.6 | [11.7 - 18.0] | 22.8 | [20.5 - 25.2] | 14.8 | [12.4 - 17.7] | 0.07 |
| **'06-16'** | *Corynebacterium rouxii* FRC0190 T | 14.5 | [11.6 - 17.9] | 22.8 | [20.5 - 25.2] | 14.7 | [12.3 - 17.6] | 0.15 |
| **'05146'** | *Corynebacterium diphtheriae* subsp. *lausannense* CHUV2995 | 14.2 | [11.4 - 17.6] | 22.8 | [20.6 - 25.3] | 14.5 | [12.0 - 17.3] | 0.64 |
| **'131002'** | *Corynebacterium rouxii* FRC0190 T | 14.4 | [11.6 - 17.8] | 22.7 | [20.4 - 25.1] | 14.7 | [12.2 - 17.5] | 0.15 |
| **'809'** | *Corynebacterium belfantii* FRC0043 | 14.2 | [11.4 - 17.6] | 22.7 | [20.4 - 25.1] | 14.5 | [12.0 - 17.3] | 0.32 |
| **'04-13'** | *Corynebacterium diphtheriae* NCTC 11397 | 14.3 | [11.4 - 17.7] | 22.7 | [20.4 - 25.2] | 14.5 | [12.1 - 17.4] | 0.91 |
| **'05-13'** | *Corynebacterium belfantii* FRC0043 | 14.2 | [11.4 - 17.6] | 22.7 | [20.4 - 25.1] | 14.5 | [12.0 - 17.3] | 0.79 |
| **'210931'** | *Corynebacterium diphtheriae* subsp. *lausannense* CHUV2995 | 14.3 | [11.5 - 17.7] | 22.7 | [20.5 - 25.2] | 14.6 | [12.1 - 17.4] | 0.64 |
| **'0211'** | *Corynebacterium belfantii* FRC0043 | 14.4 | [11.6 - 17.8] | 22.7 | [20.4 - 25.2] | 14.7 | [12.2 - 17.5] | 0.27 |
| **'4940'** | *Corynebacterium resistens* DSM 45100 | 12.9 | [10.2 - 16.2] | 22.7 | [20.4 - 25.1] | 13.3 | [10.9 - 16.0] | 3.75 |
| **'2590'** | *Corynebacterium rouxii* FRC0190 T | 14.4 | [11.6 - 17.8] | 22.7 | [20.4 - 25.2] | 14.7 | [12.2 - 17.5] | 0.07 |
| **'04-15'** | *Corynebacterium diphtheriae* subsp. *lausannense* CHUV2995 | 14.2 | [11.4 - 17.5] | 22.7 | [20.4 - 25.1] | 14.4 | [12.0 - 17.3] | 0.62 |
| **'05-13'** | *Corynebacterium diphtheriae* NCTC 11397 | 14.3 | [11.5 - 17.7] | 22.7 | [20.5 - 25.2] | 14.5 | [12.1 - 17.4] | 0.89 |

| Query | Subject | $d_0$ | C.I. $d_0$ | $d_4$ | C.I. $d_4$ | $d_6$ | C.I. $d_6$ | Diff. G+C Percent |
|---|---|---|---|---|---|---|---|---|
| **'0102'** | *Corynebacterium belfantii* FRC0043 | 14.4 | [11.6 - 17.8] | 22.7 | [20.4 - 25.2] | 14.7 | [12.2 - 17.5] | 0.27 |
| **'02-13'** | *Corynebacterium diphtheriae* subsp. *lausannense* CHUV2995 | 14.2 | [11.4 - 17.6] | 22.6 | [20.3 - 25.1] | 14.5 | [12.0 - 17.3] | 0.6 |
| **'04-13'** | *Corynebacterium belfantii* FRC0043 | 14.2 | [11.4 - 17.6] | 22.6 | [20.4 - 25.1] | 14.5 | [12.0 - 17.3] | 0.81 |
| **'04-15'** | *Corynebacterium rouxii* FRC0190 T | 14.3 | [11.5 - 17.7] | 22.6 | [20.4 - 25.1] | 14.6 | [12.2 - 17.4] | 0.1 |
| **'BR-AD22'** | *Corynebacterium belfantii* FRC0043 | 14.3 | [11.5 - 17.7] | 22.6 | [20.3 - 25.0] | 14.6 | [12.1 - 17.4] | 0.23 |
| **'02-13'** | *Corynebacterium rouxii* FRC0190 T | 14.4 | [11.6 - 17.8] | 22.6 | [20.3 - 25.0] | 14.7 | [12.2 - 17.5] | 0.11 |
| **'06-19'** | *Corynebacterium rouxii* FRC0190 T | 14.3 | [11.5 - 17.7] | 22.6 | [20.4 - 25.1] | 14.6 | [12.2 - 17.4] | 0.1 |
| **'06-19'** | *Corynebacterium diphtheriae* subsp. *lausannense* CHUV2995 | 14.2 | [11.4 - 17.5] | 22.6 | [20.4 - 25.1] | 14.4 | [12.0 - 17.3] | 0.61 |
| **'4940'** | *Corynebacterium rouxii* FRC0190 T | 14.3 | [11.5 - 17.7] | 22.6 | [20.3 - 25.1] | 14.5 | [12.1 - 17.4] | 0.11 |
| **'05146'** | *Corynebacterium belfantii* FRC0043 | 14.2 | [11.4 - 17.6] | 22.4 | [20.1 - 24.8] | 14.5 | [12.1 - 17.3] | 0.32 |
| **'210932'** | *Corynebacterium belfantii* FRC0043 | 14.3 | [11.4 - 17.6] | 22.4 | [20.1 - 24.8] | 14.5 | [12.1 - 17.4] | 0.31 |
| **'131002'** | *Corynebacterium diphtheriae* subsp. *lausannense* CHUV2995 | 14.3 | [11.5 - 17.7] | 22.4 | [20.1 - 24.9] | 14.5 | [12.1 - 17.4] | 0.56 |
| **'131001'** | *Corynebacterium belfantii* FRC0043 | 14.3 | [11.4 - 17.6] | 22.4 | [20.1 - 24.8] | 14.5 | [12.1 - 17.4] | 0.31 |
| **'210931'** | *Corynebacterium belfantii* FRC0043 | 14.4 | [11.5 - 17.8] | 22.3 | [20.0 - 24.7] | 14.6 | [12.2 - 17.5] | 0.33 |
| **'4940'** | *Corynebacterium diphtheriae* subsp. *lausannense* CHUV2995 | 14.2 | [11.4 - 17.6] | 22.3 | [20.0 - 24.8] | 14.4 | [12.0 - 17.3] | 0.6 |
| **'2590'** | *Corynebacterium diphtheriae* subsp. *lausannense* CHUV2995 | 14.2 | [11.4 - 17.6] | 22.3 | [20.1 - 24.8] | 14.5 | [12.0 - 17.3] | 0.65 |
| **'06-16'** | *Corynebacterium diphtheriae* subsp. *lausannense* CHUV2995 | 14.2 | [11.4 - 17.6] | 22.3 | [20.0 - 24.8] | 14.5 | [12.0 - 17.3] | 0.57 |
| **'04-15'** | *Corynebacterium belfantii* FRC0043 | 14.2 | [11.4 - 17.6] | 22.2 | [19.9 - 24.6] | 14.5 | [12.0 - 17.3] | 0.3 |
| **'06-19'** | *Corynebacterium belfantii* FRC0043 | 14.2 | [11.4 - 17.6] | 22.2 | [19.9 - 24.6] | 14.5 | [12.0 - 17.3] | 0.3 |
| **'02-13'** | *Corynebacterium belfantii* FRC0043 | 14.2 | [11.4 - 17.6] | 22.0 | [19.7 - 24.4] | 14.5 | [12.0 - 17.3] | 0.29 |
| **'131002'** | *Corynebacterium belfantii* FRC0043 | 14.3 | [11.5 - 17.7] | 21.9 | [19.6 - 24.3] | 14.6 | [12.1 - 17.4] | 0.25 |
| **'4940'** | *Corynebacterium belfantii* FRC0043 | 14.2 | [11.4 - 17.6] | 21.8 | [19.5 - 24.2] | 14.5 | [12.0 - 17.3] | 0.28 |
| **'2590'** | *Corynebacterium belfantii* FRC0043 | 14.3 | [11.4 - 17.6] | 21.8 | [19.6 - 24.3] | 14.5 | [12.1 - 17.3] | 0.33 |
| **'06-16'** | *Corynebacterium belfantii* FRC0043 | 14.3 | [11.5 - 17.7] | 21.8 | [19.5 - 24.2] | 14.5 | [12.1 - 17.4] | 0.25 |

#### Table 4: Strains in your dataset

Joint dataset of automatically determined closest type strains (if this mode was chosen), manually selected type strains (if selected accordingly) and the provided user strains, if provided (marked in yellow).

| Strain | Authority | Other deposits | Synonyms | Base pairs | Percent G+C | No. proteins | Goldstamp | Bioproject accession | Biosample accession | Assembly accession | IMG OID |
|---|---|---|---|---|---|---|---|---|---|---|---|
| *Corynebacterium vitaeruminis* DSM 20294 | (Bechdel et al. 1928) Lanéelle et al. 1980 | CCUG 28792; ATCC 10234; JCM 1323; IFO 12143; NBRC 12143; VKM B-1211; CIP 82.07; NCIB 9291; NCIMB 9291 | *Brevibacterium vitaeruminis*; *Corynebacterium vitaeruminis*; *Flavobacterium vitarumen* | 2931 780 | 65.5 | 2577 | Gp0023683 | PRJNA172966 | SAMN03081455 | GCA_000550805 | 2558860221 |
| *Corynebacterium diphtheriae* subsp. *lausannense* CHUV2995 | Tagini et al. 2019 | CCUG 72509; DSM 107520 | *Corynebacterium diphtheriae* subsp. *lausannense* | 3060 363 | 53.9 | 3145 | Gp0442955 | PRJEB24256 | SAMEA104679569 | GCA_900312965 | |
| *Corynebacterium belfantii* FRC0043 | Dazas et al. 2018 | DSM 105776; CIP 111412 | *Corynebacterium belfantii* | 2598 827 | 53.6 | 2557 | Gp0364753 | PRJEB22103 | SAMEA104208677 | GCA_900205605 | |
| *Corynebacterium phocae* DSM 44612 | Pascual et al. 1998 | CCUG 38205; JCM 12105; CIP 105741; strain M408/89/1 | *Corynebacterium phocae* | 2791 853 | 58.8 | 2246 | Gp0118686 | PRJNA242582 | SAMN02996498 | GCA_001941565 | |
| *Corynebacterium silvaticum* KL0182 | Dangel et al. 2020 | LMG 31313; DSM 109166; CIP 111672 | *Corynebacterium silvaticum* | 2548 487 | 54.4 | 2017 | | PRJNA517029 | SAMN10039578 | GCA_004382825 | |
| *Corynebacterium resistens* DSM 45100 | Otsuka et al. 2005 emend. Nouioui et al. 2018 | CCUG 50093; JCM 12819; GTC 2026; SICGH 158 | *Corynebacterium resistens* | 2601 311 | 57.1 | 2171 | Gp0003835 | PRJNA39683 | SAMN02603065 | GCA_000177535 | 650716029 |

| Strain | Authority | Other deposits | Synonyms | Base pairs | Percent G+C | No. proteins | Goldstamp | Bioproject accession | Biosample accession | Assembly accession | IMG OID |
|---|---|---|---|---|---|---|---|---|---|---|---|
| *Corynebacterium pseudopelargi* CCM 8832 | Busse et al. 2019 | 812CH; LMG 30627; CCUG 72167 | *Corynebacterium pseudopelargi* | 2348 160 | 57.9 | 2199 | Gp0379416 | PRJNA224116 | SAMN08449372 | GCF_003814005 | |
| *Corynebacterium rouxii* FRC0190 T | Badell et al. 2020 | DSM 110354; CIP 111752 | *Corynebacterium rouxii* | 2451 019 | 53.2 | 2365 | | PRJNA224116 | SAMEA5992727 | GCF_902702935 | |
| *Corynebacterium pelargi* DSM 46737 | Kämpfer et al. 2015 | 136/3; LMG 28174; CCM 8517; CIP 110778 | *Corynebacterium pelargi* | 2370 060 | 58.2 | 2169 | Gp0442046 | PRJNA224116 | SAMN06041739 | GCA_004114895 | |
| *Corynebacterium pseudotuberculo sis* ATCC 19410 | (Buchanan 1911) Eberson 1918 emend. Nouioui et al. 2018 | CCUG 2806; DSM 20689; ATCC 19410; NCTC 3450; JCM 9389; IFO 15363; NBRC 15363; CIP 102968 | *Bacillus pseudotuberculo sis*; *Corynebacterium pseudotuberculo sis* | 2337 763 | 52.2 | 2146 | Gp0223239 | PRJNA382169 | SAMN06701041 | GCA_002155265 | |
| *Corynebacterium pseudotuberculo sis* DSM 20689 | (Buchanan 1911) Eberson 1918 emend. Nouioui et al. 2018 | CCUG 2806; DSM 20689; ATCC 19410; NCTC 3450; JCM 9389; IFO 15363; NBRC 15363; CIP 102968 | *Bacillus pseudotuberculo sis*; *Corynebacterium pseudotuberculo sis* | 2338 546 | 52.2 | 2084 | Gp0220522 | PRJNA442833 | SAMN08778220 | GCA_003634885 | 2756170169 |
| *Corynebacterium ulcerans* NCTC 7910 | (ex Gilbert and Stewart 1927) Riegel et al. 1995 | CCUG 2708; DSM 46325; ATCC 51799; JCM 10387; CIP 106504 | *Corynebacterium ulcerans* | 2453 761 | 53.3 | 2178 | Gp0262745 | PRJEB6403 | SAMEA4504038 | GCA_900187135 | |

| Strain | Authority | Other deposits | Synonyms | Base pairs | Percent G+C | No. proteins | Goldstamp | Bioproject accession | Biosample accession | Assembly accession | IMG OID |
| --- | --- | --- | --- | --- | --- | --- | --- | --- | --- | --- | --- |
| *Corynebacterium humireducens* DSM 45392 | Wu et al. 2011 emend. Nouioui et al. 2018 | CGMCC 2452; NBRC 106098; MFC-5 | *Corynebacterium humireducens* | 2681 312 | 68.6 | 2545 | Gp0023681 | PRJNA172965 | SAMN03283197 | GCA_000819445 | |
| *Corynebacterium kutscheri* DSM 20755 | (Migula 1900) Bergey et al. 1925 emend. Nouioui et al. 2018 | CCUG 27535; ATCC 15677; NCTC 11138; JCM 9385; IFO 15288; NBRC 15288; CIP 103423 | *Bacterium kutscheri*; *Corynebacterium kutscheri* | 2354 065 | 46.5 | 2047 | Gp0110293 | PRJNA276037 | SAMN03365283 | GCA_000980835 | |
| *Corynebacterium mustelae* DSM 45274 | Funke et al. 2010 emend. Nouioui et al. 2018 | 3105; CCUG 57279 | *Corynebacterium mustelae* | 3474 226 | 52.6 | 3110 | Gp0114696 | PRJNA282348 | SAMN03568800 | GCA_001020985 | |
| *Corynebacterium diphtheriae* NCTC 11397 | (Kruse 1886) Lehmann and Neumann 1896 emend. Nouioui et al. 2018 | DSM 44123; ATCC 27010; CIP 100721 | *Bacillus diphtheriae*; *Corynebacterium diphtheriae*; *Corynebacterium diphtheriae* subsp. *diphtheriae* | 2463 666 | 53.5 | 2337 | Gp0132011 | PRJEB6403 | SAMEA2517360 | GCA_001457455 | |
| 02-13 | | | | 2518 913 | 53.3 | 2320 | | | | | |
| 03-8664 | | | | 2329 785 | 53.6 | 2342 | | | | | |
| 04-13 | | | | 2548 286 | 54.4 | 2528 | | | | | |
| 04-15 | | | | 2446 713 | 53.3 | 2202 | | | | | |
| 04-3911 | | | | 2492 680 | 53.3 | 2289 | | | | | |
| 04-7514 | | | | 2497 845 | 53.5 | 2375 | | | | | |
| 05-13 | | | | 2551 141 | 54.4 | 2531 | | | | | |
| 06-16 | | | | 2516 702 | 53.4 | 2330 | | | | | |

| Strain | Authority | Other deposits | Synonyms | Base pairs | Percent G+C | No. proteins | Goldstamp | Bioproject accession | Biosample accession | Assembly accession | IMG OID |
|---|---|---|---|---|---|---|---|---|---|---|---|
| 06-19 | | | | 2446 966 | 53.3 | 2205 | | | | | |
| 0102 | | | | 2579 188 | 53.4 | 2349 | | | | | |
| 0211 | | | | 2579 078 | 53.4 | 2350 | | | | | |
| 809 | | | | 2502 095 | 53.3 | 2250 | | | | | |
| 2590 | | | | 2501 366 | 53.3 | 2279 | | | | | |
| 4940 | | | | 2419 371 | 53.3 | 2188 | | | | | |
| 05146 | | | | 2466 435 | 53.3 | 2208 | | | | | |
| 131001 | | | | 2483 321 | 53.3 | 2237 | | | | | |
| 131002 | | | | 2434 569 | 53.4 | 2185 | | | | | |
| 210931 | | | | 2509 428 | 53.3 | 2292 | | | | | |
| 210932 | | | | 2484 335 | 53.3 | 2235 | | | | | |
| BR-AD22 | | | | 2606 374 | 53.4 | 2406 | | | | | |

#### Results

##### Type-based species and subspecies clustering

The resulting species and subspecies clusters are listed in Table 4, whereas the taxonomic identification of the query strains is found in Table 1. Briefly, the clustering yielded 15 species clusters and the provided query strains were assigned to 3 of these. Moreover, user strains were located in 4 of 16 subspecies clusters.

##### Figure caption genome tree

**Figure 1.** Tree inferred with FastME 2.1.6.1 [6] from GBDP distances calculated from genome sequences. The branch lengths are scaled in terms of GBDP distance formula $d_5$. The numbers above branches are GBDP pseudo-bootstrap support values > 60 % from 100 replications, with an average branch support of 36.5 %. The tree was rooted at the midpoint [7].

##### Figure caption SSU tree

**Figure 2.** Tree inferred with FastME 2.1.6.1 [6] from GBDP distances calculated from 16S rDNA gene sequences. The branch lengths are scaled in terms of GBDP distance formula $d_5$. The numbers above branches are GBDP pseudo-bootstrap support values > 60 % from 100 replications, with an average branch support of 44.4 %. The tree was rooted at the midpoint [7].
